# Exposure route drives SARS-CoV-2 infection patterns in non-human primates

**DOI:** 10.64898/2026.03.03.709384

**Authors:** Celine E. Snedden, Dylan H. Morris, Thomas C. Friedrich, James O. Lloyd-Smith

## Abstract

Public health policy and clinical interventions against infectious diseases rely on understanding the factors that govern the initiation and progression of infections inside hosts. Animal infection experiments are essential tools to study these complex, multi-scale processes under controlled conditions, and they have shown that the dose and route of exposure can influence within-host disease dynamics. However, observed differences are difficult to quantify and to disentangle from confounding factors because small sample sizes limit the statistical power and scientific scope of individual studies. Here, by compiling and analyzing the largest published database of non-human primate challenge experiments (107 studies; 721 animals; 22,183 observations), we quantify how exposure conditions and demographic factors shape within-host SARS-CoV-2 infection kinetics in the respiratory and gastrointestinal tracts. We show that exposure route has stronger effects on kinetics than dose, age, sex, or species, with directly exposed tissues exhibiting distinct spatiotemporal kinetics from non-exposed tissues. We estimate 50% infectious doses for different tissues and show that they vary greatly (from <10^1.2^ up to >10^7.4^ pfu) depending on the exposure route. We find that dose effects on kinetics are also route-and tissue-specific, primarily influencing nasally-inoculated animals. Our results suggest that exposure route drives infection kinetics more strongly than dose in a critical model system for translational medicine, and they demonstrate the untapped potential for meta-analysis to extract additional insights from costly animal experiments.

## Introduction

Epidemiologists and microbiologists have long recognized that the dose and route of pathogen exposure influence disease manifestations, within-host tissue distribution, and shedding profiles of infected hosts^1–13^, but these patterns have not been formally quantified or systematically explored. Existing knowledge stems primarily from animal challenge experiments, where exposure conditions are known and controllable, unlike in natural, community-acquired infections. Non-human primate (NHP) models are especially important for translating experimental insights to clinical and epidemiological contexts, given the genetic, immunologic, and physiological similarity between NHPs and humans^14^. However, ethics, logistics, and costs constrain the sample sizes of NHP experiments, leading to weak statistical power, limited scientific scope, and high risk of confounding factors in individual studies. Differences in the estimated strength or direction of individual effects occur frequently among studies of the same virus (e.g., the relationship between SARS-CoV-2 infection duration and illness severity^15^) and of different viruses (e.g., the relationship between dose and infection AUC for influenza A^16^ and Zika^17^ viruses). Many fundamental questions about infectious diseases are therefore difficult to answer, especially quantitatively. How do exposure dose and route determine the extent and timing of within-host dissemination patterns? Do shedding and severity scale with exposure dose, and is this pattern route-specific? Does individual-level variation in infections arise chiefly from exposure conditions or host characteristics? In order to improve human health while addressing the ethical imperatives^18^ and funder objectives^19^ to reduce animal use, novel analytical methods that can extract more insights from these costly experiments are needed.

Taken together, animal infection experiments have generated an immense amount of data, despite small sample sizes in individual studies. Qualitative reviews synthesize this information for many pathogens^20–22^. Although there are numerous potential benefits of analysing this data jointly using quantitative methods^23^, such meta-analyses are exceptionally rare, largely due to concerns about variable protocols across laboratory groups^24,25^. Specifically, differences in experimental design such as sampling schedules, animal demographics, inoculum volume, and pathogen quantification methods (e.g., RT-qPCR vs. cell culture) are often viewed as confounding the ability to make meaningful comparisons across studies. However, meta-analytic and Bayesian methods that account for study-specific differences have enabled robust advances across scientific disciplines, including in neurobiology^26^, virological diagnostics^27^, and ecology^28^. Parametric survival models^29^ can overcome sample time inconsistencies by treating observations as censored data, while also allowing researchers to estimate the effects of cofactors (e.g., demographics) on event times of interest. Harnessing the latent potential of existing data with these modern statistical approaches could allow us to address standing questions in microbiology, namely those for which individual experiments are underpowered.

Here we report the largest-ever quantitative meta-analysis of within-host infection patterns for a respiratory pathogen based on individual-level data from mammal challenge experiments. By assembling a comprehensive database of SARS-CoV-2 challenge experiments in non-human primates (NHP; 107 studies; 721 animals; 22,183 observations) and analyzing it with a Bayesian survival model, we characterize the effects of exposure dose, exposure route, and demographic factors (age, sex, species) on the probability, onset, peak, and conclusion of detectable infection across the respiratory and gastrointestinal tracts. We identify exposure route as the primary driver of variation in within-host SARS-CoV-2 infection patterns. We show that the effects of dose are route-and tissue-specific and that aerosol inoculation results in significantly different spatiotemporal kinetics than all other tested exposure routes. Our results shed new light on SARS-CoV-2 infection kinetics in a critical model system for translational medicine research^14^.

## Results

### Data and model overview

Following a comprehensive literature search, we constructed a database of viral titer measurements from 107 articles^7,30–135^ that challenged NHPs with ancestral SARS-CoV-2 strains (i.e., ones that circulated prior to the emergence of the Alpha variant; **Fig. S1**; **Table S1**). From these studies, we extracted 22,183 observations from 721 animals across the respiratory, gastrointestinal, and other systems (**Fig. S2**). We used this large database to construct the dataset for the analyses herein (**Fig. 1a**; 103 studies had data available for inclusion^7,30–131;^ **Fig. S1**), which contains many exposure doses (10^1.2^-10^7.4^ pfu), three NHP species (rhesus macaque, cynomolgus macaque, African green monkey), both sexes, and three age classes (juvenile, adult, geriatric). Studies in this dataset used many different exposure routes, which we grouped into five categories for modeling purposes (**Table S2**). In all analyses, we report results for one representative route from each category, namely intranasal (IN), intratracheal (IT), combined 50% intranasal and 50% intratracheal (IN+IT), aerosol (AE), and intragastric (IG) exposures (**Fig. 1a**; note that, for visual clarity, data from all similar exposure routes are included in each labelled category; e.g., intrabronchial is included with IT). Multiple methods of viral quantification were reported in the constituent studies, including RT-qPCR (targeting genomic RNA [gRNA]; subgenomic RNA [sgRNA]; or gRNA and sgRNA simultaneously [total RNA]) and culture assays (endpoint dilution [TCID50]; plaque assay [pfu]). Our model utilizes all of this data, but we primarily report results in pfu. Our analyses also focus on the nose, throat, trachea, lung, upper gastrointestinal tract, and lower gastrointestinal tract (**Fig. 1b**). We include data from swabs throughout infection in addition to invasive samples at necropsy for all tissues except the upper GI, which was only sampled invasively (see **Table S3** for all sample types available for each tissue). Sample size coverage across these study dimensions is generally good, though fewer geriatric animals were available, nearly all lower-dose (<10^4^ pfu) inoculations used aerosol exposure, and few studies used intragastric exposure (**Fig. S3**).

**Figure 1.**
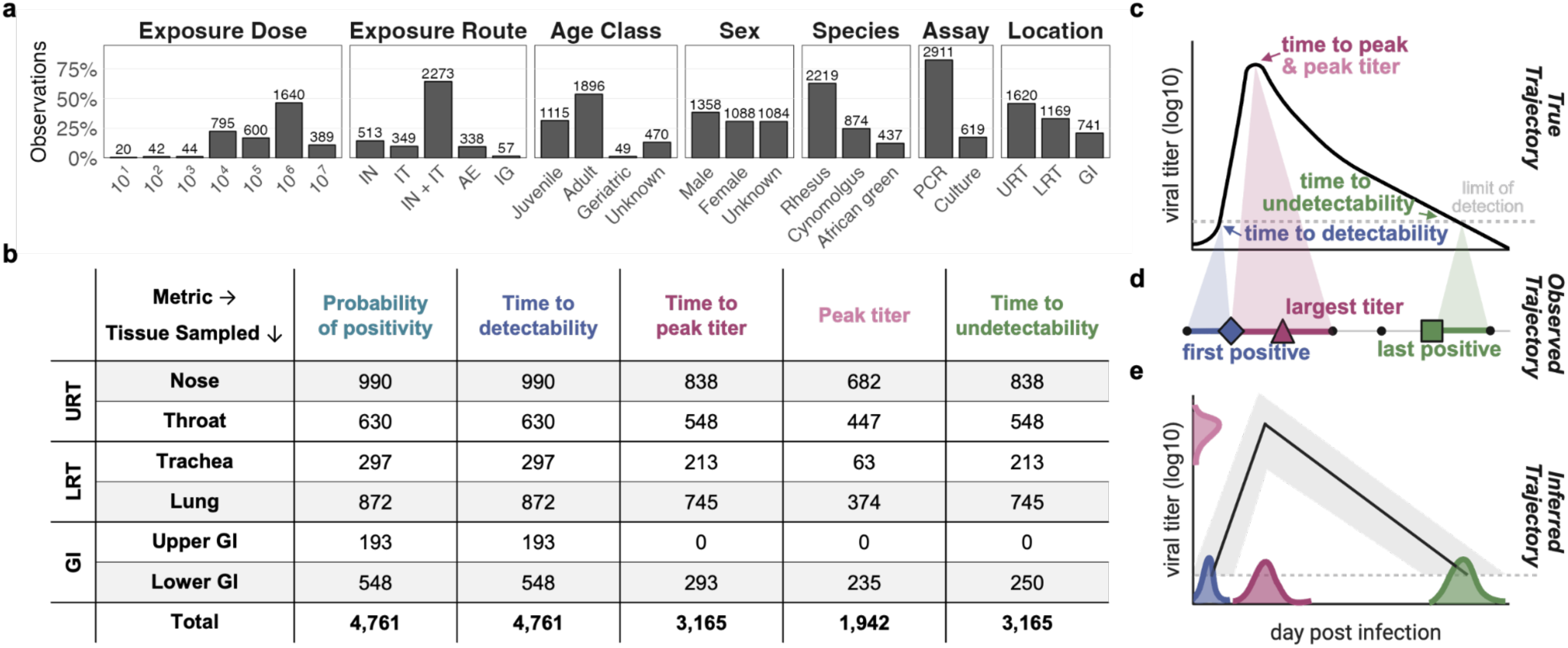
Data summary and diagram of modeling approach. **a**, Data distribution across key cofactors for the dataset analyzed with our Bayesian survival model. Bar heights give the percentage of all available samples. Annotated numbers give the total number of observations, where one observation corresponds with one individual that was sampled in one tissue location by one assay type, but may encompass multiple time-points. One individual may yield multiple observations if they were sampled in multiple locations or if their sample was run with multiple assay types. Exposure dose is given in pfu. Exposure routes are categorized into five broad types (**Table S2**). We chose a representative route for each category, which we used to label this figure: IN, intranasal; IT, intratracheal; IN+IT, combined intranasal and intratracheal; AE, aerosol; IG, intragastric. Note that data from other similar routes are included in these categories for visual clarity in this figure (e.g., intrabronchial is included with IT; **Table S2**). **b**, Sample sizes across tissues and metrics. Tissues with shaded rows are the primary focus for the analyses in this study. URT: upper respiratory tract; LRT: lower respiratory tract; GI; gastrointestinal tract. **c**, A hypothetical infection trajectory with labels for the key metrics (colored as in panel b; probability of positivity not pictured). **d**, Hypothetical observed quantities for the time series in c, which includes observed event times (first positive, diamond; largest titer, triangle; last positive, square) and other sampling days without an observed event (black dots). Observed event times are censored. Bold colored lines indicate the possible true times given the censored observations. Shaded regions point to the latent (unobserved) true time. **e**, Hypothetical model predictions for the observed trajectory in d. Densities are estimated probabilities for a given time or titer (colored as in c). The black line gives the median inferred trajectory, with the grey region indicating the 90% prediction interval.

To model within-host kinetics, we characterized infection in each tissue by: (i) the probability of ever testing positive for virus, (ii) the time to virus detectability, (iii) the time to peak viral titer, (iv) the peak titer itself, and (v) the time to virus undetectability (**Fig. 1c**). From these metrics, we also calculated the duration of infection and the area under the infection curve (AUC). Due to sampling constraints, these events are not directly observable. However, for any given individual in our database, we bound each metric within an interval based on their sampling dates, first observed positive, largest observed titer, and last observed positive (sample sizes in **Fig. 1b**; hypothetical example in **Fig. 1d**; real examples from our dataset in **Extended Data Fig. 1** and **Fig. S4**). This censoring approach allows us to explicitly account for differences in sampling times and frequency across tissues and studies. We used these observations to fit our Bayesian survival model, which estimates the effects of exposure conditions (dose, route), demographic factors (age, sex, species), and methodological variation (assay, and a study-level random effect encompassing all other effects specific to a given article) on overall infection patterns while also inferring true event times and peak titers for each individual (hypothetical example in **Fig. 1e**). Formally, we fit each event as a delay from the previous event (e.g., days between detectability and peak titer), but for interpretability we report all results in days since inoculation.

The model fits the data well for all metrics, generating predictions that are highly concordant with the observed number of positive individuals (**Extended Data Fig. 2a; Fig. S5**) and the bounded event times (**Extended Data Fig. 2b,c,e**; **Figs. S5-7**). Inferred peak titers are highly correlated with the largest observed titers (r=0.98 for all PCR assays, **Extended Data Fig. 2d**; r=0.95 for plaque assay, **Fig. S8**) but are on average larger than the largest measured titers (by 0.40 log10 RNA copies for PCR, **Extended Data Fig. 2d**, inset; by 0.73 log10 pfu for culture, **Fig. S8**). This reflects the reality that it is highly unlikely to sample exactly when an individual peaks, and so the largest observed value reflects an individual’s still-increasing or already-decreasing viral load.

Model predictions were highly variable across articles (**Extended Data Fig. 3**). However, differences were small among those that used similar exposure routes and detection assays (**Fig. S9**), especially when generating predictions for a fixed set of covariates (**Fig. S10**). Parameter estimates for lab effects were also small (**Figs. S11,12**), and so our subsequent analyses report results for a baseline study (i.e., accounting for article effects but not visualizing them). **Tables S4-S8** report all other parameter estimates. All subsequent analyses display inferred distributions on median event times, which can be sampled to obtain individual-level trajectories (**Fig. S13**).

### Exposure route has the largest effect on infection patterns

Infection patterns varied substantially within each tissue when considering model predictions for all possible combinations of exposure route, dose, age, sex, and species that we tested (example median trajectories for all cofactors in the nose in **Fig. 2a**; depiction and quantification of variance for all tissues in **Figs. S14,S15**). Strikingly, most of the variation was attributable to differences in exposure route, which had the largest effect on 35 of the 37 total tissue-level metrics that we tested (indicated by bold cell borders in **Fig. 2b**). For example, the predicted difference in the duration of culture-detectable lung infection was 3.34 days when averaged across all pairwise route comparisons. The mean difference in this metric between age classes was 0.97 days, which is only 29% of route’s effect (indicated by color intensity in **Fig. 2b**; reported numerically in **Fig. S16**).

**Figure 2.**
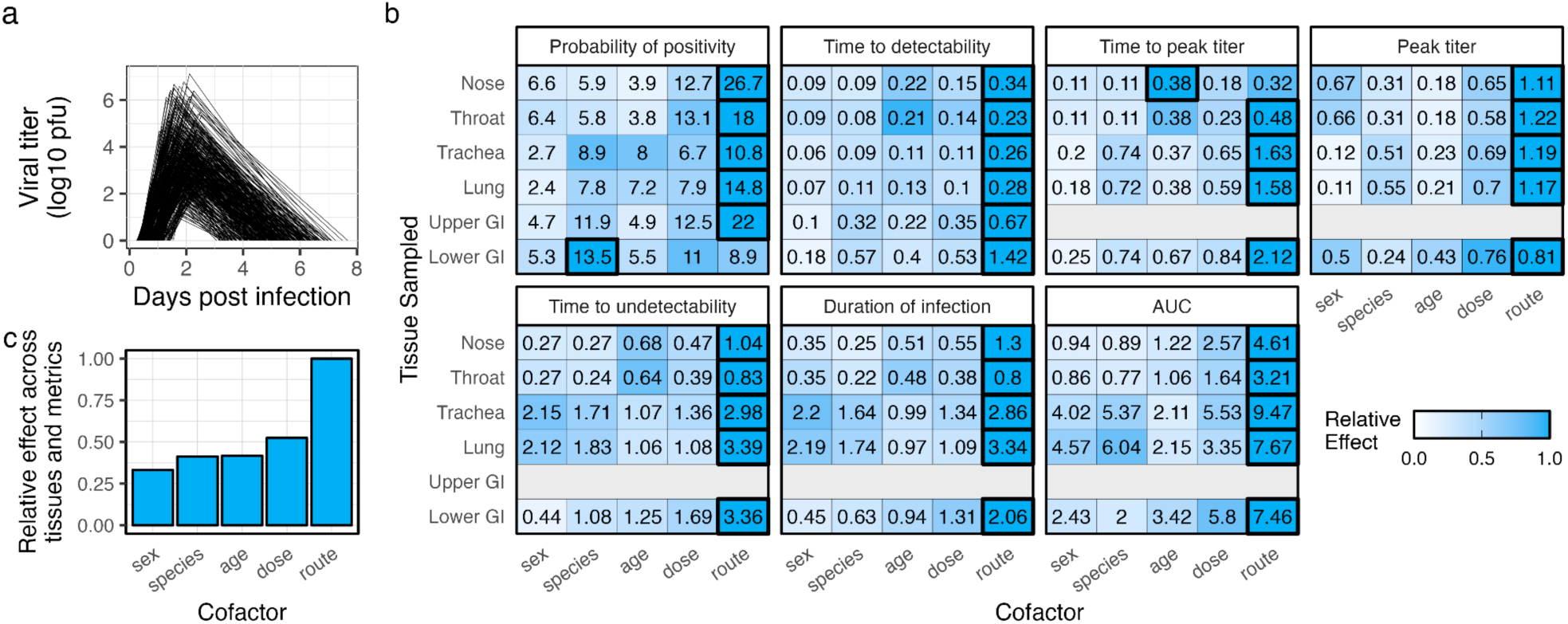
Exposure route is the top determinant of within-host kinetics. Results shown in this figure are based on model predictions for all ages, sexes, species, and exposure routes, but only doses of 10^4^ and 10^7^ pfu, given that most of the data occurs in this range (**Fig. 1a**). All results are for culture detection. **a**, Example median infection trajectories in the nose. Two sampled trajectories are depicted for each possible combination of cofactors (e.g., there two trajectories for a female, adult rhesus macaque exposed intranasally to 10^4^ pfu), for a total of 360 plotted trajectories. **b**, Mean pairwise differences in model predictions for each metric (panel label) and sampled tissue (row label) among the constituent types of each cofactor (column label) when conditioning on all other covariates (e.g., for age class: juvenile vs. adult, adult vs. geriatric, and juvenile vs. geriatric, when holding all other covariates constant; see Methods). Bold, outlined cells indicate the cofactor with the largest mean difference for the given metric and tissue. Color intensities indicate the relative effect, scaled against the largest mean effect for each metric and tissue. The Upper GI was only sampled invasively, so we only estimated the probability of positivity and time to detectability for this tissue. The Lower GI was sampled non-invasively (e.g., via rectal swabs), so we estimated all metrics. Quantitative comparisons in peak titers across tissues should be made cautiously in this and subsequent figures, given they often involve different sample types (e.g., nasal swabs vs. BAL; **Table S3**). **b**, The sum of the relative effects for each cofactor across all metrics and tissues in panel a, which were then scaled against the largest such sum.

Across all tissues, only two metrics were not driven primarily by route. Age was most important for the time to peak titer in the nose (**Fig. 2b**), with geriatric individuals having peak titers that occurred significantly later than for juveniles or adults (time series for aerosol-exposure in **Extended Data Fig. 4** and averaged across routes in **Fig. S18**; significances in **Fig. S19**). Species had the strongest effect on lower GI positivity (**Fig. 2b**) because African green monkeys had significantly higher probabilities of testing lower GI positive than rhesus or cynomolgus macaques (**Extended Data Fig. 4; Figs. S18,19**).

The data available for African green monkeys is not clearly skewed towards cofactors with higher probabilities of GI positivity (**Fig. S3**), suggesting this may reflect true species variability. Other differences among demographic groups were evident and significant (**Fig. S19**), including that males and African green monkeys had the highest lung AUC values, suggesting they may generally experience more severe infections.

When considering all metrics and tissues jointly, exposure route was most influential by a large measure (calculated using the sum of all relative effects; **Fig. 2c**). Although dose never had the largest effect for any individual metric (**Fig. 2b**), it ranked second in aggregate importance across metrics and tissues (**Fig. 2c**). The finding that route and dose ranked highest overall was robust to calculations based on different observational assays (total RNA vs. culture), to choice of summary statistic, and to alternative dose comparisons (**Fig. S17**). Cofactor variability (**Fig. 2b**) and broader infection patterns (**Fig. S20**) in the throat and trachea largely mirrored the nose and lung, respectively, so subsequent analyses focus chiefly on the nose, lung, and lower GI, which had larger sample sizes (**Fig. 1b**).

### Early infection kinetics differ among exposed and non-exposed tissues

Within-host dissemination and early spatiotemporal infection patterns clearly varied among exposed and non-exposed tissues, where we classified the following tissues as exposed: (i) nose and throat for IN; (ii) trachea for IT; (iii) nose, throat, trachea, and lung for AE; and (iv) GI tissues for IG (see **Methods**). Exposed tissues had higher probabilities of culture positivity (darker colors in **Fig. 3a**), and they became culture-detectable earlier than non-exposed tissues (**Fig. 3b**; many pairwise comparisons were significant, **Figs. S21,22**; median predictions and 90% intervals for all metrics in **Tables S9-S14**). Notably, IT exposure was the only route that resulted in substantially higher probabilities of lung positivity than nasal positivity (**Fig. 3a**), with a clear pattern of infection spreading from the LRT to the URT and then the GI (**Fig. 3b**). This contrasts starkly with IN exposure, which resulted in the lowest probability of lung detection among respiratory routes (**Fig. 3a**) and was the only route where nasal positivity preceded lung positivity when both occurred (**Fig. 3b**). The probability of an individual ever having culture-positive lower GI specimens was small for all routes (**Fig. 3a**; few pairwise differences were significant; **Fig. S21**). Culture detectability in the GI always occurred after the respiratory tract, even for IG exposure, though time to detectability in the GI was lowest for IG exposure (**Fig. 3b**). Total RNA assays recapitulated these patterns (**Fig. S23**), though with overall higher probabilities of positivity and slightly earlier times to detectability (**Fig. S24**).

**Figure 3.**
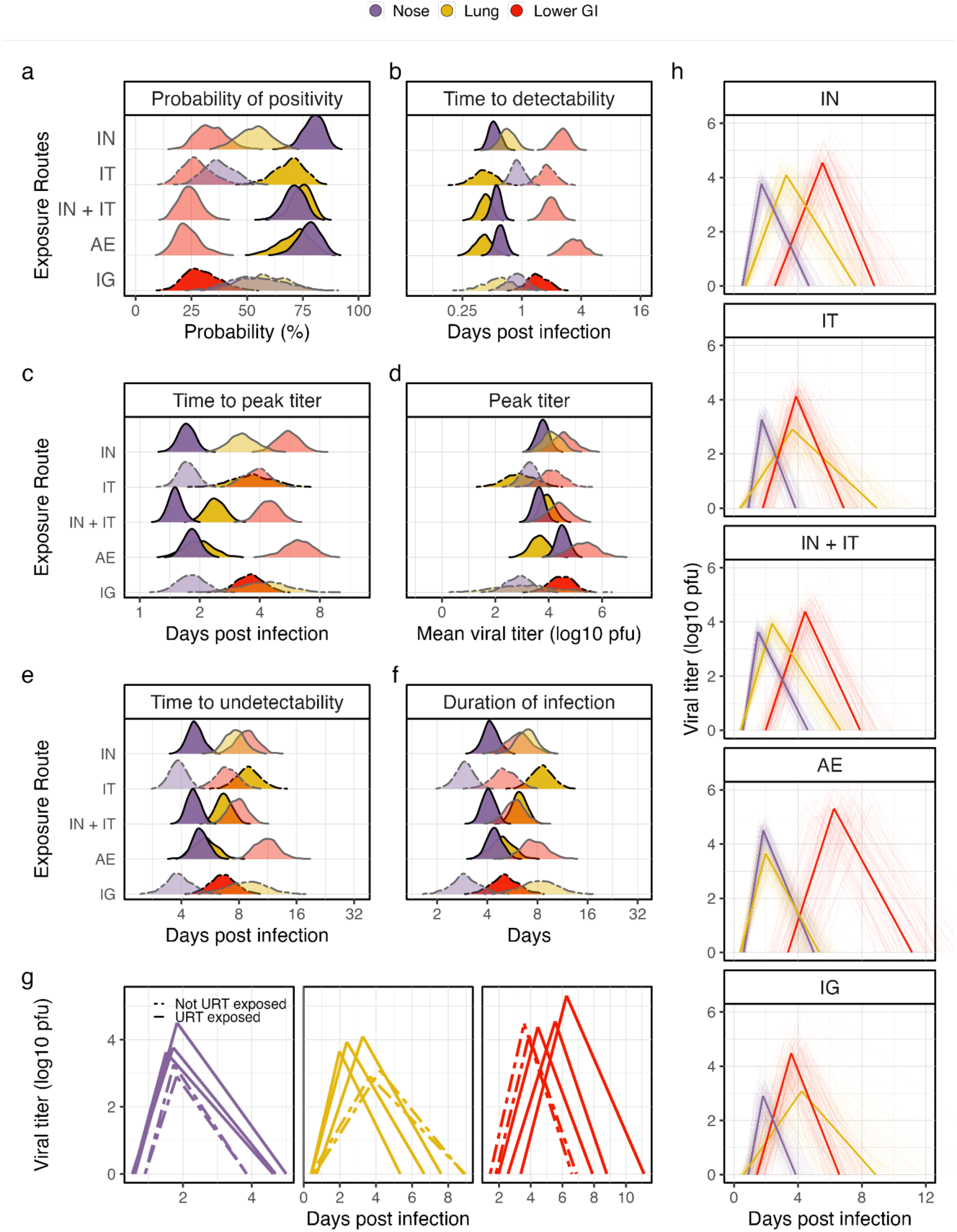
Effects of exposure route on spatiotemporal spread of infectious virus. All results are displayed for viral culture and, for visual clarity, were generated for an adult, female rhesus macaque exposed to 10^4^ pfu by the indicated route. Colors for all panels correspond with the sampled tissue (purple: nose; yellow: lung; red: lower GI). **a-f**, Model predictions across exposure routes for: **a,** the probability of positivity; **b**, the median time to detectability; **c,** the median time to peak titer; **d**, the mean peak titer; **e**, the median time to undetectability; and **f**, the median duration of infection. Solid lines indicate routes that include upper respiratory exposure (IN, IN+IT, AE) while dashed lines are routes that do not (IT, IG). More intense colors correspond with exposed tissues, which we consider to be those directly inoculated as well as those adjacent to inoculated tissues (e.g., the lung for IT exposure, given that fluid can drain immediately to the lung). **g,** Median trajectories for all five routes separated into panels by tissues. Inoculation routes are categorized by whether they include nasal exposure (solid lines; IN, IN+IT, AE) or not (dashed; IT, IG). **h**, Trajectories for each route indicated in the panel label. Thick opaque lines are the median infection trajectories. Thin transparent lines display trajectories based on 100 posterior samples.

### Tissue connectivity structure aligns with physical proximity

To estimate a within-host connectivity structure for SARS-CoV-2, we analyzed model predictions for the probability and timing of RNA detectability following all possible single-tissue inoculations (e.g., nose-only or throat-only exposure). With these predictions, we identified the primary virus outflow destination (black cells with a ‘1’ in **Fig. 4a**; quantities standardized within rows) and virus inflow source (black cells with a ‘1’ in **Fig. 4b**; quantities standardized within columns) for each tissue (**Fig. S25**), which we then translated into a connectivity diagram (**Fig. 4c**). The resulting structure clearly aligns with physical proximity and macroscopic anatomy, with strong evidence of viral spread both up and down the respiratory tract and from the throat to the GI tract. Similar directionality was evident when analyzing the mean inferred times to detectability of all animals in the dataset that were exposed via single-route inoculations (**Fig. S26**). Viral flow from the upper GI to the lung (**Fig. 4a,c**) could reflect dissemination via lymph and blood^136^ but could also be a consequence of the inoculation procedure or sampling protocol. Together, these analyses support the mucociliary escalator, aspiration, and swallowing as crucial mechanisms of within-host viral dissemination.

**Figure 4.**
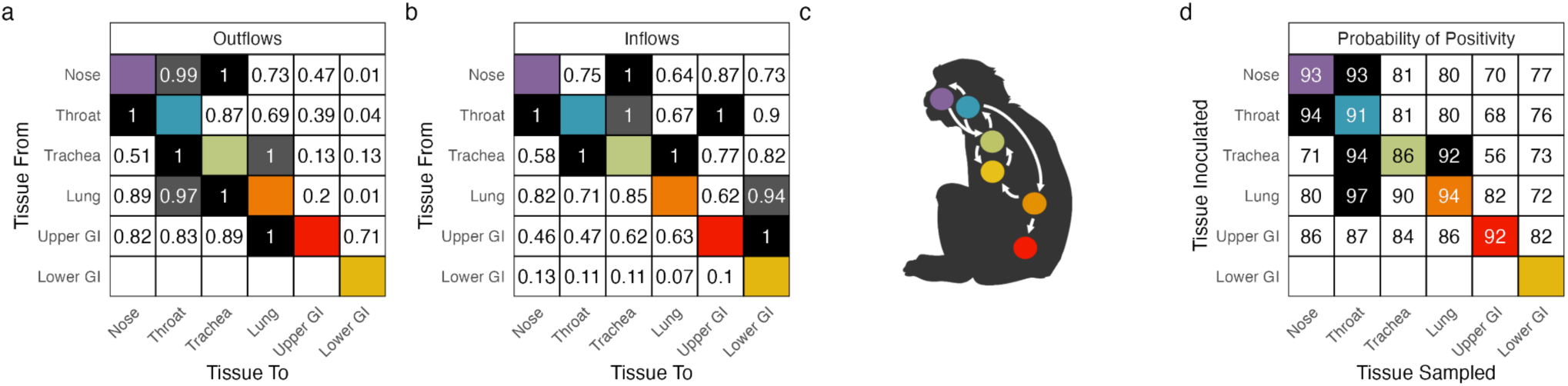
Estimated tissue connectivity structure. All results are based on the order that tissues become detectable after single-tissue inoculations with 10^7^ pfu (see Methods). **a,** Outflows from the y-axis tissue into the x-axis tissues. Black cells with a white annotated ‘1’ indicate that, after inoculation into the y-axis tissue, the x-axis tissue was most frequently the next one to become detectable. All other row-specific values are frequencies relative to that tissue. Diagonal cells are colored based on the x-and y-axis tissue. Tissues with values >0.9 reflect fairly strong connections and are denoted with a dark grey cell with white text. For example, after inoculation into the throat (second row from the top), most of the predictions resulted in the nose testing positive next; the trachea had 87% as many predictions where it tested positive next. **b**, Inflows into the x-axis tissue from the y-axis tissues. Black cells with a white annotated ‘1’ indicate that the y-axis tissue was most likely to test positive before the tissue on the x-axis. All other column-specific values are frequencies relative to that tissue. Cells are colored as in a. For example, the throat is most likely to become detectable before the nose does (leftmost column). **c,** Connectivity structure based on all directional connections with a ‘1’ in panels a and b. Colors distinguish between tissues, as in panel a. The trachea had two highly similar inflow sources (both marked ‘1’); both are included in the diagram. **d**, The probabilities of the x-axis tissue ever testing positive following single-tissue inoculations into the y-axis tissue. Black cells with white text have probabilities above 90%. Cells along the diagonal are colored as in panel a.

Our model predictions also highlight that detectable within-host spread among tissues is not guaranteed during infection, even when a strong dissemination pathway exists between two tissues. Infections initiated in one tissue of the respiratory tract often fail to establish detectable infection in other respiratory tissues, with probabilities that tend to decrease with increasing physical distances (**Fig. 4d**) and likely also reflect sampling methods and tissue competence. This finding is consistent with our route-specific results (**Fig. 3a**) and with the percent of animals in our dataset that tested positive following single-route inoculations (**Fig. S26**).

### The peak and conclusion of infection are sensitive to upper respiratory exposure

Differences in the peak and conclusion of infection were largely driven by whether inoculation included URT exposure (**Fig. 3c-g**; **Fig. S21**). Peak times for culture in the nose were remarkably consistent across routes and typically occurred before peak titers in the lung (**Fig. 3c**). However, the nose had larger peak titers (**Fig. 3d,g**), later times to undetectability (**Fig. 3e,g**), and overall longer infection durations (**Fig. 3f,g**) when it was exposed (IN, IN+IT, AE; solid lines) than when it was not exposed (IT, IG; dashed lines). Some of these effects were reversed in the lung, which often had earlier peak times (**Fig. 3c,g**), larger peak titers (**Fig. 3d,g**), earlier times to undetectability (**Fig. 3e,g**), and shorter infection durations (**Fig. 3f,g**) when the nose was exposed. In the lower GI, URT inoculation resulted in later peak times (**Fig. 3c,g**), similar (though slightly larger) peak titers (**Fig. 3d,g**), later times to undetectability (**Fig. 3e,g**), and longer infection durations (**Fig. 3f,g**). Patterns were similar for total RNA (**Fig. S23**). Interestingly, peak times rarely deviated by more than one day between total RNA and culture, but total RNA remained detectable much longer than viable virus in the respiratory tract, in contrast to the GI where undetectability occurred almost simultaneously (**Fig. S24**).

### Aerosol exposure leads to distinct infection kinetics

When considering the full course of infection across tissues, AE exposure clearly resulted in different spatiotemporal patterns than all other routes (**Fig. 3h**; 7/103 studies included AE exposure). AE exposure was the only route where infection kinetics in the lung clearly matched those in the nose, with almost simultaneous times to detectability, peak titer, and undetectability for viral culture following inoculation with 10^4^ pfu. AE exposure also resulted in higher peak titers in the nose and lower GI, as well as delayed and prolonged GI positivity. These patterns ultimately resulted in shorter infection durations in the lung but longer durations in the nose and lower GI for AE exposure compared to all other routes. Crucially, despite IN+IT exposure being commonly used as a tractable model for AE exposure, the resulting infections differed significantly for many metrics (**Figs. S21, S22**) and their kinetics were visibly distinct (**Fig. 3h**). Ultimately, these discrepancies emphasize that no other exposure route approximates all features of infection patterns following AE inoculation.

### ID50 estimates are strongly dependent on exposure route, tissue sampled, and detection assay

We used our model to estimate the median dose that causes detectable infection in 50% of exposed individuals (i.e., ID50), given the importance of this metric in clinical, epidemiological, and research settings. Strikingly, when using viral culture results to determine infection status, our ID50 estimates varied from <10^1.2^ up to >10^7.4^ pfu, depending on the exposure route and tissue sampled (**Fig. 5a**; values integrate over all demographic groups). Estimates for the dose required to yield culture-detectable infection in the nose ranged from 10^1.2^ pfu for IN exposure up to >10^7.4^ pfu for IT exposure, with AE exposure falling at the lower end of this range (10^2^ pfu). ID50 estimates for the lung were 10^1.2^ pfu for IT exposure but as high as 10^4.1^ pfu for IN exposure. These findings are consistent with our connectivity analyses, which showed that infections initiated in one respiratory tissue can fail to establish infection in another respiratory tissue (**Fig. 4c**) despite the existence of a dissemination pathway (**Fig. 4b**). The lower GI had consistently large ID50 estimates for culture positivity, ranging from 10^6.5^ to >10^7.4^ pfu. Broadly, ID50 values were much lower in exposed tissues than non-exposed tissues (dark vs. light cells in **Fig. 5a**).

**Figure 5.**
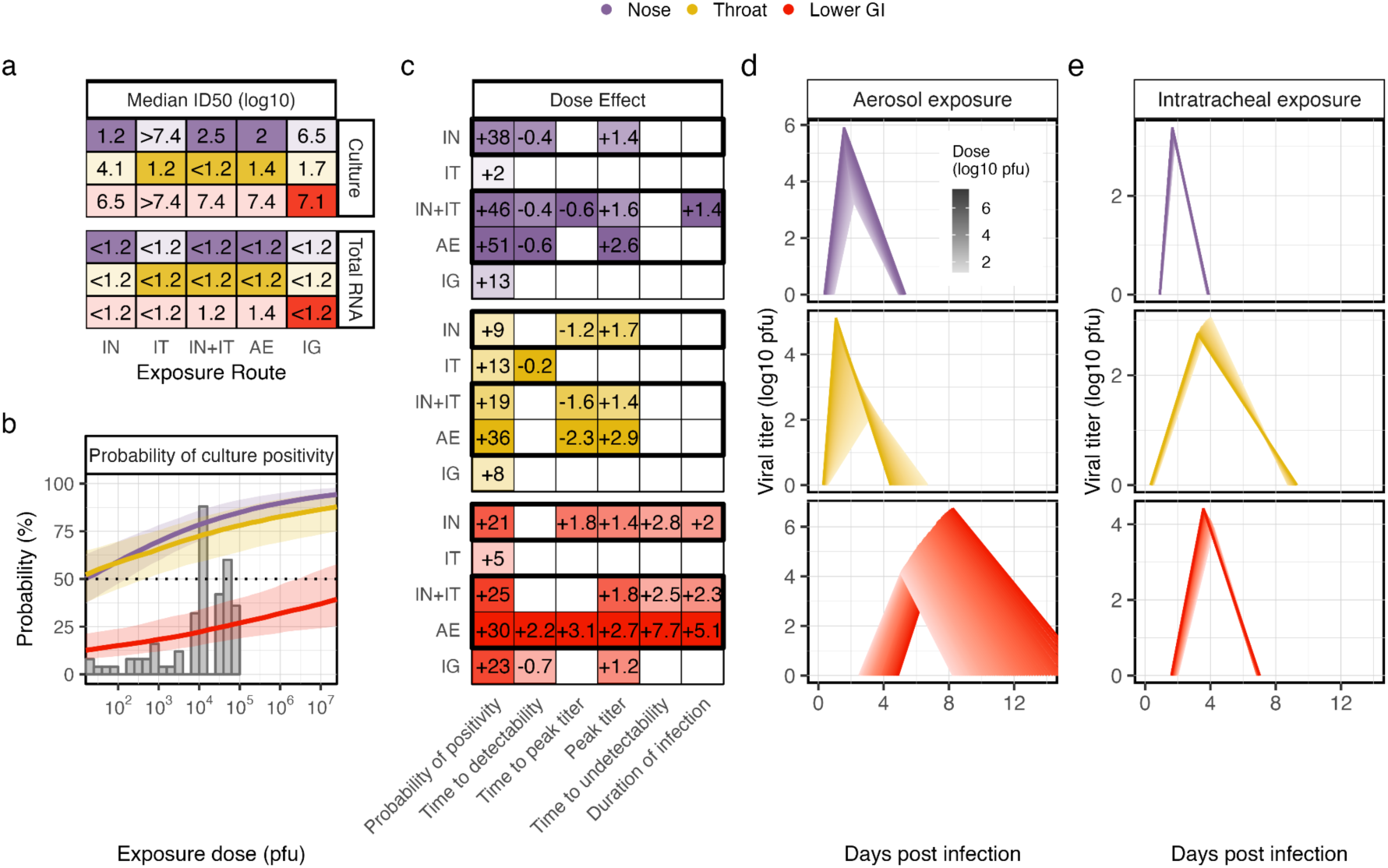
Effects of exposure dose on spatiotemporal spread of infectious virus. Colors in all panels correspond with the sampled tissue, as in the top legend (purple: nose; yellow: lung; red: lower GI). **a**, Median ID50 estimates for each route and for both total RNA and culture assays. Rows and colors correspond with different tissues. Darker shaded cells indicate exposed tissues. **b**, The probability of an individual ever testing culture positive across exposure doses for a female, adult, AE-exposed rhesus macaque. Thick lines are median predictions, and shaded regions are the 90% prediction interval. Overlaid histograms give the distribution of doses available for model fitting from an AE-inoculated individual for all assays. **c,** Effects of dose for each combination of metric (columns), route (rows), and tissue (panels). Annotated text is the mean difference in predictions between the maximum (10^7.4^ pfu) and minimum doses in our database (10^1.2^ pfu) when the difference is significant. White cells without text indicate the difference between doses is not significant. Units vary by metric (probability: percent; event time: days; peak titer: log10 pfu). Color intensity increases with relative effects, which are standardized within each column and panel. Dark boxes enclose routes with upper respiratory exposure. **d,** Median trajectories in each tissue for a female, adult, rhesus macaque exposed via aerosol with doses ranging from 10^1.2^ to 10^7.4^ pfu. Darker colors correspond with larger doses. **e**, As in panel d, except for intratracheal exposure. Median predictions and 90% prediction intervals for all metrics are reported quantitatively in **Tables S9-S15**.

These stark differences in route-specific ID50 values were hidden when instead determining infection status based on RT-qPCR positivity. Our ID50 estimates based on PCR were ≤10^1.2^ pfu for nearly all routes and tissues, and they were always smaller than the tissue-and route-matched estimates based on culture positivity (**Fig. 5a**). These results suggest that viral RNA can disseminate widely, even at doses that are smaller than the levels needed to establish culture-measurable replication in a given tissue.

### Larger doses primarily impact infection kinetics when administered to the URT

Exposure dose affected infection patterns in many ways, although the direction, strength, and significance of these effects often varied across exposure routes and tissues. As expected, larger doses led to significantly higher probabilities of detectable infection across all tissues and routes (effect direction, strength, and significance for all metrics in **Fig. 5c**; dose-specific predictions for aerosol exposure in **Fig. 5b**; predictions for all other routes in **Fig. S27**).

Dose effects on temporal infection kinetics often differed between routes that did or did not include URT exposure. For routes with URT exposure (IN, IN+IT, AE; dark outline in **Fig. 5c**), larger doses significantly: (i) decreased the time to detectability in the nose but had few discernible effects on this metric in the lung or lower GI, (ii) decreased the time to peak titer in the lung and increased them in the lower GI or had no effect, (iii) increased peak titers in all tissues, (iv) increased time to undetectability in the lower GI but not in the respiratory tract, and (v) increased infection duration in the lower GI (**Fig. 5c**; for AE exposure, see trajectories in **Fig. 5d** and individual effects in **Extended Data Fig. 5**; results for other routes in **Figs. S27,S28**). These effects were largest for AE exposure (**Fig. 5c**), which also showed evidence of shorter infection durations in the lung at higher doses (**Fig. 5d**). For routes without URT exposure (IT, IG), there were few measurable dose effects (**Fig. 5c**; for IT exposure, see trajectories in **Fig. 5e**). Larger doses decreased the time to detectability in adjacently-exposed tissues (e.g., lung for IT, lower GI for IG), and they increased peak titers in the lower GI for IG exposure, but they otherwise had no significant effects (**Fig. 5c**). All patterns were similar for total RNA (**Figs. S29-31**). Overall, dose had the biggest impact on the kinetics of individuals exposed in the URT (**Fig. 5c**), perhaps due to the centrality of the throat in the tissue connectivity structure (**Fig. 4c**).

### Clinical profiles vary across exposure conditions

Finally, we computed infection AUC values in the nose, GI, and lung as our best available proxies for respiratory shedding, GI shedding, and disease severity, respectively (median values in **Fig. 6a,b**; variability in **Fig. S32**). Once again, AE exposure had a different profile, with significantly higher nose and GI shedding than all other routes and significantly lower lung severity than IN and IN+IT exposures (**Fig. 6a,b**; significances in **Fig. S21**). Crucially, IN+IT and AE exposures differed significantly for both shedding types and for severity (**Fig. S21**), and IN+IT was more similar to IN exposure than to AE. IT and IG exposures resulted in nearly identical profiles (**Fig. 6a,b**), given they only differed significantly for three metrics across all tissues (**Fig. S22**). Larger doses significantly increased respiratory and GI shedding for IN, IN+IT, and AE exposure (**Fig. 6c**), with AE exposure being most sensitive (**Fig. 6a**). There is some evidence that dose also increases severity for IN, IN+IT, and AE exposures (**Fig. 6b**), though these differences were not statistically significant (**Fig. 6c**).

**Figure 6.**
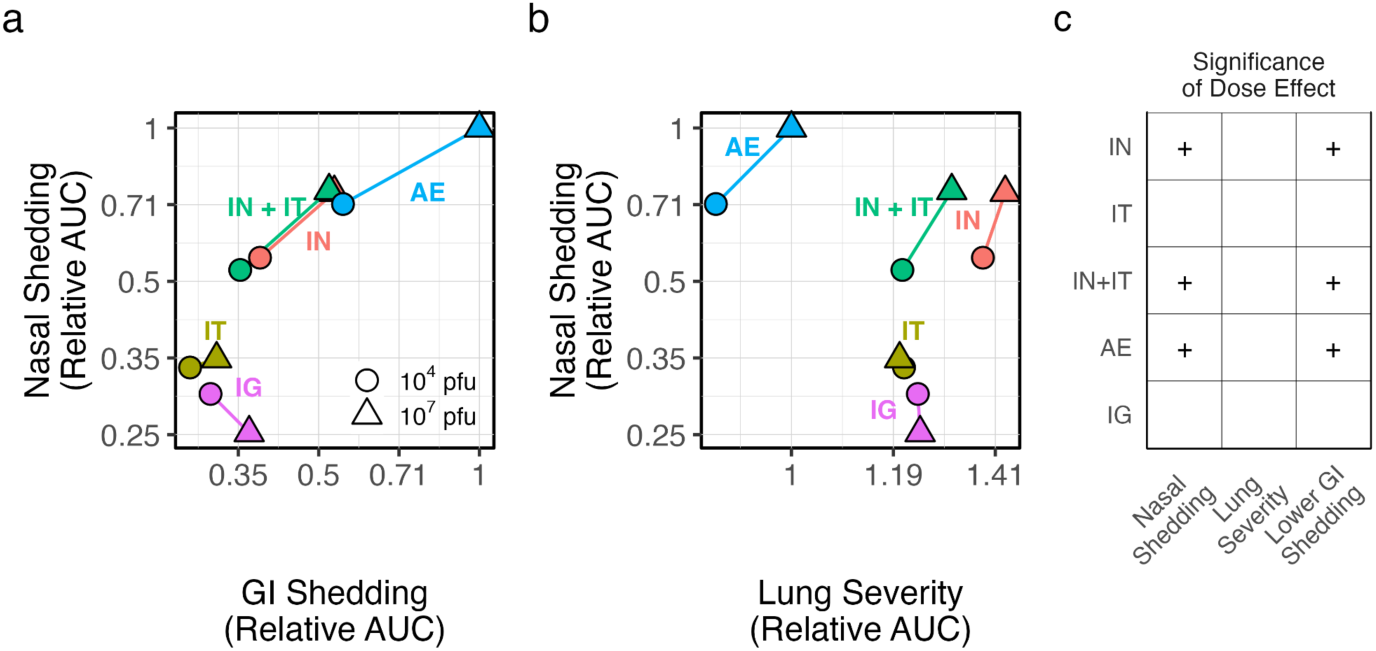
Shedding and severity profiles across exposure conditions. Panels in the top row display relationships between median: **a**, nasal and GI shedding; and **b**, nasal shedding and lung severity. Color corresponds with exposure route, which is also indicated by adjacent, color-matched text labels. Shapes distinguish between doses (circles: 10^4^ pfu; triangles: 10^7^ pfu). Lines connect the points for a given exposure route. Results were first calculated as the median culture AUC value from the infection trajectories of an adult, female, rhesus macaque exposed by the indicated route and dose. All values were then expressed relative to the median AUC value for aerosol exposure with 10^7^ pfu. **d,** Significance of the difference in AUC values between 10^4^ and 10^7^ pfu across all demographic covariates. “+” indicates that AUC increases significantly with dose. Empty cells indicate the difference is insignificant. Shedding and severity profiles across demographic factors are available in **Fig. S19**. Median AUC predictions and their 90% prediction intervals when integrating across demographic factors can be found in **Table S15**.

## Discussion

Historically, small sample sizes have limited the quantitative insights obtained from animal infection experiments, especially given long-standing concerns about combining data across studies conducted with different protocols. Here, by completing the most comprehensive quantitative meta-analysis of mammal challenge experiments to date, we demonstrate that multi-protocol meta-analyses are feasible and powerful, with the potential to yield otherwise unattainable insights. We show that exposure route drives within-host SARS-CoV-2 kinetics in NHPs more strongly than exposure dose, and that the effect of larger doses is evident only for exposures to the upper respiratory tract. No prior study has identified these patterns because no individual experiment has spanned the necessary range of exposure conditions. Concordances between some of our other results and existing work (e.g., similar estimated ID50 values by multiple benchmark criteria^137^) increases confidence in these novel findings, further emphasizing that meta-analyses in virology need not be limited to experiments from a single lab^24,138,139^ or resource-intensive, prospective collaborations across multiple groups^140^. Multi-protocol meta-analyses like ours offer a promising platform to answer fundamental questions in microbiology and pathogenesis, all while supporting objectives to reduce animal use in biological research^18,19^.

Public health and research practices are heavily influenced by observations that larger exposure doses increase infection probability and in some cases disease severity^8,13,141–147^. Our work challenges this focus on dose in the literature: we present the first quantitative evidence that exposure route modulates infection kinetics more strongly than dose, for early SARS-CoV-2 strains in immunologically-naive NHPs. This result is concordant with evidence in various pathogen-host systems that infection probability, spatiotemporal kinetics, and pathogenesis are often highly route-specific^3–6,10,12,13,34,148,149^, while dose effects on kinetics can be small^11,146,150^. The underappreciation of route effects relative to dose may be because direct comparisons are not possible without multi-protocol meta-analysis, given that single studies rarely include multiple exposure routes alongside multiple doses, especially with the factorial design needed to assess multiple drivers. Dose effects may be stronger in other contexts, like those where pre-existing immunity alters virus access, infectivity, or replication in permissive tissues, or at lower dose ranges than are used in NHP experiments. We also found that demographic factors had fewer and weaker effects on kinetics than route, though we did find evidence of higher lung severity for males, echoing findings for male COVID-19 patients^151^. Ultimately, our work emphasizes that route effects warrant further attention in both basic and applied studies across diverse pathogen-host systems.

Our model inferred that SARS-CoV-2 infection kinetics following aerosol inhalation differ significantly from all other routes, including combined intranasal/intratracheal inoculation, despite both routes exposing both upper and lower respiratory tissues. This discrepancy was not apparent at the smaller sample size available in a study from our database that considered both routes^35^, but differences in infection kinetics are common between inoculation methods that are more experimentally tractable and those that better reflect natural exposures, including for mosquito-borne^11,12^ and respiratory^149^ viruses. Notably, all articles that reported infectious respiratory particle (IRP) sizes in our database^7,34,49,66^ used median aerodynamic diameters below 5µm. Given the recent emphasis on transmission of respiratory pathogens through the air^152^ including the physical and virological effects of IRP sizes^153,154^, more research on infection kinetics following aerosol exposure is needed. Future experiments should prioritize low-dose aerosol exposures at different IRP sizes. Given those data, our modeling framework would enable us to assess when (if ever) the administration of liquid inoculum results in similar kinetics to aerosol inhalation. In the meantime, extreme caution is warranted when extrapolating insights from any other inoculation procedure to aerosol exposure. This includes transmission experiments, where differences in infection kinetics between donor and sentinel animals^155^ could be explained by differences in the route of exposure in addition to differences in dose.

ID50 estimates generated from animal and human challenge experiments often vary by exposure route^10,149,153^, but our results demonstrate that they also depend heavily on the tissue used to classify infection status. For example, for intranasal inoculation and culture-based detection, there was a 794-fold difference in the estimated ID50 in the nose versus the lung (10^1.2^ and 10^4.1^, respectively). In general, ID50 estimates were smallest in tissues intentionally exposed during inoculation (e.g., the nose for intranasal exposure). Our ID50 estimates based on PCR detection, however, never exceeded 10^1.4^ pfu for any tissue or exposure route, suggesting that sensitivity of PCR can obscure meaningful patterns in tissue permissivity or that culture success depends more on sample type and material (e.g., respiratory swab vs. stool samples). Our findings emphasize that infection outcomes vary beyond infected/non-infected classifications, and that researchers and practitioners should view ID50 not as a unitary measure but as a function of exposure route and target tissue.

Greater focus on tissue-specific ID50 values for locations that drive between-host transmission (e.g., the nose^156–160^) or disease severity (e.g., the lung^6,161^) could benefit epidemiological and clinical risk assessments for future emerging pathogens. As preliminary proof of principle, we compared evidence related to intranasal exposure with three human coronaviruses. Our ID50 estimates based on PCR-detectable nose infection following intranasal exposure with SARS-CoV-2 were smaller (<10^1.2^ pfu) than comparable estimates for SARS-CoV in NHPs (10^3.4^ pfu^162^), but a similar intranasal dose of hCoV-229E resulted in 50% of participants developing respiratory illness in a human challenge trial (10^1.1^ TCID50^163^). Notably, the two endemic pathogens (SARS-CoV-2, hCoV-229E) have up to a 200-fold lower ID50 value than SARS-CoV, which is concordant with their greater epidemiological success in humans. Tissue-specific ID50 values may also serve as a useful quantitative metric to investigate tissue tropism, given that they reflect the outcome of many biological processes that govern both tissue permissivity and accessibility. In our study, ID50 estimates based on culture-detectable lung infection averaged across routes were smaller (<10^1.9^ pfu) than those based on culture-detectable nose (>10^3.9^ pfu) or lower GI (>10^7.2^ pfu) infection, in accordance with the lung tropism of early SARS-CoV-2 strains^164^. Applying our modeling framework to data from tissues like the heart, liver, and brain could illuminate the experimental and clinical contexts where extrapulmonary sampling or monitoring should be prioritized.

Our results are concordant with and offer further insight into observations from human SARS-CoV-2 infections. Direct comparisons are often difficult given that exposure time, route, and dose are unknown for naturally-acquired infections, but human challenge trials and prospective sampling studies create valuable opportunities to compare NHP and human findings. Our ID50 estimate based on PCR-detectable URT infection following IN exposure (<10^1.2^ pfu) is nearly identical to a dose that resulted in PCR-detectable URT infection of 53% of participants in the SARS-CoV-2 human challenge trial by the same route (≈10^0.69^ pfu^165^). The challenge participants were not in high-risk demographic groups and did not display evidence of pulmonary disease; this is consistent with our estimation that ID50 values based on culture positivity in the lungs of otherwise healthy NHPs (10^4.1^ pfu) are substantially higher than the dose administered (≈10^0.69^ pfu). When monitoring the nose via PCR, our model predictions for infection event times for low-dose aerosol-exposed macaques (10^1^-10^2^ pfu; all ages and sexes) differed by less than 1 day from analogous estimates from a human prospective sampling study^166^, namely for proliferation time (i.e., time between detection and peak titers; 2.3 vs. 3.2 days), clearance time (9.3 vs. 8.5 days), and overall infection duration (11.6 vs. 11.7 days). The human challenge trial^165^ observed remarkable consistency in participants’ infection trajectories following exposure by a single route to a single dose. Taken together with our finding that kinetics in NHPs depend strongly on exposure conditions, this suggests that some of the kinetic heterogeneity observed in naturally-acquired human infections^167^ may arise from variation in the route or dose of exposure. Broadly, the similarities in SARS-CoV-2 kinetics observed in our results and in humans reinforces that NHPs are a powerful animal model for translational research in this system.

Prior work using genetic methods, fluorescent labelling, and barcoded viruses has demonstrated that within-host virus population dynamics are spatially structured and that within-host virus dissemination ranges from restricted to efficient (depending on the virus and host)^156,158,168,169^. Our work corroborates these findings using a novel quantitative framework that does not require special technology or dedicated experiments. As is evident in many pathogen-host systems^170–172^ but has been difficult to quantify systematically, infections in our analyses began in exposed tissues before spreading to physically proximate locations. Yet, despite all tissues in our study being considered permissive to SARS-CoV-2 infection^173,174^, system-wide spread was not guaranteed, even after inoculation with high doses (10^7^ pfu) and even when a highly-connected neighboring tissue was infected. Various processes could explain individual-level heterogeneity in virus spread, including immune responses, the state of physical barriers (e.g., respiratory mucosa), or stochastic effects. Ecologists study similar patterns of limited population spread at landscape scales, which they refer to as dispersal limitation^175^. Theories and modeling techniques developed in invasion ecology may help identify the processes governing spatial within-host infection patterns^176^, especially when employed alongside modern techniques of labelling viruses.

Our findings add nuance to existing theories that larger doses increase disease severity. We used infection AUC values in the lung as our best available proxy for severity, given that prior work has linked severity or pathology with higher titers^6,177^ and delayed clearance^178^, especially in the lower respiratory tract of respiratory viruses^6,161,177^. We found evidence that severity tends to increase with dose (though not significantly) for routes that include upper respiratory exposure (e.g., intranasal, aerosol) but found minimal effect for other routes (e.g., intratracheal). While many SARS-CoV-2 studies have demonstrated positive dose-severity relationships using various metrics^8,146,147,153^ (e.g., weight loss, survival), no prior work to our knowledge has noted this route-dependent effect, possibly because experiments that evaluate different doses almost always inoculate the upper respiratory tract. We also found evidence that aerosol exposure results in milder pulmonary disease than intranasal or combined intranasal/intratracheal exposures for a given nominal dose. These findings partially align with existing work: they agree with the worse lung radiographic scores^34^ but not the similar clinical metrics, bronchial brush viral loads, or respiratory pathology^46,179^ observed in NHPs exposed via aerosol to lower doses than multi-route exposures. Contrasting findings across different plausible severity metrics have also occurred in other pathogen-host systems; for example, one NHP study on influenza H5N1 noted worse lung CT scores following multi-route exposure than aerosol inhalation^180^, while another found more widespread lower respiratory infection for aerosol inhalation than multi-route inoculation^181^. It will remain difficult to identify consensus patterns until the best correlate of severity is identified for non-lethal animal models (like SARS-CoV-2 in NHPs) and until the dependence of infection outcomes on IRP sizes and other methodological aspects of aerosol exposure are better characterized in these systems. Future work could adapt our framework for these investigations, namely to explore the relationships between AUC values, symptom severity scores, and other clinical metrics.

As is common for respiratory pathogens^6,16,24^, we approximated shedding potential using infection AUC values in the upper respiratory tract. These analyses revealed that aerosol inhalation results in significantly higher nasal shedding than other respiratory exposure routes. When coupled with evidence that viruses exhaled from the upper respiratory tract of donor animals drive transmission to sentinel animals for SARS-CoV-2 and influenza^156–159^ and that SARS-CoV-2 transmission success increases alongside viral titers in nasal lavage samples of donor animals^160^, this result suggests that aerosol-exposed individuals may pose higher transmission risk. Transmission experiments could directly test this hypothesis by including aerosol-exposed donor animals in addition to standard intranasally-inoculated donors.

Despite compiling the largest ever database of non-human primate challenge experiments, limitations in the available data restricted our analyses. Available doses and overall sample sizes in our database were starkly imbalanced across routes: those including combined intranasal/intratracheal inoculation comprised 62% of the full database, and aerosol inhalation was the only route with doses below 10^3.5^ pfu. The popularity of high-dose, multi-route exposures may reflect investigators’ desire to guarantee successful infection, but they limit the scientific scope of each study and of all retrospective analyses to doses well above transmission bottleneck sizes estimated by genetic means (<50 viruses for SARS-CoV-2^182^; ∼2 genomes for two influenza A subtypes^183^), and they obscure the possible influence of particular exposure routes. Lower experimental doses should still result in successful infections in this system, given that our ID50 estimates based on PCR detection did not exceed 10^1.3^ pfu for any respiratory exposure route or tissue. Recent calls to prioritize studies that translate more clearly to human applications^184^ reinforce the need for animal experiments to use more naturalistic dose ranges. Such data would enable future work to determine whether dose effects are more prominent at lower ranges, and whether this alters our finding that route drives kinetics more strongly than dose. A striking number of studies in our database also did not publish their source data or link individual-level data across figures, which limited the information that could be extracted from these studies. We hope that recent data-sharing mandates^185^ and the success of this multi-protocol meta-analysis will inspire more comprehensive reporting and thus maximize the scientific discoveries from hard-earned experimental data.

This study focused on SARS-CoV-2 in NHPs because copious NHP data was generated during the COVID-19 pandemic and because NHPs are an important animal model for translational research^14^. Many of the patterns we identified align with findings in other pathogen-host systems that were obtained more qualitatively in smaller sample sizes, including strong dependence of virus kinetics on route^3–6,12,13,148,149^ and higher titers at larger doses^8,13,16,186^. Variable protocols and constrained sample sizes, however, can lead to individual studies identifying conflicting patterns (e.g., similar^99^ or worse^34^ lung pathology for SARS-CoV-2-infected cynomolgus vs. rhesus macaques), which make it difficult to establish clear overarching principles. Our quantitative framework can and should be applied to aggregate datasets for other pathogens and host species to resolve existing discrepancies and further probe similarities. Influenza A infection in ferrets offer a promising next step for these analyses and would extend recent efforts to aggregate data from studies with standardized protocols in this system^24,138,140^. The framework could also be used to analyze clinical and challenge trial data from humans, enabling a formal connection between animal model studies and their biomedical applications. Meta-analytical approaches like ours could reveal unifying principles that shape viral kinetics and pathogenesis. Whether exposure route has equally strong effects on within-host kinetics for other coronaviruses, for other respiratory viruses, or for other pathogen families is a key question for subsequent work.

Our study presents the most comprehensive and quantitative characterization to date of how exposure route and dose shape infection patterns inside hosts, in ways that affect both disease risk and transmission potential. With our work, we demonstrate the ability of meta-analysis to answer fundamental questions that are inaccessible for individual experiments (such as comparing the effects of exposure route and dose), ultimately enhancing the value of each constituent study while reducing the total number of animals needed for scientific discovery. Future studies can increase their statistical power by comparing their results to our database of historical control individuals. Possible applications include comparing the viral kinetics of distinct SARS-CoV-2 variants, evaluating the efficacy of pharmaceutical interventions, and identifying differences by host or pathogen species. Quantitative tools based on mechanistic principles show immense promise to illuminate pathogen-host biology, and to launch a new era of microbiology that leverages computation to magnify the benefits of animal experiments and improve human health.

## Online Methods

### Database compilation

We have previously described the construction of our core database (Snedden and Lloyd-Smith 2024), which contains SARS-CoV-2 viral load and infectious virus data from non-human primate experiments. We expanded the database for this study by conducting another systematic literature search with the same keywords. The first search included articles from January 1, 2020 through March 11, 2021, and the second search spanned March 12, 2021 through April 9, 2024 (**Fig. S1**). For both searches, we identified articles that met the following criteria: (i) they experimentally infected rhesus macaques (*Macaca mulatta*), cynomolgus macaques (*Macaca fascicularis*), or African green monkeys (*Chlorocebus sabaeus*) with SARS-CoV-2 (restricted to basal strains, excluding any studies that reported a named variant or reported that their virus included the D614G sequence; we did not screen genome sequences deposited in repositories), and (ii) they published quantitative or qualitative measurements of viral load (measured by RT-qPCR) or infectious virus (measured by plaque assay or endpoint titration) from at least one biological specimen for at least one individual and at least one sample time post infection.

Of the studies that met our criteria during the first search, single-route exposures (e.g., intranasal only, aerosol) were the least common and had more limited dose ranges. To increase sample sizes for these crucial exposures, we only considered studies with single-route exposures during our second search. Altogether, our database includes 107 articles (**Fig. S1; Table S1**). For both searches, raw data were used when available (published or obtained via email correspondence); otherwise, one author (CES) extracted data from published figures using the package ‘digitize’ in R.

### Anatomical characterization

In this study, our analyses focused on the nose, throat, trachea, lung, upper gastrointestinal tract, and the lower gastrointestinal tract (GI). We categorized all reported sample types into these six tissue categories, and we distinguished between invasive and non-invasive sampling methods (**Table S3**). We also grouped the six tissues into three broader categories as follows: (i) upper respiratory tract (URT): nose and throat; (ii) lower respiratory tract (LRT): trachea and lung; and (iii) gastrointestinal tract: upper and lower GI. We refer to these broad tissue categories as ‘organ groups.’

### Key metrics for characterizing kinetics

To guide our analysis, we selected five key metrics to characterize within-host infection kinetics in any given tissue for any individual, namely: (i) the probability of ever testing positive, (ii) the time until becoming detectably infected, (iii) the time until reaching the peak titer, (iv) the quantitative peak titer, and (v) the time until the infection becomes undetectable (**Fig. 1c**). From the five primary metrics, we also calculated the duration of detectable infection (the time elapsed between becoming detectable and undetectable) and the area under the infection curve (0.5 * the duration of detectable infection * the peak titer). We extracted all of these metrics for the following five primary tissues that can be sampled repeatedly by non-invasive methods: nose, throat, trachea, lung, and the lower gastrointestinal tract. For the upper gastrointestinal tract, which was only accessible by invasive methods, we only estimated the probability of testing positive and the time until becoming detectably infected.

### Extracting metrics for animals with ID names

Given that sampling occurs at discrete times, our metrics cannot be observed exactly. To account for this, we extracted bounded intervals during which the events could have occurred for every individual that had data for any given tissue location and assay type. Brief descriptions of these extraction methods are outlined below for each metric, with a hypothetical example shown in **Fig. 1c-e** and example individuals from the database illustrated in **Fig. S4**. We included both invasive tissue samples as well as non-invasive sampling methods (e.g., swabs) to characterize event times, where invasive samples contribute information on the probability of positivity, time to detectability, and the time to undetectability, but not the peak titer. Non-invasive samples contribute to all metrics.

*Probability of testing positive and time to detectability.* For all individuals that tested positive in a given tissue location for a given assay, we bounded their time to detectability between the day of their first observed positive test and the day of their previous negative test. For individuals that never tested positive in that tissue for that assay, we considered two possibilities: either the individual would have never tested positive or they would have tested positive if observed later post infection. In the latter case, we assumed that their time to detectability must be bounded between their last observed negative test and day forty-five post infection, reflecting that a month and a half (or more) delay for initial positivity is highly unlikely.

*Peak time and peak titer.* For all individuals with positive quantitative titers for at least two days post inoculation, we identified the day with their largest observed titer. We used the previous and subsequent sampling days as the lower and upper bounds, respectively, for that individual’s time to peak titer. If the individual’s highest titer was observed on the final sampling day, then we set the upper bound as day fifty post infection. We used the titer from the observed peak time for the peak titer model. For all animals whose peak titers were reported in TCID50, we converted them to pfu using a standard conversion factor (1 TCID50=0.69 pfu^187^).

*Time to undetectability.* For all individuals with at least one negative sample after their final observed positive sample, we bounded their time to undetectability between the day of their last observed positive sample and their subsequent (negative) sample. If an individual tested positive on their final sample day, then we set their upper bound as day fifty-five post infection, reflecting that prolonged infection beyond approximately two months post inoculation is highly unlikely in otherwise healthy animals.

### Extracting metrics for animals without ID names

Some studies presented their data in figures without correlating observations to particular animals across time points (e.g., scatterplots without lines connecting unique animals). We were still able to extract some censored event times in these instances, though fewer and with worse resolution than if the data were labelled. We used the following approaches:

*Probability of testing positive and time to detectability.* We identified the first day that any animal(s) tested positive by a given assay in a given tissue, which we used to extract interval censored event time(s) with the previous sampling day as the lower bound. We then searched all subsequent sampling days to check if more animals tested positive in that tissue by that assay on any other day. If so, we extracted an interval censored event time for each *new* animal that tested positive by that day. For example, if two animals tested positive on day two but three animals tested positive on day five, then at least one additional animal became detectable between day two and day five. If this process resulted in fewer interval censored event times than the total number of animals shown in the figure, we extracted right censored event times for the additional animals, where the lower bound was the first day that any animal tested positive and the upper bound was day forty-five (as for the animals with ID names).

*Peak time and peak titer.* The day post infection with the largest observed titer across all data points for a given tissue and detection assay must include the peak titer for one animal. In this case, we extracted one interval censored peak time with bounds set by the previous and subsequent sampling days on which animals were tested. On the same day of that first animal’s peak titer, we determined if any of the other titers reported exceeded all other titers reported on all previous or subsequent sampling days. For the titers bigger than those on all previous *and* subsequent days, we extracted another interval censored event time with bounds set by the previous and subsequent sampling days on which animals were tested. For the titers bigger than those on all previous days but not all subsequent days, we extracted a right censored event time with a lower bound set by the previous sample day on which animals were tested. If there is ever a subsequent sampling day on which *all* animals test negative, we instead use that as the upper bound for an interval censored time. For the titers bigger than those on all subsequent days but not all previous days, we set an interval censored time with day 0 as the lower bound and the subsequent sampling day on which animals were tested as the upper bound.

*Time to undetectability.* For the animals where we were able to extract peak titers, we set conservative bounds on the time to undetectability. In particular, we used right censored event times such that the time to undetectability must occur after the day of the observed peak titer when available or after the lower bound on the day of the peak titer.

### Modeling framework

Survival analysis methods can estimate event times from bounded (or ‘censored’) observations^29^. We used these techniques as a basis to develop our Bayesian model, given that, during animal infection experiments, samples are collected at discrete times to study an on-going infection process. As described above, we characterized within-host kinetics for all considered tissues according to three event times: the time to viral detectability (TD), the time to peak viral titer (TP), and the time to viral undetectability (TU). We used an accelerated failure time model for all three, where we assumed that the time delay between any two sequential events (i.e., between inoculation and TD, between TD and TP, and between TP and TU) was distributed according to the Weibull distribution, which allows for flexible event time distributions. We fit our model to the median of each distribution, which is given by the equation λ(ln2)^1/k^, where λ is the Weibull scale parameter and k is the Weibull shape parameter. We assumed that the medians on the log scale were predicted by a linear combination of a set of j covariates (X_j_; see next section), all with their own unique regression coefficients (β_TD,j_, β_TP,j_, β_TU,j_). On top of those standard covariates, the time to peak titer had an additional term that captured any dependence on the time to detectability (β_TP,TD_TD), and the time to undetectability had an additional term to capture any dependence on the peak titer (β_TU,PT_PT). We also estimated Weibull shape parameters (k_TD_, k_TP_, k_TU_) that were shared among cofactors but unique to each event time and organ group (URT, LRT, GI). For numerical stability, we required that these shape parameters be greater than or equal to one.

To capture that some individuals may never test positive (e.g., because of failed infection in a given tissue or low assay sensitivity), we adapted our time to detectability model into a mixture cure model^188^ that also estimated whether an individual will ever test positive (P) in a given location. We assumed that P was Bernoulli distributed with probability p, which is predicted by a linear combination of the standard j covariates (β_P_X_P_) with a logit link function. Thus, the overall probability that an individual will test positive by a given time *t,* which we represent by the function S(*t*), depends both on p as well as the probability of testing positive by time *t* when conditioned on ever testing positive (S_p_(*t*)). All other metrics are conditional on the individual ever testing positive.

We modeled peak titers differently, as they are not event times. We assumed that all true peak titer values (in log10 units; PT_true_) are normally distributed, where the mean is a linear combination of our standard j covariates with their own regression coefficients (β_PT_X_PT_). We included an additional term to capture any dependence of the peak titer on the time to peak titer (β_PT,TP_TP). We allowed this relationship to differ between exposed and not exposed tissues, because exposed tissues often peak within the first day, possibly due to resampling the inoculum. The standard deviation of these titers (σ_PT_) is specific to the organ group. Given that an individual’s true peak titer likely differs from their observed largest titer (PT_obs_), we treated PT_obs_ as normally distributed around the sum of PT_true_ and a term that captures lab and assay effects (δ_lab,assay_). When estimating true peak titer values, we bound them between one order of magnitude less than the observed peak titer and 14 log10.

To capture quantitative differences in all metrics due to methodological variation among articles, we included linear offset terms for article and assay effects (see *Covariates* for more details). We also allowed the standard deviations of the observed peak titers to differ across articles, given that sampling schemes and other methodological choices may affect variability in observed titers.

Below is the general form of this model, where ɑ are the intercept parameters and all other parameters are described above. For numerical stability when fitting, the models for event times included rescaling by-1/10.

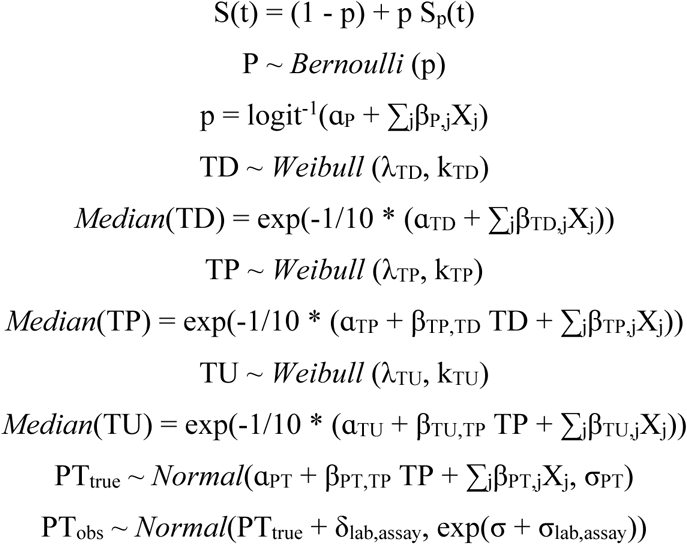

As described above, we cannot exactly observe when an individual experiences their three true event times (TD_true,i_, TP_true,i_, TU_true,i_). Instead, we treated them as bounded quantities, such that the true time for any individual *i* occurs within their lower and upper bound (e.g., [LB_TD,i_, UB_TD,i_] for TD_i_). Each individual’s true times must occur within these intervals, where the probabilities of any given event time within that interval are given (as above) by the Weibull distribution for their unique combination of cofactors (e.g., TD_true,i_ ∼ *Weibull*(λ_TD,i_, k_TD_) where the median is *exp*(-1/10 * (ɑ_TD_ + ∑_j_β_TD,j,i_X_j,i_))). In order to model the time delay between sequential events for each individual when fitting our model, we adjusted the upper and lower bounds as follows: for the time to peak titer, we subtracted both bounds by the estimated waiting time until detectability; for the time to undetectability, we subtracted both bounds by the sum of the estimated waiting time until detectability and waiting time until peak titer. These lower bounds were not allowed to be negative, which enforces the sequential nature of these event times.

### Covariates

We allowed all of our metrics to vary across multiple covariates, which have been hypothesized to influence within-host kinetics. All categorical cofactors with more than two groups (exposure route, age, species, article, assay) were treated as unordered index variables. Binary predictors (sex, tissue location) were treated as indicator variables. When age, sex, assay, or tissue location (within an organ group) were unknown, we marginalized over all possibilities.

### Biological cofactors

We included exposure route as a categorical cofactor with the following classifications: (i) upper respiratory exposure, (ii) lower respiratory exposure, (iii) combined upper and lower respiratory exposure via liquid administration, (iv) combined upper and lower respiratory exposure via aerosol inhalation, and (v) upper gastrointestinal exposure. We grouped all reported inoculation routes (e.g., ocular, intranasal) into these categories (**Table S2**). For all analyses, we generated predictions for the following representative route for each category: (i) intranasal, (ii) intratracheal, (iii) equal combination of intranasal and intratracheal, (iv) aerosol, and (v) intragastric.

We included exposure dose by stratifying the total inoculation dose (in log10 pfu) into tissue-specific categories. For every individual, we calculated the dose administered specifically to the nose, throat, trachea, lung, and stomach. We assumed that the total dose was split among locations for each inoculation route as follows: (i) ocular: 100% nose; (ii) intranasal: 50% nose, 50% throat; (iii) oral: 100% throat; (iv) intratracheal: 100% trachea; (v) intrabronchial: 100% lung; (vi) aerosol: 25% nose, 25% throat, 25% trachea, 25% lung; and (vii) intragastric: 100% stomach. We assumed ocular inoculation resulted in nasal exposure, given that fluids administered to the eye rapidly drain into the nose via the nasolacrimal duct^189^. A small number of studies included intravenous inoculation in addition to other routes (**Table S2**), but we did not include a parameter in our model for this given small sample sizes. When studies reported inoculation dose as TCID50 values, we converted to pfu using a standard conversion factor (1 TCID50=0.69 pfu^187^). When fitting the model, we divided the raw dose in pfu administered in each tissue for each individual by the maximum dose administered in that tissue across all individuals, such that tissue-specific doses ranged from 0 to 1.

We included age, sex, and non-human primate species as our demographic effects. For species, we distinguished between rhesus macaques, cynomolgus macaques, and African green monkeys. For age class, we distinguished between juvenile, adult, and geriatric individuals, where we assigned age classes as described in our earlier work^27^. Briefly, ages less than 5 years were considered juveniles for all three species. Adults included ages 5-19 years for both macaque species and 5-14 for African green monkeys. All older individuals were considered geriatrics.

Finally, we included a binary predictor for tissue location to account for differences between the two distinct tissues studied within an organ group (i.e., for nose and throat in the model of the upper respiratory tract, and for the trachea and lung in the model of the lower respiratory tract).

### Methodological cofactors

We incorporated multiple cofactors to handle methodological variation among articles. Given that virus detection and virus quantification can vary across assays (e.g., PCR vs. virus culture), we included a cofactor for assay type with the following four categories based on our prior work^27^: (i) total RNA assays targeting the N gene (referred to as ‘total RNA’ in all analyses), (ii) any genomic RNA assay or total RNA assays targeting less expressed genes (referred to as ‘gRNA’ in all analyses), (iii) sgRNA assays (referred to as ‘sgRNA’ in all analyses), and (iv) plaque assay (referred to as ‘culture’ in all analyses). To account for differences in other methodological factors among articles (e.g., RNA extraction methods, viral strains), we included a linear term for each article and assay combination. For example, if an article included data from both a total RNA and a culture assay, then that article would have its own unique offset term for each assay.

### Parameter sharing within organ groups

Each organ group (URT, LRT, and GI) has their own set of equations, such that, for example, the nose and throat share most regression coefficients with each other but they do not share any with the LRT or GI tissues. In particular, the following parameters are shared within organ groups: intercepts, demographic effects (age, sex, species), location effect, assay effects, lab effects, and the dependence on previous metrics (i.e., β_TP,TD_, β_PT,TP,_ β_TU,TP_).

### Cofactor combinations

When generating predictions from our model, it is necessary to choose a route, total dose, age, sex, and species of interest. For most of our analyses, we focus on total doses of 10^4^ and 10^7^ pfu. Given there are five exposure route categories, two focal dose values, three species, three ages, and two sexes, there are 180 possible combinations of cofactors to choose from. We refer to any one of these 180 options as an individual ‘cofactor combination.’ In many of our analyses, we report results that include predictions for all 180 possible cofactor combinations, or more cofactor combinations when we consider additional doses. Because of parameter sharing, our model can generate predictions for cofactor combinations that do not exist explicitly in the dataset used for fitting, and we use these predictions in our analyses. See **Fig. S14** for comparisons of the predictions generated for cofactors within the dataset and those extrapolated from shared parameters.

### Prior probabilities

For all covariates in our model, we used weakly informative priors to exclude implausible relationships, but we made no *a priori* prediction on the direction of those individual effects (i.e., the priors for each cofactor were centered on zero). The only exception was for the relationship between dose and probability of positivity, where we assumed that larger doses cannot decrease the probability of positivity and they most likely increase the probability. Prior predictive simulations confirmed variable but reasonable expectations for all metrics.

### Model predictions and determining significance

For all presented results, we used 1000 post-warmup samples of the model posteriors to generate predictions unless otherwise noted. Each prediction is generated using grouped parameter samples (e.g., samples from the same chain and iteration) to preserve correlation structure. We also assume there is no article effect when generating predictions, unless otherwise stated (i.e., we set all article terms to zero). Unless otherwise noted, all visualizations also integrate predictions for all ages, sexes, and species. To determine whether any differences in predictions were significant, we analyzed 90% prediction intervals. If they did not include zero, then we considered the difference to be significant.

### Ranking the importance of biological cofactors

To determine which biological cofactors had the largest impact on each metric for each tissue, we generated predictions for all cofactor combinations. From these predictions, we calculated the differences for all possible pairwise combinations of the categories within each cofactor (e.g., juvenile vs. adult, adult vs. geriatric, and juvenile vs. geriatric for age). Then, for each cofactor and metric, we computed the mean value of their differences. The cofactor with the largest such mean for a given metric thus had the largest average difference among its categories and thus exerted the greatest influence on that given metric. We assigned this cofactor an effect of ‘1’ for that metric. Finally, we calculated metric-specific relative effects for all other cofactors by scaling their mean differences by the largest mean difference.

### Generating a within-host connectivity structure

To generate our tissue connectivity structure in **Fig. 4**, we first used our model posteriors to obtain predictions for the probability and time to detectability in the nose, throat, trachea, lung, upper GI, and lower GI following all possible single-tissue inoculations individually. For example, we generated predictions following the administration of 10^7^ pfu only into the nose, which are distinct predictions from those generated following inoculation only into the throat. We considered nose-only, throat-only, trachea-only, lung-only, and upper GI-only inoculation. Our database does not include lower GI-specific inoculation, so we did not consider this scenario. We used 1000 samples of our model posteriors to generate predictions for each tissue-specific inoculation. We consider all possible combinations of species, age classes, and sexes. Given PCR detection is more sensitive than viral culture, we only consider predictions when monitoring infection via total RNA PCR. For each sample, we determine whether each non-inoculated tissue will test positive by randomly sampling from a Bernoulli distribution where the probability is the predicted percent of individuals that test positive for that sample. With these predictions, we then characterized the directional flow of virus out from each inoculated tissue by computing a unique 6×6 adjacency matrix for each single-tissue inoculation, where the columns and rows correspond with the six tissues. We filled in this matrix by incrementing the [X, Y] entry by 1 for each sample where tissue Y is predicted to become detectable after tissue X, under the following assumptions: (i) the inoculated tissue becomes detectable immediately, and is followed by all other tissues that are predicted to become detectable before day 1 post infection (i.e., each other tissue that tests positive before 1 dpi has its adjacency score with the inoculated tissue increased by 1), (ii) whenever a new tissue tests positive, the adjacency score is increased by 1 with all tissues that became detectable on the previous day that any new tissue became detectable, and (iii) all tissues that are not predicted to test positive do not increment any connections for that particular parameter sample. These analyses result in 5 distinct adjacency matrices, namely those following nose-only, throat-only, trachea-only, lung-only, and upper GI-only inoculation (**Fig. S25**).

With these 5 adjacency matrices, we first characterized outflows from each tissue (**Fig. 4a**). The most direct and least confounded information on outflow from tissue X stems from infections following inoculation only into tissue X. As such, for each tissue X, we focus on row X in the adjacency matrix for inoculation of that tissue. Within this row, we rank each tissue Y by how likely tissue X is to flow into that tissue, with higher values indicating higher flow. The largest value thus corresponds with the tissue Y that tissue X primarily flows to. Conducting these calculations for each tissue X results in the row-wise orderings shown in **Fig. 4a**, where each row is extracted from the corresponding analyses conducted on that tissue-specific matrix. It is important to note that these predictions incorporate spread between tissues that may result from the direct flow of the inoculum to nearby tissues or from virus that has replicated in the inoculated tissue and then disseminated. Both offer valuable information about tissue connectivity; disentangling them would require samples at higher resolution before day 1 post infection.

To obtain a more comprehensive picture of connectivity, we then characterized the tissues that primarily flow into each tissue Y (**Fig. 4b**). When tissue Y is inoculated, we assume tissue Y is immediately detectable. Thus, the corresponding adjacency matrix for tissue Y cannot contribute information on inflow to tissue Y, and so we excluded it for analyses of inflow into tissue Y. Instead, we focus on column Y in the adjacency matrices for the other four inoculated tissues, which offer information on virus inflow to tissue Y when other tissues are inoculated. We calculate an aggregate inflow value by computing the elementwise mean values of column Y across those four adjacency matrices. For this mean column, we rank each tissue X by how likely tissue Y is to receive inflow from that tissue, with higher values indicating higher flow. The largest value thus corresponds with the tissue X that tissue Y primarily receives flow from, averaging over multiple inoculation routes. Conducting these calculations for each tissue Y results in the column-wise orderings shown in **Fig. 4b**, where each column is extracted from the corresponding analyses conducted for that tissue Y.

To create the connectivity network in **Fig. 4c**, we include directional arrows connecting all tissue pairs with a rank ‘1’ in the outflow (**Fig. 4a**) and inflow (**Fig. 4b**) matrices.

### Computational methods and software

All data preparation, analysis, and plotting were completed with R version 4.2.0. Posterior sampling of the Bayesian model was performed with No-U-Turn Sampling (NUTS) via the probabilistic programming language Stan using the interface CmdStanR version 0.5.2. All model fits were generated by running four replicate chains with 4000 iterations each, of which the first 2000 iterations were treated as the warmup period and the final 2000 iterations were used to generate parameter estimates. Model convergence was assessed by the sampling software using 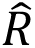, effective sample sizes, and other diagnostic measures employed by CmdStan by default. No issues were detected.

## Data Availability

The full database compiled during the literature search will be made available on Zenodo at the time of publication, as will the dataset of censored event times used for analysis in this study.

## Code Availability

All code used to produce the results and figures in this manuscript will be made available on Zenodo upon publication.

## Acknowledgements

Our study would not have been possible without the immensely valuable research completed by the authors of the articles included in our database. We sincerely thank all of these researchers, many of whom kindly corresponded with us to provide clarifications and raw data. J.O.L.-S. and C.E.S were both supported by the Defense Advanced Research Projects Agency DARPA PREEMPT #D18AC00031. C.E.S was also supported by the National Institutes of Health (grant 5T32 GM008185-33) and the UCLA Office of the Vice Chancellor for Research 3R Grant. J.O.L.-S. was also supported by the National Science Foundation (DEB-1557022 and DEB-633 2245631) and the UCLA AIDS Institute and Charity Treks. T.C.F. was supported in part by the National Institutes of Health Office of Research Infrastructure Programs (NIH-ORIP) grant P51OD011106. D.H.M. is an employee of the US Centers for Disease Control and Prevention, but conducted this work in a personal capacity, as an uncompensated consulting scientist working with the J.O.L.-S. lab at UCLA. The content of the article does not necessarily reflect the position or the policy of the US government or the Centers for Disease Control and Prevention, and no official endorsement should be inferred.

## Author Contributions

C.E.S and J.O.L-S conceptualized the study. C.E.S, D.H.M., and J.O.L-S. developed the statistical model. C.E.S. completed the systematic literature review, extracted all data, and curated the database, with guidance from J.O.L-S. C.E.S. wrote the code for the statistical model, for model fitting, for analyzing the model outputs, and for visualizing results. T.C.F. provided insight into the design of animal experiments and contributed virological expertise to the interpretation of our findings. C.E.S. wrote the original draft. C.E.S, D.H.M, T.C.F, and J.O.L-S reviewed and edited the manuscript.

---

**Extended Data Figure 1.**
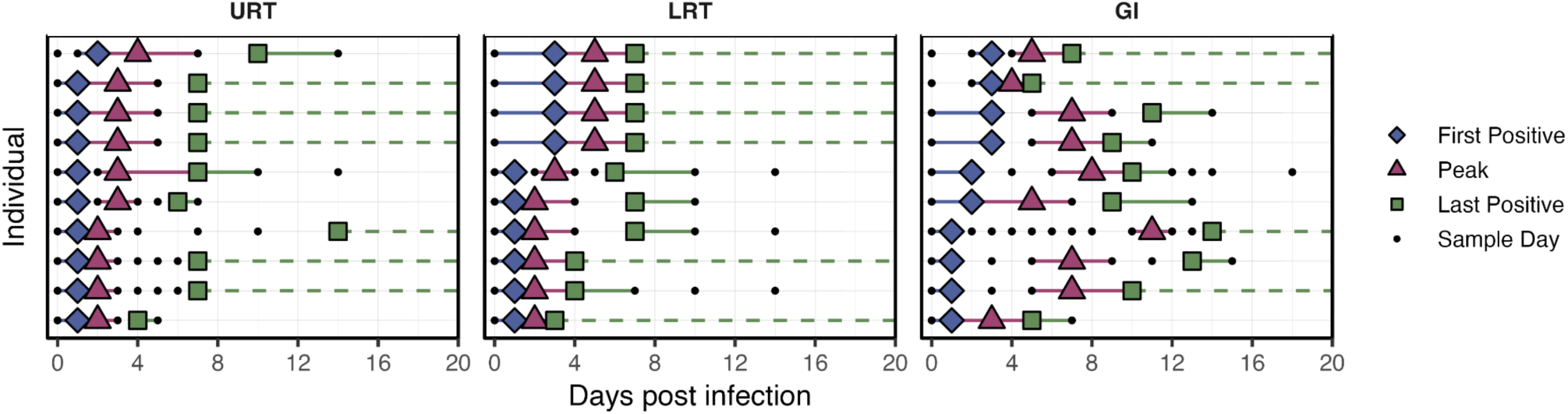
Example censored data used for model fitting. These data are a small subset of what we used to fit our model. Each row within each panel gives a unique individual’s time series for the location indicated in the panel label. Observed event times are shown with colored shapes (first positive, blue diamond; largest titer, purple triangle; last positive, green square), and their censored intervals are indicated by the colored lines. Solid lines are interval-censored observations. Dashed lines indicate the individual was positive on their final sample day, so their event is right censored. Individuals can differ across rows in different panels. For visual clarity, only individuals with multiple positive samples and distinct first positive and peak titer days are shown as examples for this figure. See **Fig. S4** for more examples.

**Extended Data Figure 2.**
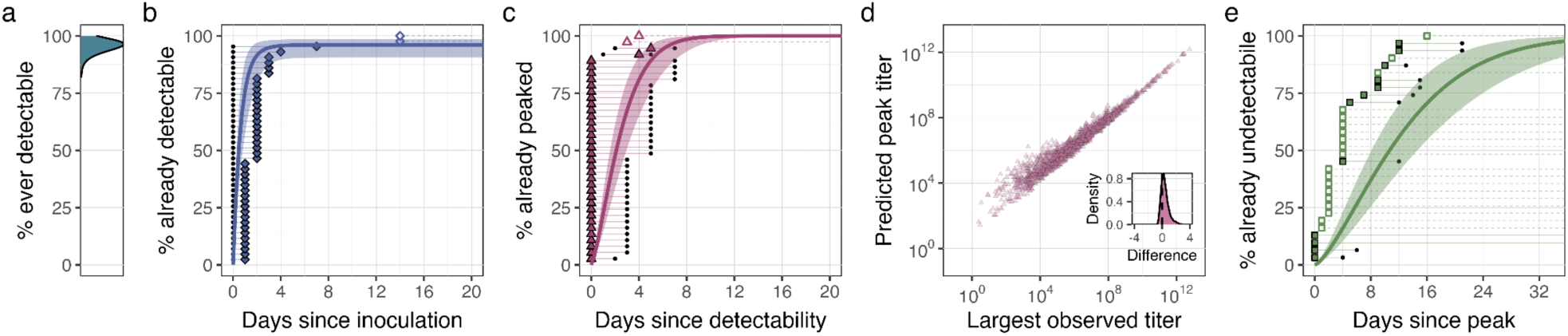
Example model fits to data. All panels except d show model predictions based on 500 posterior samples ranging from 10^4^-10^7^ (in increments of 1 order of magnitude), specifically for IN-exposed individuals sampled by total RNA assay. The plotted data are also restricted to these cofactors but allowing all PCR assays. Models are fit to the censored intervals and not to the symbols. Filled points are interval-censored observations, and open points with dashed lines are right-censored observations. **a,** Model predictions for the percent of individuals that will ever test positive, corresponding with the observed event times in b. **b, c, e**, Median model fit (bold line) and 90% credible intervals (shaded regions) against individual bounded event times (points in rows) for the times to (b) detectability, (c) peak titer, and (e) undetectability. The y axis displays the percent of individuals predicted to have experienced the event by the given days since the previous event. Model fits to all available data stratified by exposure route and tissue location are shown in **Figs. S5-S7**. **d**, Relationship between the largest observed titer and the mean true peak titer predicted by the model. Each point is an individual sampled in a given tissue by any PCR assay. Mean predictions use all samples from the model for the given individual. The inset shows the distribution of differences between the true and observed titers. **Fig. S8** shows results for all assay types.

**Extended Data Figure 3.**
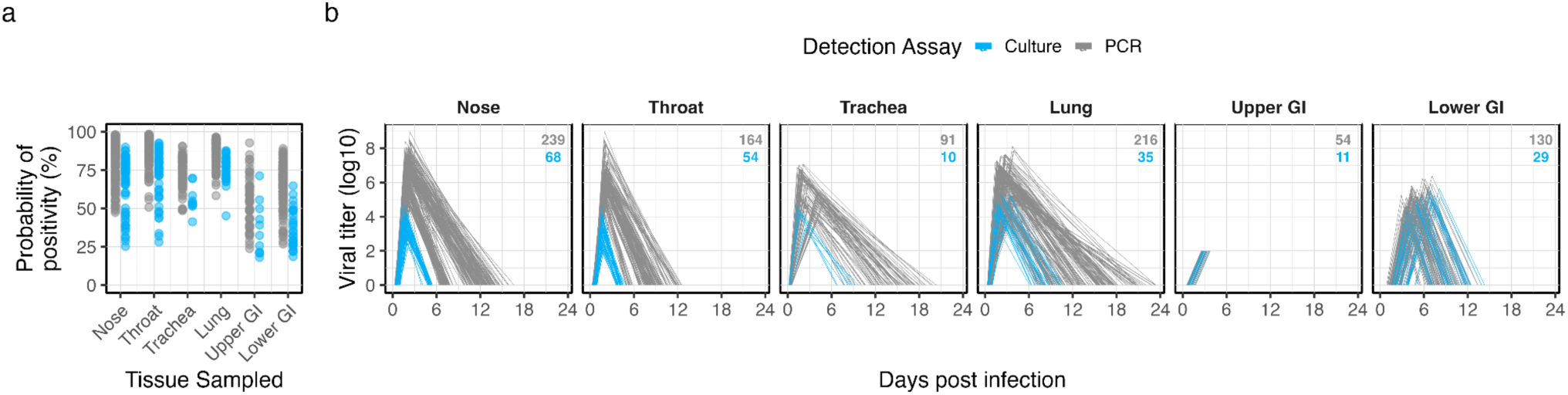
Model predictions for the unique covariates used in each article. Results show median model predictions for the exposure route(s), dose(s), age(s), sex(es), species, and assay(s) used in each article and for the tissue(s) each article sampled. Each point and line corresponds with the median prediction for one such covariate combination. **a**, Predictions for the probability of positivity. **b**, Predictions for infection trajectories, stratified into panels by tissue. All PCR assays are in grey; culture assays are in blue. Annotated numbers in **b** give the number of lines plotted for each assay category in the given panel. Upper GI trajectories are truncated because we can only infer the times to detectability. **Fig. S9** shows results further stratified by exposure route and assay type.

**Extended Data Figure 4.**
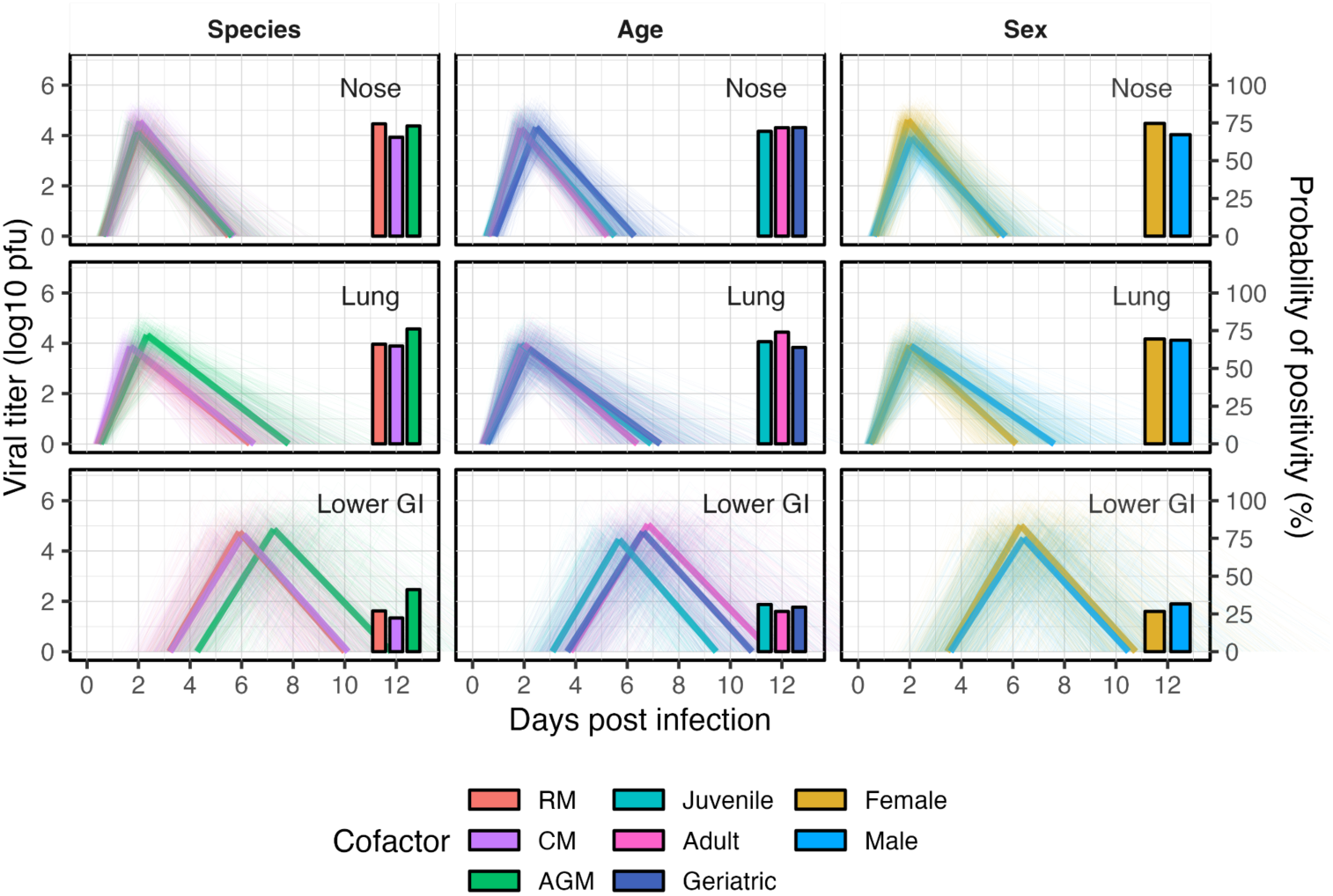
Effects of demographic factors on within-host kinetics for aerosol exposure. Each row of panels displays model predictions for one tissue (top: nose; middle: lung; bottom: lower GI), and each column stratifies predictions based on the cofactor indicated in the column label (e.g., Species distinguishes results among rhesus macaques [RM], cynomolgus macaques [CM], and African green monkeys [AGM]). Colors within a column of panels distinguish among the constituent cofactors, as shown in the legend. Thick, opaque lines are median infection trajectories following aerosol-exposure with 10^4^ pfu. Thin, transparent lines are trajectories from 100 samples of model posteriors. Bar plots centered at 12 days post infection display the median predicted probability of positivity, which corresponds with the y axis label on the right-hand side.

**Extended Data Figure 5.**
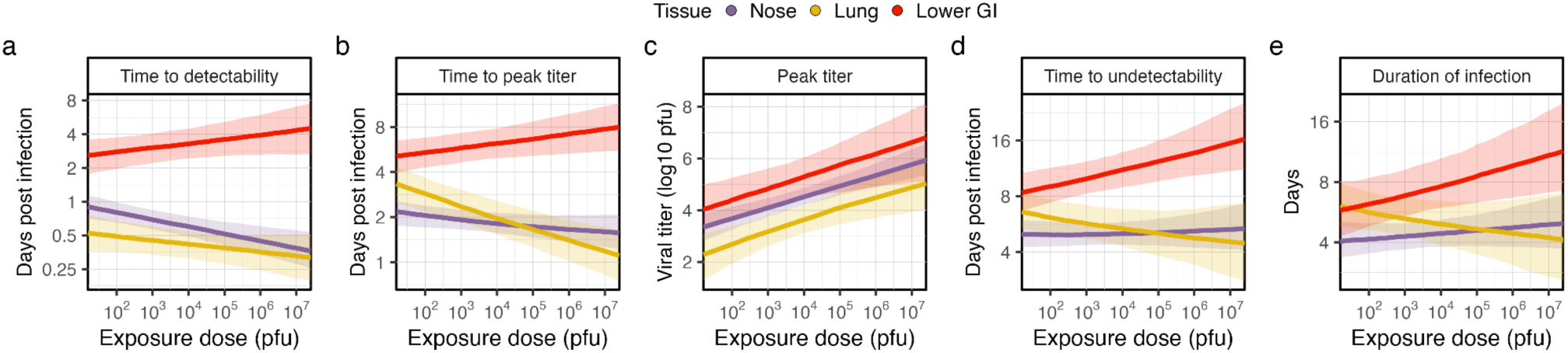
Effects of aerosol exposure dose on spatiotemporal spread of infectious virus. Thick, dark lines are the median predictions. Shaded regions correspond with the 90% prediction interval. Color distinguishes between tissues as indicated in the legend (purple: nose; yellow: lung; red: lower GI). Panels present results for our infection metrics, as follows: **a**, time to detectability; **b,** time to peak titer; **c**, peak titer; **d**, time to undetectability; and **e**, duration of infection. Results in all panels are for a female, adult, AE-exposed rhesus macaque.

