## Supplementary Figures and Tables for "Exposure route drives SARS-CoV-2 infection patterns in non-human primates"

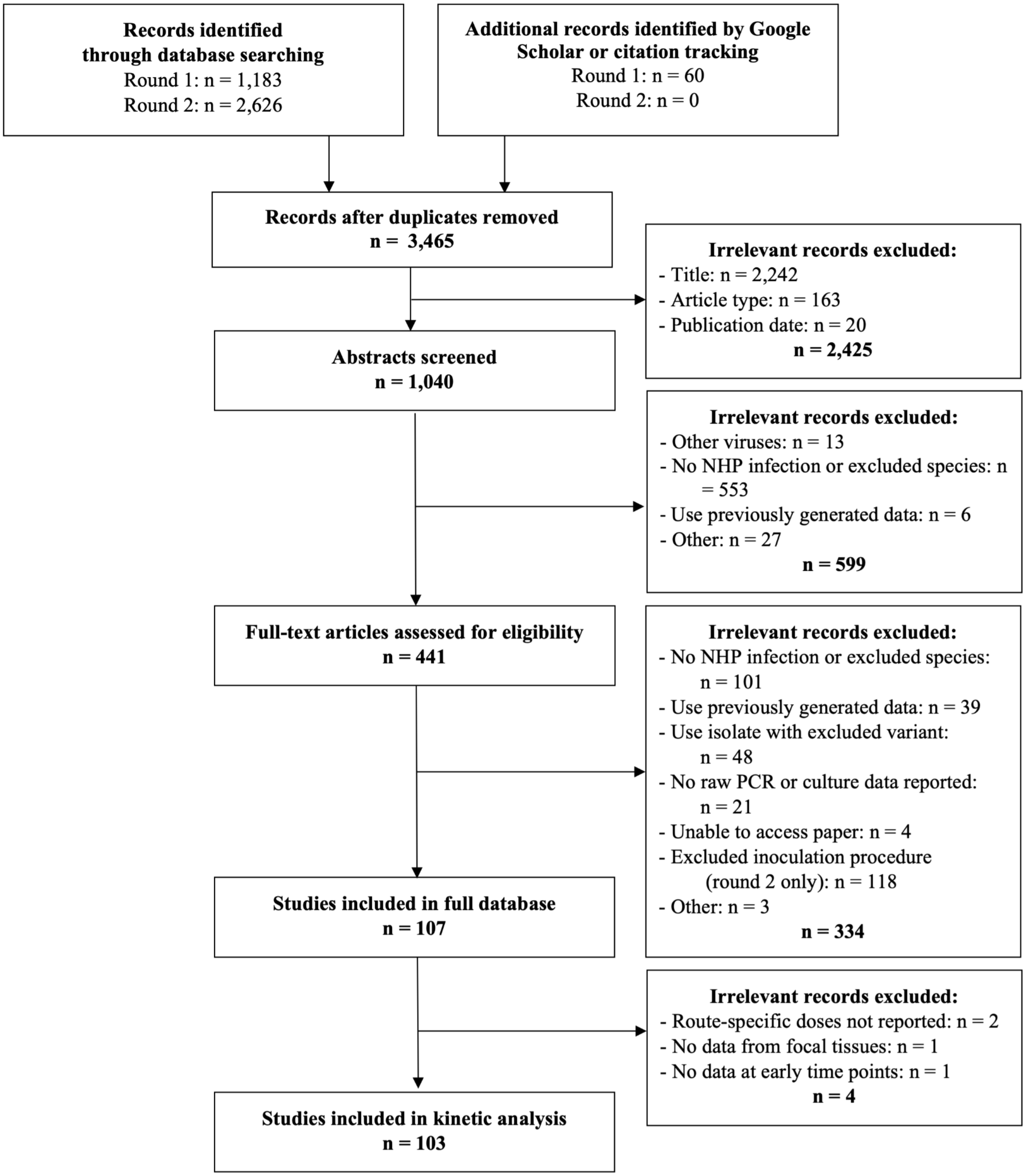


**Figure S1. Screening and selection procedure for the compilation of the full database and for the construction of the dataset used for kinetic analyses in this study.**

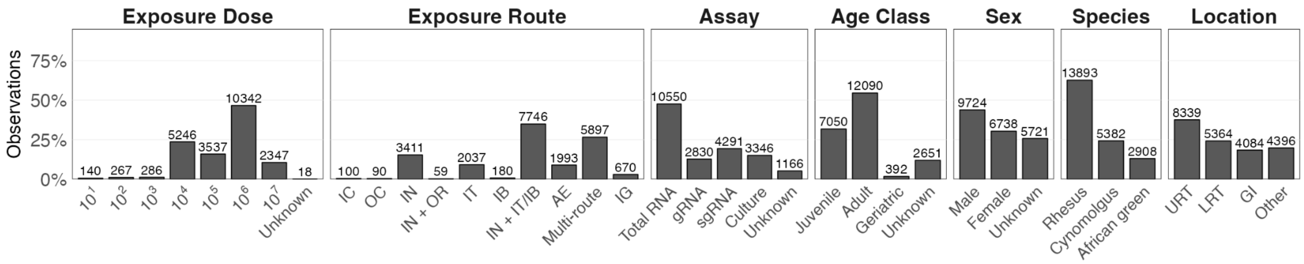


**Figure S2. Data distribution across key cofactors for the full database.**

Bar heights give the percentage of all samples, and annotated numbers give the total number of observations in the full database. One observation is one sample time (e.g., day 1 post infection) in one tissue location (e.g., nose) by one sampling method (e.g., nasal swab is separate from nasal tissue sample at necropsy) processed by one assay type (e.g., sgRNA PCR is separate from total RNA PCR), all from one individual. One animal may yield multiple observations if they were sampled at multiple time points or in multiple locations, or if their sample was run with multiple assay types. Exposure route acronyms are as follows: IC, intracranial; OC, ocular; IN, intranasal; OR, oral; IT, intratracheal; IB, intrabronchial; AE, aerosol; IG, intragastric. Multi-route includes all exposures via three or more distinct locations (Table S2).

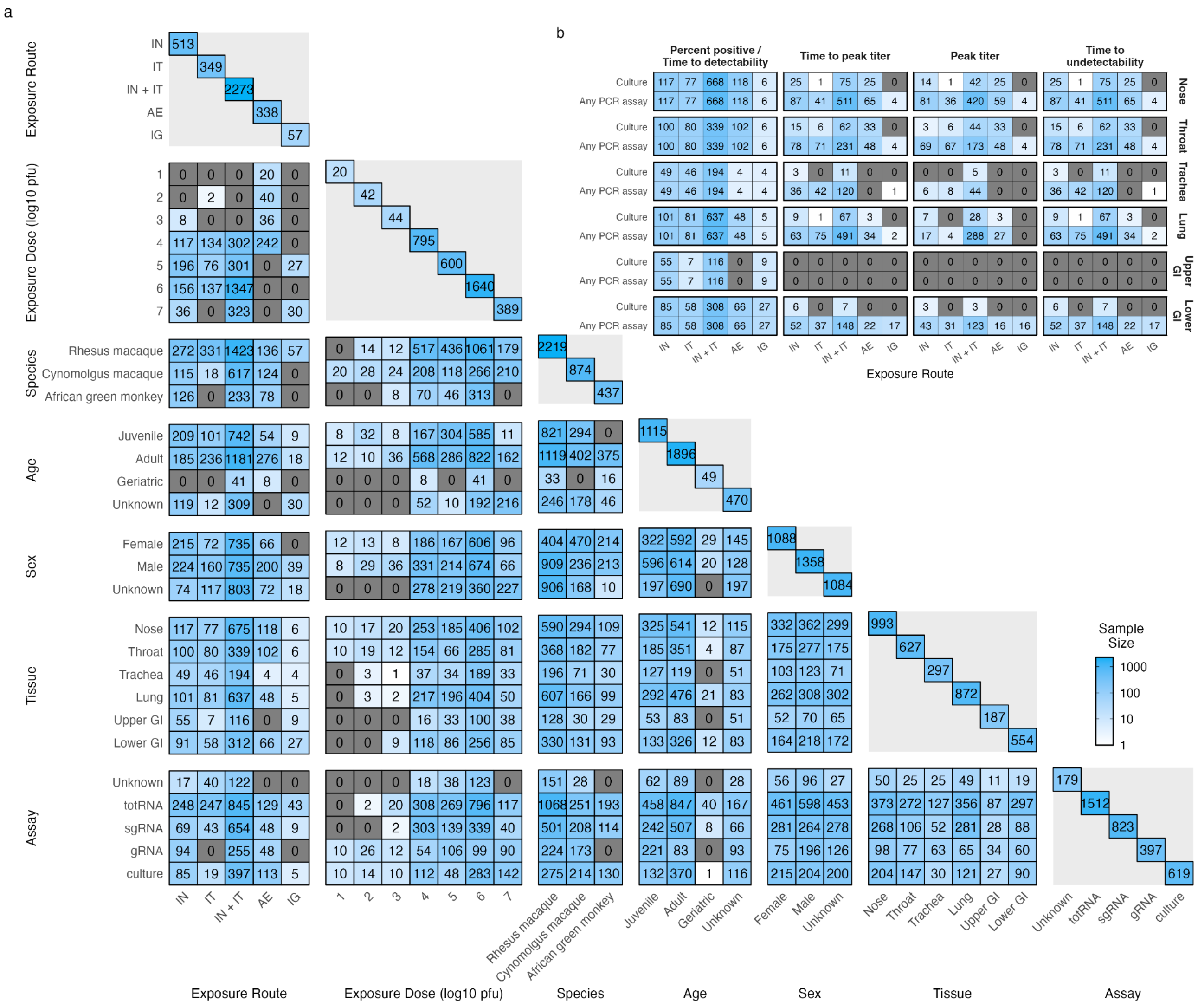


**Figure S3. Data distribution across cofactors and metrics for kinetic analysis.**

**a**, The text in each cell gives the maximum number of datapoints available for the pair of cofactors indicated by the row and column labels when fitting our survival model (not for the full database). Colors scale from white to blue based on the number of available datapoints. Grey cells correspond with combinations without available data. Entries along the diagonal give the datapoints for one cofactor, not a cofactor pair. **b**, The text in each cell gives the number of datapoints available for the metric indicated in the column label, for the assay(s) indicated on the lefthand side, for the tissue indicated on the right-hand side, and for the exposure route on the x-axis. Cells are colored as in panel a. Note that because of parameter sharing, our model can generate predictions for combinations of assay, route, tissue, and metric with zero entries.

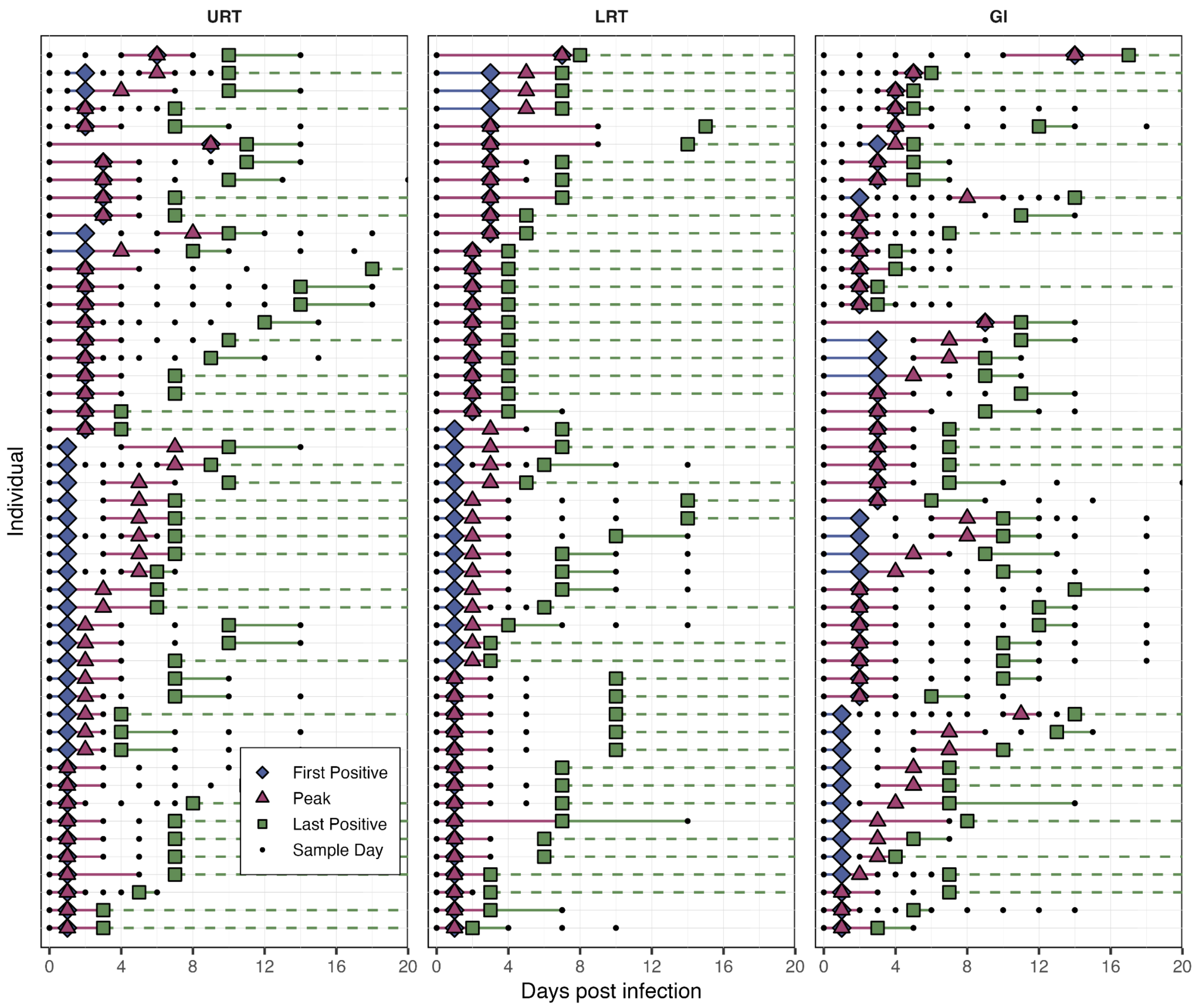


**Figure S4. Example censored data used for model fitting.**

The data presented here represent a small subset of the available time series from our database that we used to fit our survival model. Each row within each panel gives a unique individual's time series for the tissue location indicated in the panel label (URT: upper respiratory tract; LRT: lower respiratory tract; GI: gastrointestinal tract). Individuals are not consistent across the rows in different panels. Observed event times are shown with colored shapes (first positive, blue diamond; largest titer, purple triangle; last positive, green square). Each event time is censored and must occur within the time range indicated by the colored lines. Solid lines correspond to interval-censored observations. Dashed lines indicate the individual was positive on their final sample day, so their event is right censored. For visualization, only individuals with multiple positive samples whose observed peak time occurred on a different day from their lower bound on the time to undetectability were chosen as examples for this figure. All of the available data is displayed alongside the model fits in Figures S5-7.

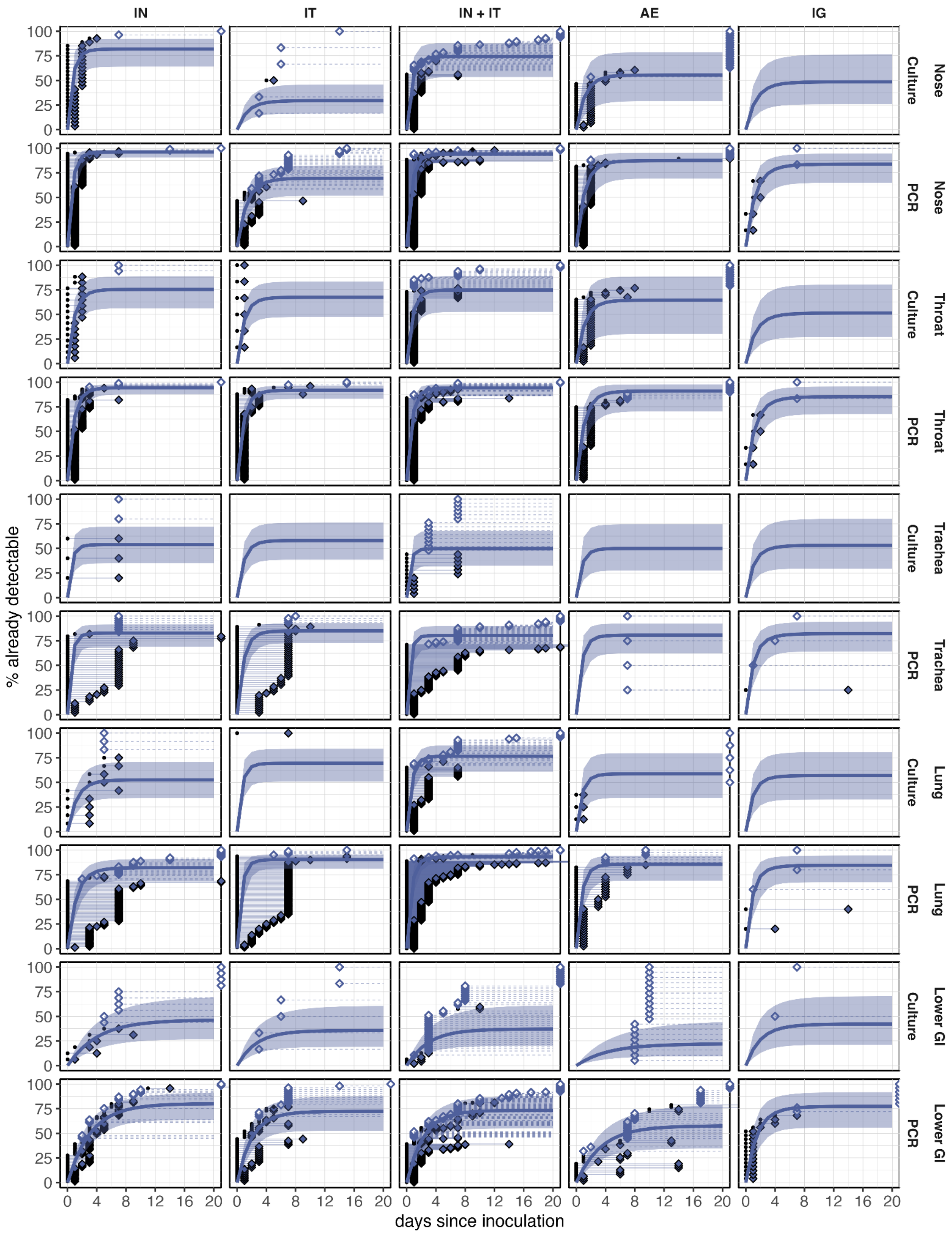


**Figure S5. Model fits for the probability of positivity and time to detectability.**

All panels are presented as in Extended Data Figure 1, except this figure includes PCR and culture assays, all tissue locations except the upper GI, and all exposure routes. Some observations occurred after day 20 post inoculation, which are plotted along the far right side of each panel. We were still able to generate predictions for panels without data because of information sharing in our model. Predictions are based on 500 samples of model posteriors.

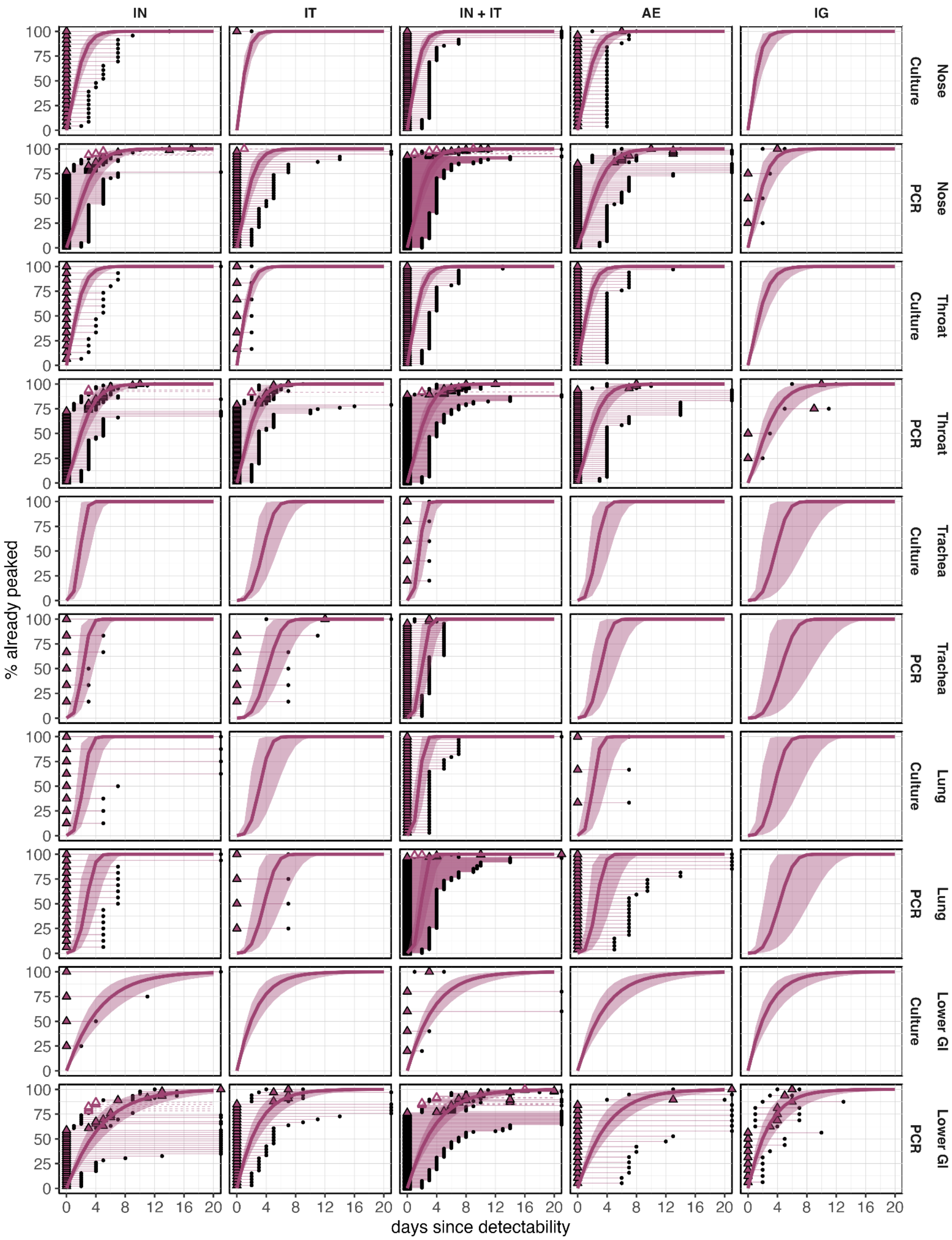


**Figure S6. Model fits for the time to peak titer.**

All panels are presented as in Extended Data Figure 1, except this figure includes both PCR and culture assays, all tissue locations except the upper GI, and all exposure routes. Some observations occurred after 20 days since detectability, which are plotted along the far right side of each panel. We were still able to generate predictions for panels without data because of information sharing in our model. Predictions are based on 500 samples of model posteriors.

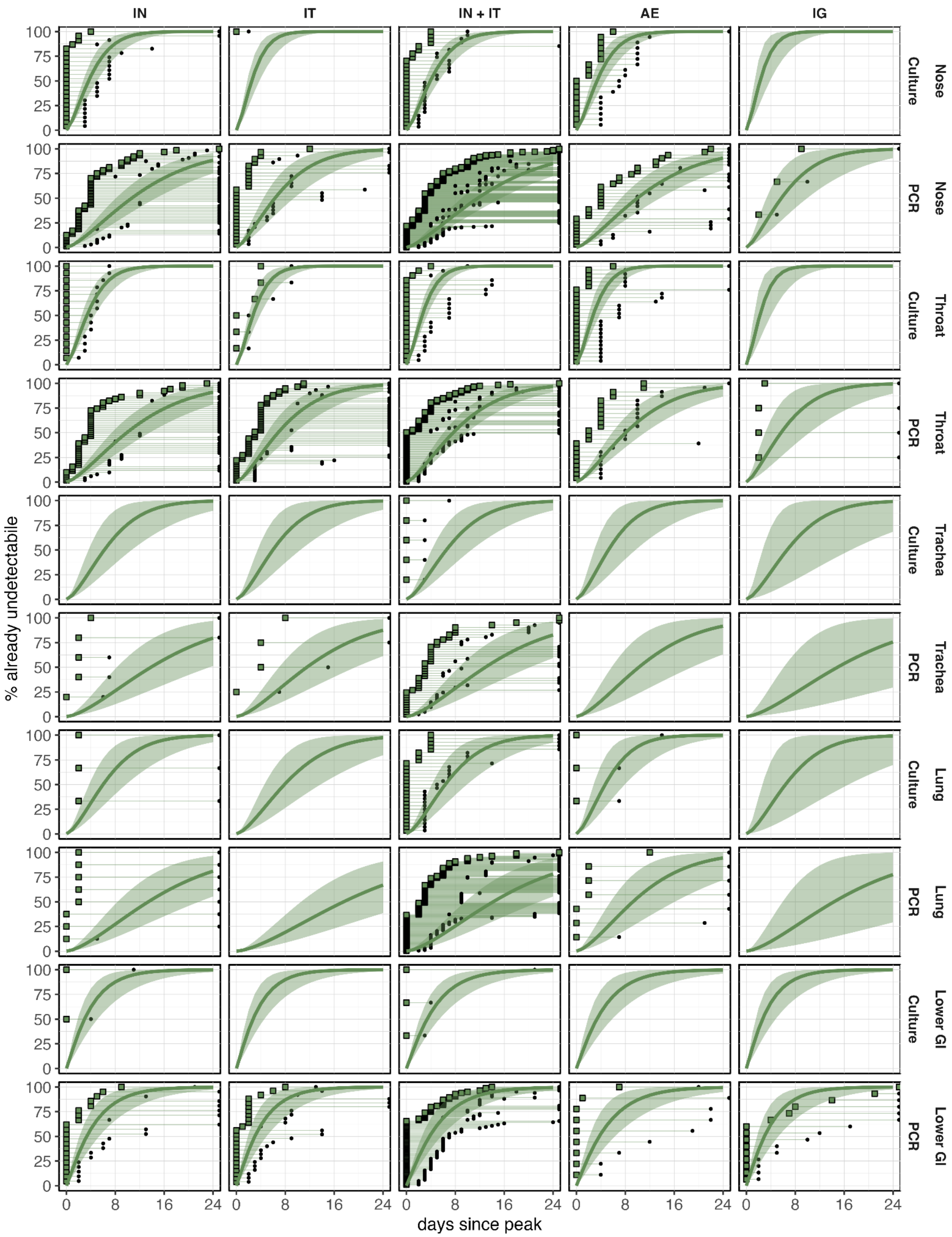


**Figure S7. Model fits for the time to undetectability.**

All panels are presented as in Extended Data Figure 1, except this figure includes both PCR and culture assays, all tissue locations except the upper GI, and all exposure routes. Some observations occurred after 24 days since peak titer, which are plotted along the far right side of each panel. We were still able to generate predictions for panels without data because of information sharing in our model. Predictions are based on 500 samples of model posteriors.

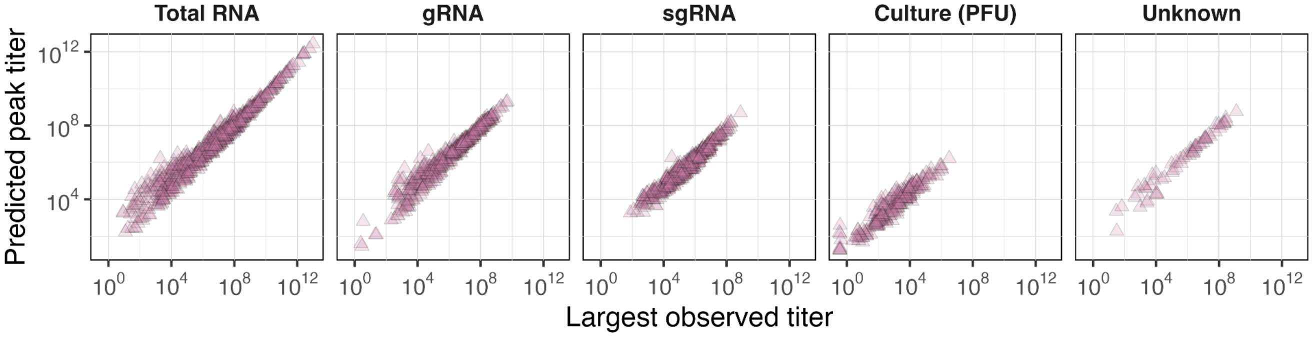


**Figure S8. Relationship between the largest observed titer and the mean true peak titer predicted by the model.** Results are stratified into panels based on the detection assay (labelled at the top). Each point is from an individual sampled in a given tissue. Predictions use all posterior samples.

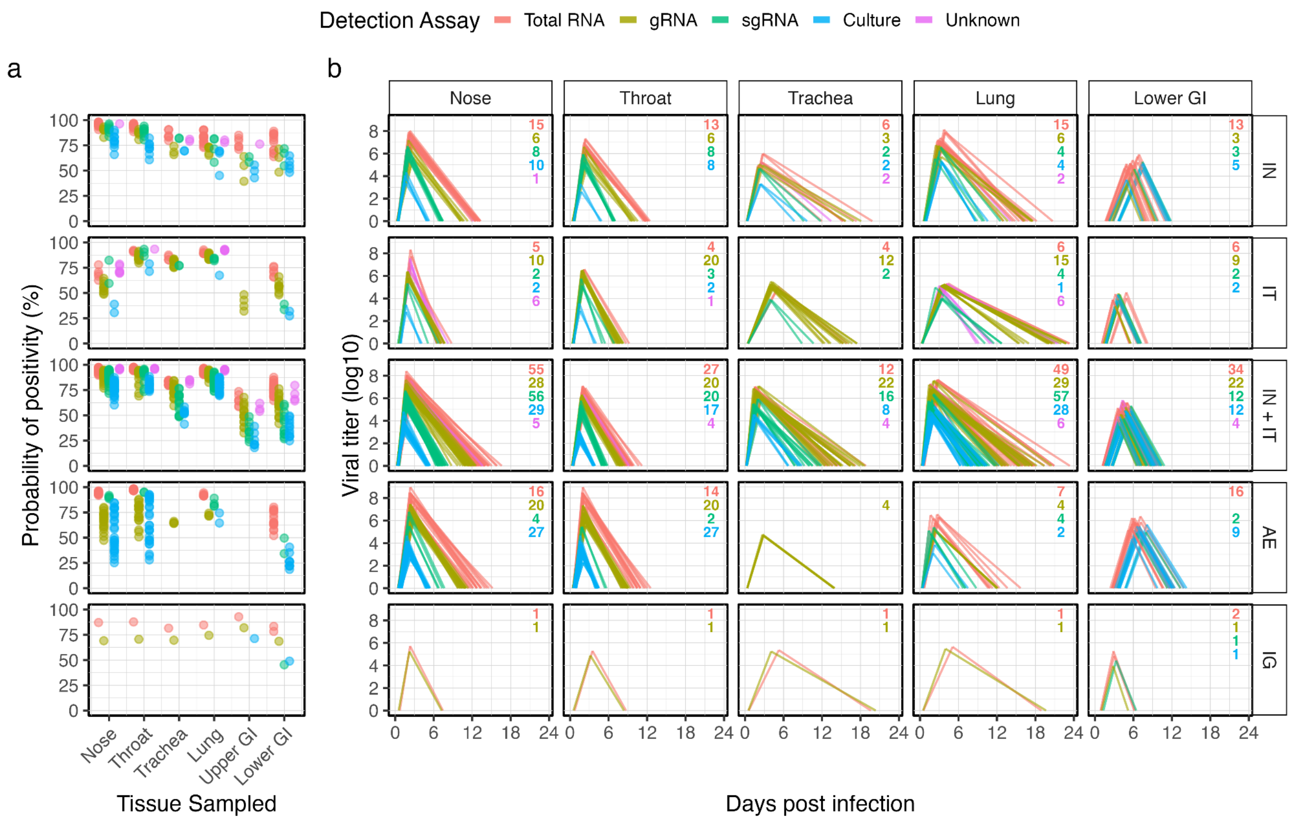


**Figure S9. Model predictions across the unique cofactor combinations used in each article.**

For each article, we generated median model predictions for all infection metrics for the exposure route(s), dose(s), age(s), sex(es), species, and assay(s) used in that article and only for the tissue(s) that article sampled. Predictions are stratified into panels based on the five main categories of route types (row labels), including the probability of positivity in panel **a** and infection trajectories in panel **b**. Sampled tissues are indicated by the x-axis label in panel a and by the column labels in panel b. Colors distinguish between the assays used for detection. Each point and line corresponds with the median prediction for one article. Annotated numbers in the top right corner of all panels in **b** give the number of lines plotted for each assay in the given panel. For articles where the PCR assay was unknown (i.e., purple lines), for the purposes of this figure we generated predictions for a total RNA assay given they are very commonly used.

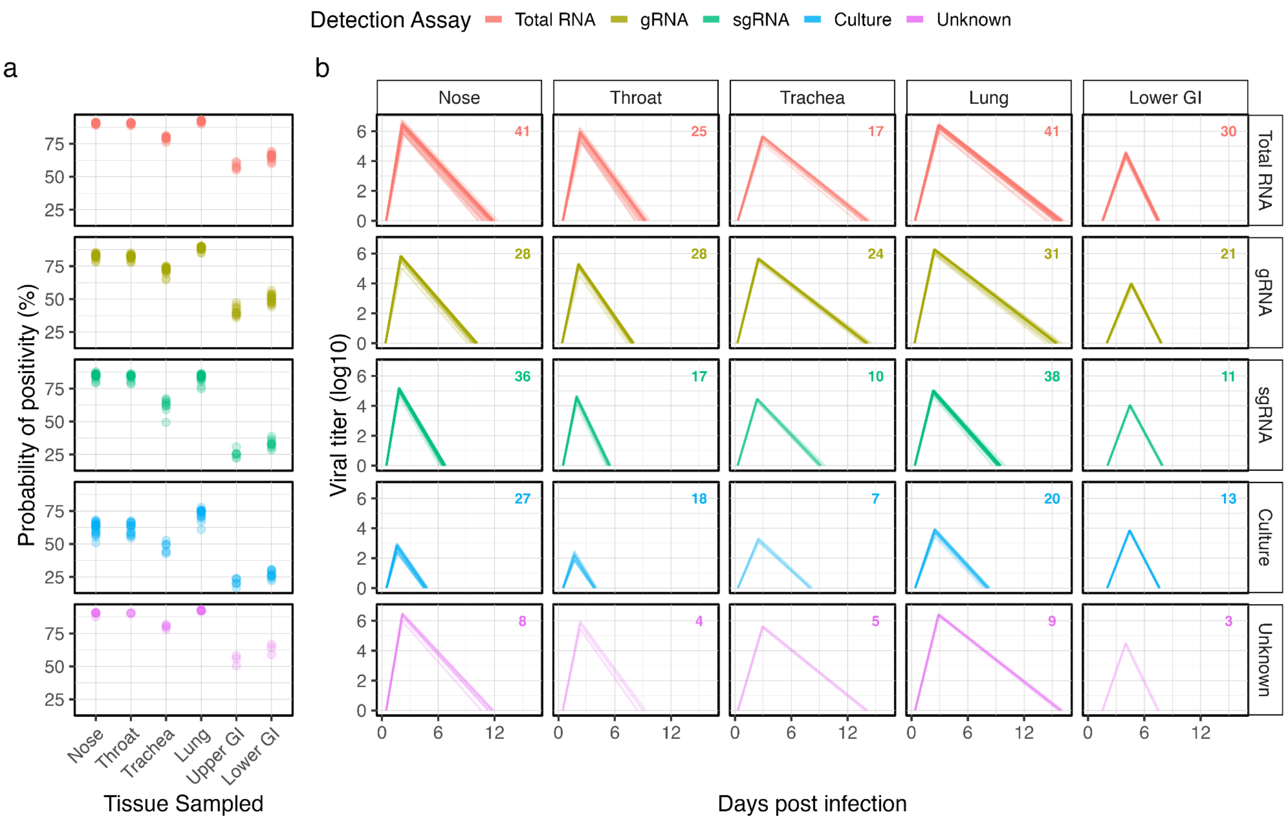


**Figure S10. Model predictions across articles for a fixed cofactor set.**

For each article, we generated median model predictions for all infection metrics under a fixed set of cofactors, namely for an adult, male rhesus macaque exposed via IN+IT to $10^4$ total pfu. We only generated predictions for the particular assay(s) used and tissues sampled in each article. By assuming all articles used the same experimental procedures, this isolates the inferred effects of unmodeled methodological variation among articles. Predictions are stratified into panels based on detection assay (row labels), including the probability of positivity in panel **a** and infection trajectories in panel **b**. Sampled tissues are indicated by the x-axis label in panel a and by the column labels in panel b. Colors distinguish between the assays used for detection. Each point and line corresponds with the median prediction for one sampled tissue, detection assay, and article. Annotated numbers in the top right corner of all panels in **b** give the number of lines plotted in the given panel, which also apply to the number of points plotted for each assay-tissue combination in panel a. For articles where the PCR assay was unknown (i.e., purple lines), we generated predictions for a total RNA assay given they are very commonly used.

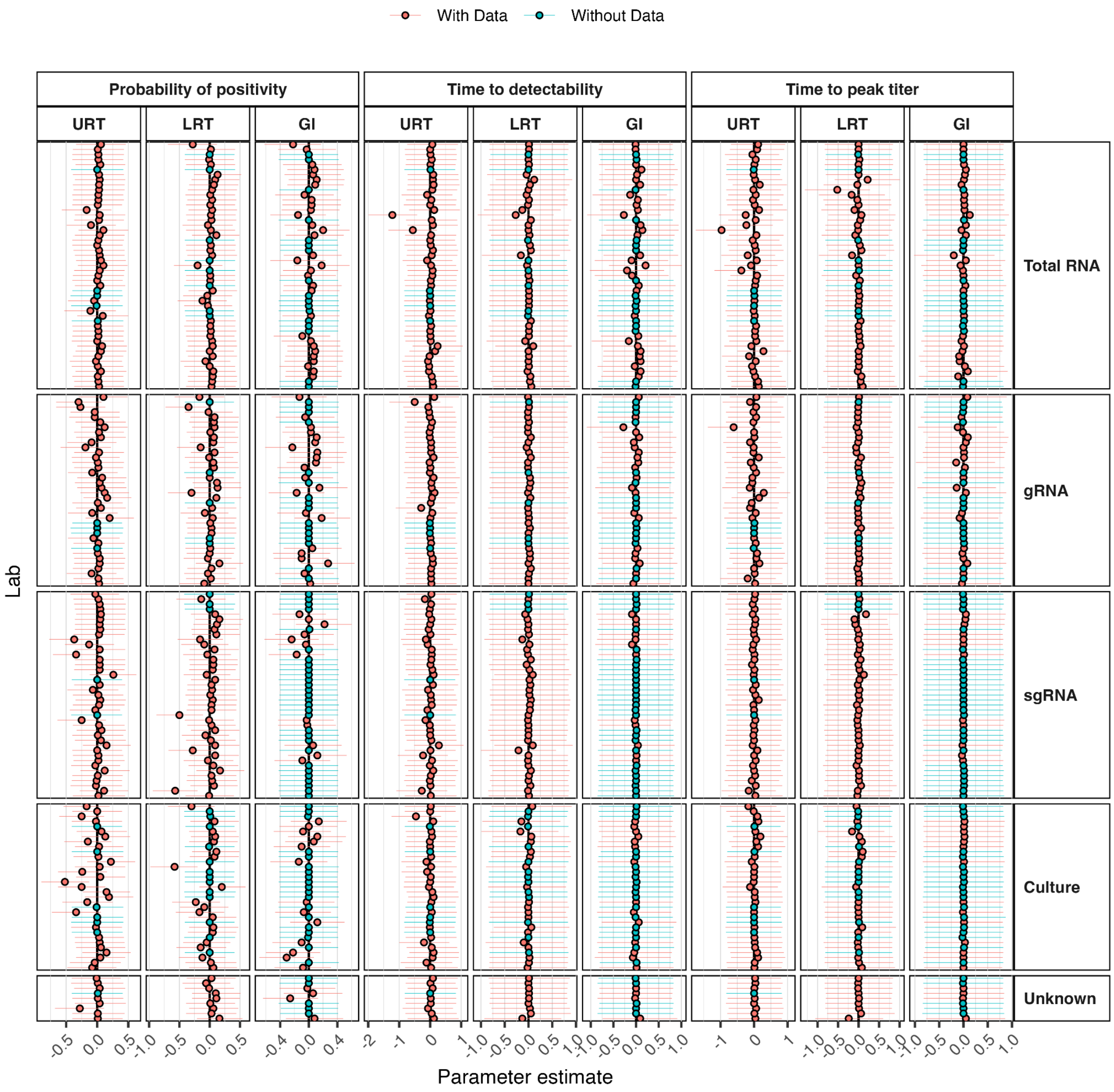


**Figure S11. Parameter estimates for article-specific effects on the probability of positivity, time to detectability, and time to peak titer.**

Each panel presents the median parameter estimates (points) and their 90% credible intervals (line segments) for each infection metric (outermost column labels), the three organ groups (innermost column label), and assay types (row label) for all corresponding articles (y axis). Colors indicate whether the particular article included samples from the indicated organ group. Notably, our model estimates the effects on all three organ groups even if the particular organ group was not sampled, though in these cases the parameter estimates are clearly centered at zero and we do not make article-specific predictions for these tissues in Figures S9 and S10. For some labs, the particular PCR assay was not reported, which are labeled as Unknown.

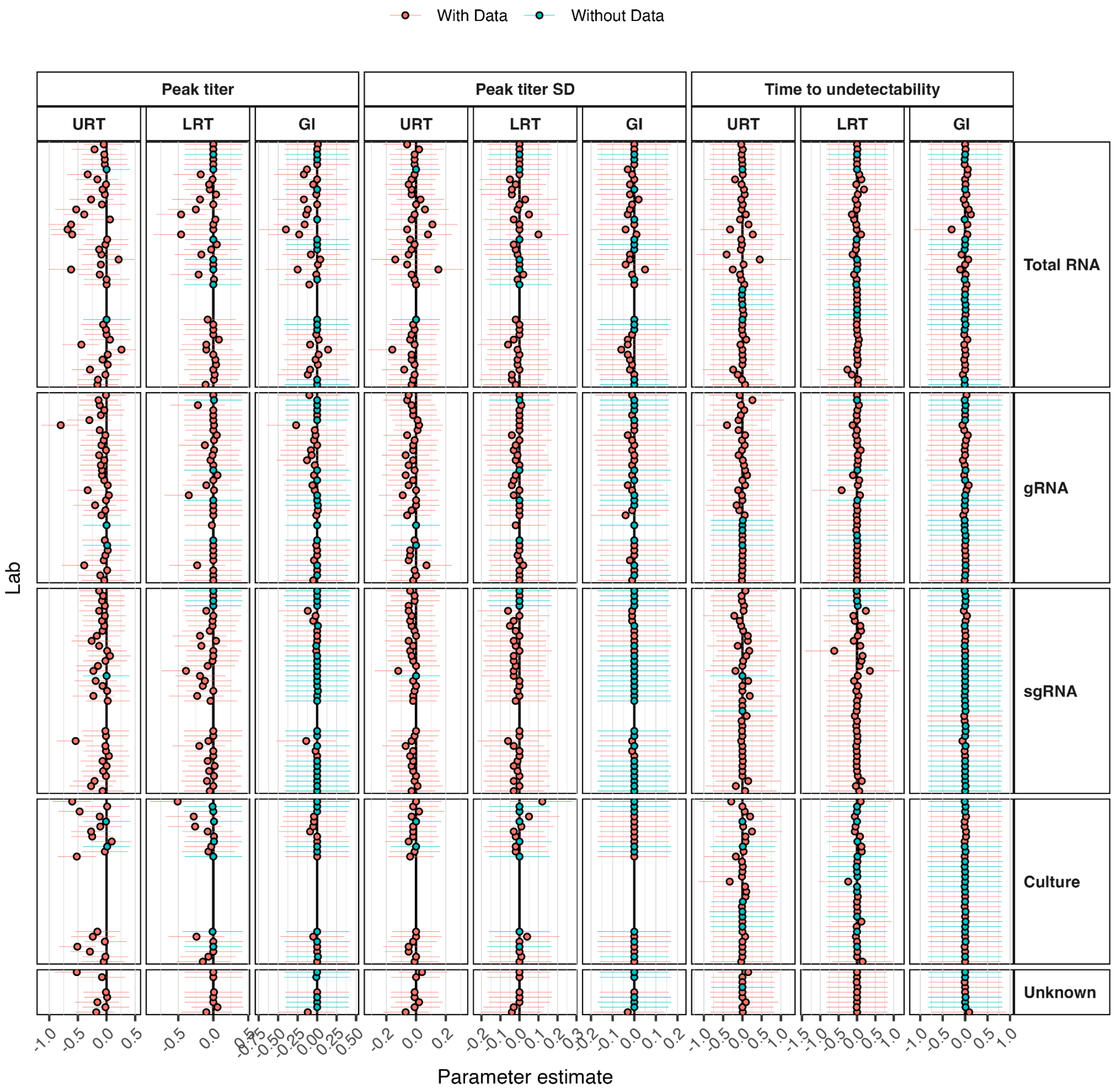


**Figure S12. Parameter estimates for article-specific effects on the peak titer, standard deviation around the peak titer, and time to undetectability.**

Each panel presents the median parameter estimates (points) and their 90% credible intervals (line segments) for each infection metric (outermost column labels), the three organ groups (innermost column label), and assay types (row label) for all corresponding articles (y axis). Some labs did not contribute information on peak titers and are empty spaces in this figure. Colors indicate whether the particular article included samples from the indicated organ group. Notably, our model estimates the effects on all three organ groups even if the particular organ group was not sampled, though in these cases the parameter estimates are clearly centered at zero and we do not make article-specific predictions for these tissues in Figures S9 and S10. For some labs, the particular PCR assay was not reported, which are labeled as Unknown.

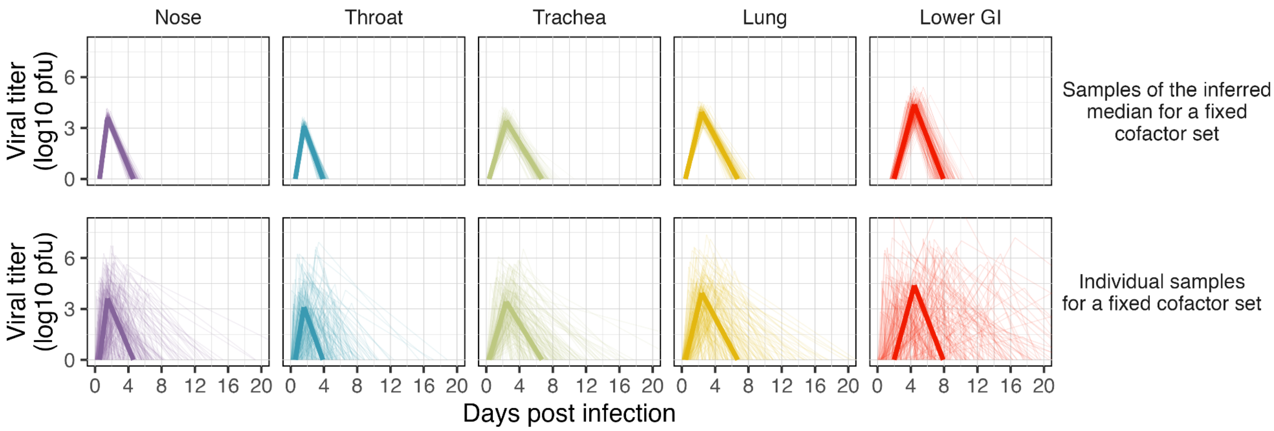


**Figure S13. Variation in individual trajectories.**

Panels distinguish between the tissue sampled (columns; also colors) and the type of predictions being displayed (rows). All panels show a total of 180 sampled trajectories with semi-transparent lines. Predictions are only shown for culture results. In the top row, each thin, semi-transparent line displays one trajectory sampled from the distribution of median trajectories, particularly for an adult, female rhesus macaque exposed via IN+IT inoculation to $10^4$ pfu. The thick, dark line displays the median of the distribution of medians. In the bottom row, each thin, semi-transparent line displays one individual-level trajectory for the same cofactor set as in the top row. These predictions were obtained by randomly sampling an event time from a Weibull distribution with a median that was sampled from the distribution of medians. The thick, dark line displays the median of the distribution of medians, as in the top row.

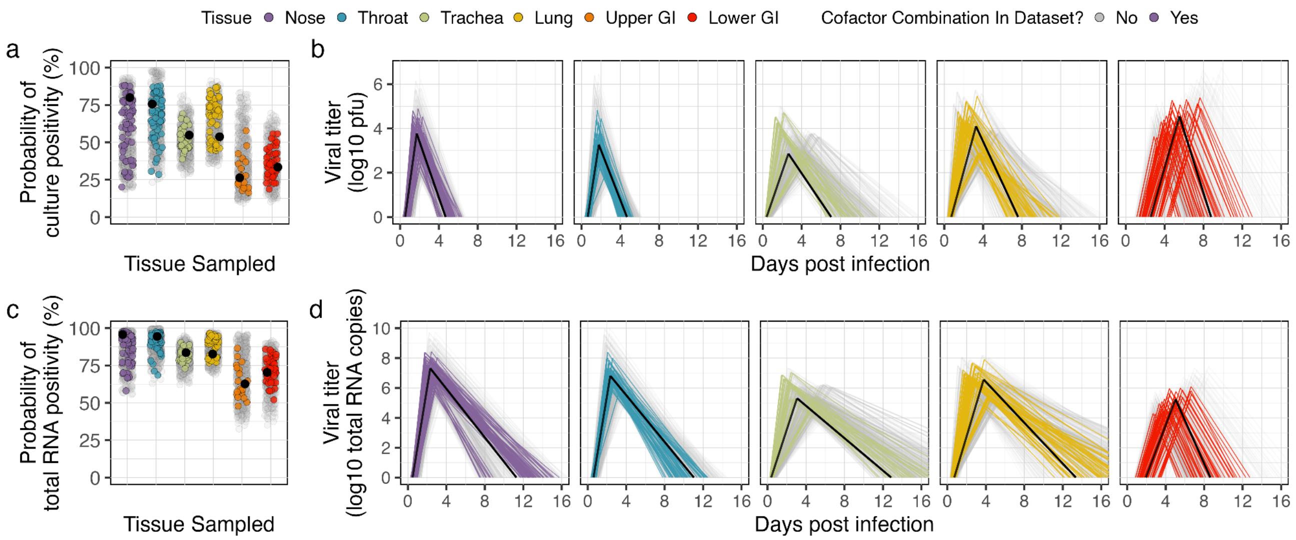


**Figure S14. Median infection trajectories across cofactor combinations.**

Colors in all panels distinguish between sampled tissues, as indicated in the legend at the top (purple: nose; blue: throat; green: trachea; yellow: lung; orange: upper GI; red: lower GI). Model predictions were generated and compared for the 630 possible combinations of age class (juvenile, adult, geriatric), sex (male, female), species (rhesus macaque, cynomolgus macaque, African green monkey), exposure route (five representative types: IN, IT, IN+IT, AE, IG), and exposure dose ($10^1$, $10^2$, $10^3$, $10^4$, $10^5$, $10^6$, and $10^7$ pfu). We refer to each unique combination of these cofactors as a 'cofactor combination' – e.g., female, adult rhesus macaques intranasally challenged with $10^4$ pfu are one possible cofactor combination (depicted with black dots and lines in all panels). Model predictions for cofactor combinations with data explicitly used in fitting for each tissue are displayed using tissue-specific colors. Note that for these classifications we took the floor of the log10 total pfu administered to each animal. The predictions for all other cofactor combinations extrapolated from the available data are shown in grey. For each of the 630 possible cofactor combinations, we compute their median probability of ever testing positive by culture assay (panel **a**) or by total RNA PCR assay (panel **c**) and their median infection trajectories as measured by culture assay (panel **b**) or by total RNA PCR assay (panel **d**) in each tissue. In panels **a** and **c**, each point is the median probability of positivity for one cofactor combination. In panels **b** and **d**, each line gives the median value for the median time to detectability, median time to peak titer, mean peak titer, and median times to undetectability for one cofactor combination. We computed each metric based on 200 samples of model predictions for each cofactor combination. Quantitative comparisons in peak titers across tissues should be made cautiously, given they often involve different sample types (e.g., nasal swabs vs. BAL; Table S3). See Figure S15 for quantification of the variance in each metric for each tissue.

| | | All cofactor combinations | | | | | | Only cofactor combinations in dataset | | | | | |
|---|---|---|---|---|---|---|---|---|---|---|---|---|---|
| Assay | Metric | Nose | Throat | Trachea | Lung | Upper GI | Lower GI | Nose | Throat | Trachea | Lung | Upper GI | Lower GI |
| Total RNA | Probability of positivity | 43 | 37 | 33 | 29 | 64 | 56 | 40 | 29 | 17 | 20 | 39 | 34 |
| Total RNA | Time to detectability | 1.2 | 1 | 0.8 | 1.1 | 1.6 | 4 | 0.7 | 0.6 | 0.4 | 0.6 | 0.8 | 2.7 |
| Total RNA | Time to peak titer | 2.3 | 2.7 | 5.4 | 5.5 | | 6.1 | 1.6 | 1.4 | 3.7 | 3.2 | | 4 |
| Total RNA | Peak titer | 4.1 | 4.7 | 3.6 | 4.1 | | 4.4 | 2.4 | 2.7 | 2.5 | 3 | | 2.3 |
| Total RNA | Time to undetectability | 9.7 | 6.9 | 18 | 24.5 | | 13.1 | 8.3 | 4.7 | 9.4 | 12.8 | | 7.6 |
| Culture | Probability of positivity | 75 | 75 | 53 | 57 | 74 | 59 | 68 | 59 | 30 | 42 | 41 | 37 |
| Culture | Time to detectability | 1.3 | 1.1 | 0.8 | 1.1 | 2.1 | 5.4 | 0.8 | 0.6 | 0.4 | 0.6 | 1.1 | 3.5 |
| Culture | Time to peak titer | 1.9 | 2 | 4.6 | 4.7 | | 7.4 | 1.1 | 1 | 3.1 | 2.7 | | 4.8 |
| Culture | Peak titer | 4.3 | 5.1 | 3.4 | 3.9 | | 4.4 | 2.7 | 3 | 2.4 | 2.8 | | 2.3 |
| Culture | Time to undetectability | 3 | 2.5 | 10.3 | 12.1 | | 13.5 | 2.5 | 1.6 | 5.2 | 5.5 | | 7.9 |

Metric / Tissue Sampled

**Figure S15. Maximum differences in each metric across cofactor combinations.**

This figure quantifies the variance in predictions across cofactor combinations that are depicted visually in Figure S14. Each cell gives the largest difference in the median prediction for the associated metric (y axis) in the sampled tissue (x axis) either among all possible cofactor combinations (left panels) or among only the cofactor combinations included in the data used for fitting (right panels). These values are further stratified according to the assay type, namely total RNA PCR (top panels) or culture (plaque assay; bottom panels). Units for each row correspond to the metric, so the probability of positivity displays the difference in percentage, temporal metrics display differences in days, peak titer displays differences in either log10 total RNA copies (top panels) or log10 pfu (bottom panels). Cells are lightly colored according to the sampled tissue, which are also labelled along the x axis.

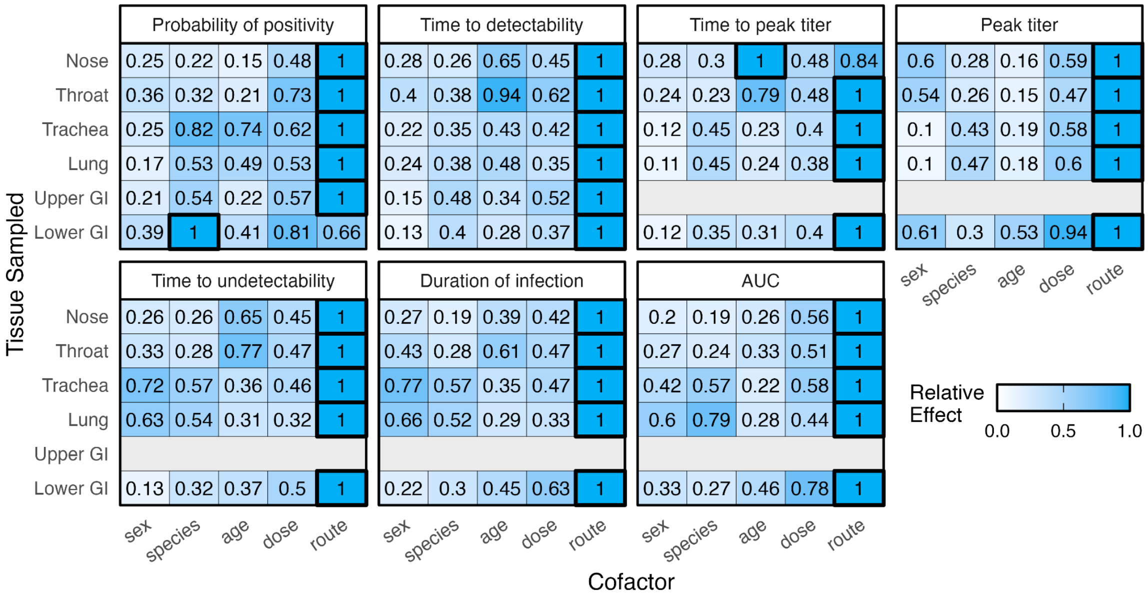


**Figure S16. Relative effects of each cofactor on infection metrics.**

Results were generated and plotted as described in Figure 3, but the text labels provide the relative effects of each cofactor on the indicated metric instead of the mean differences. The cofactor with a '1' and the darkest blue had the largest mean effect. The intensity of the color for all other cofactors are scaled according to their relative effect against the largest mean effect. These results were generated for culture assays.

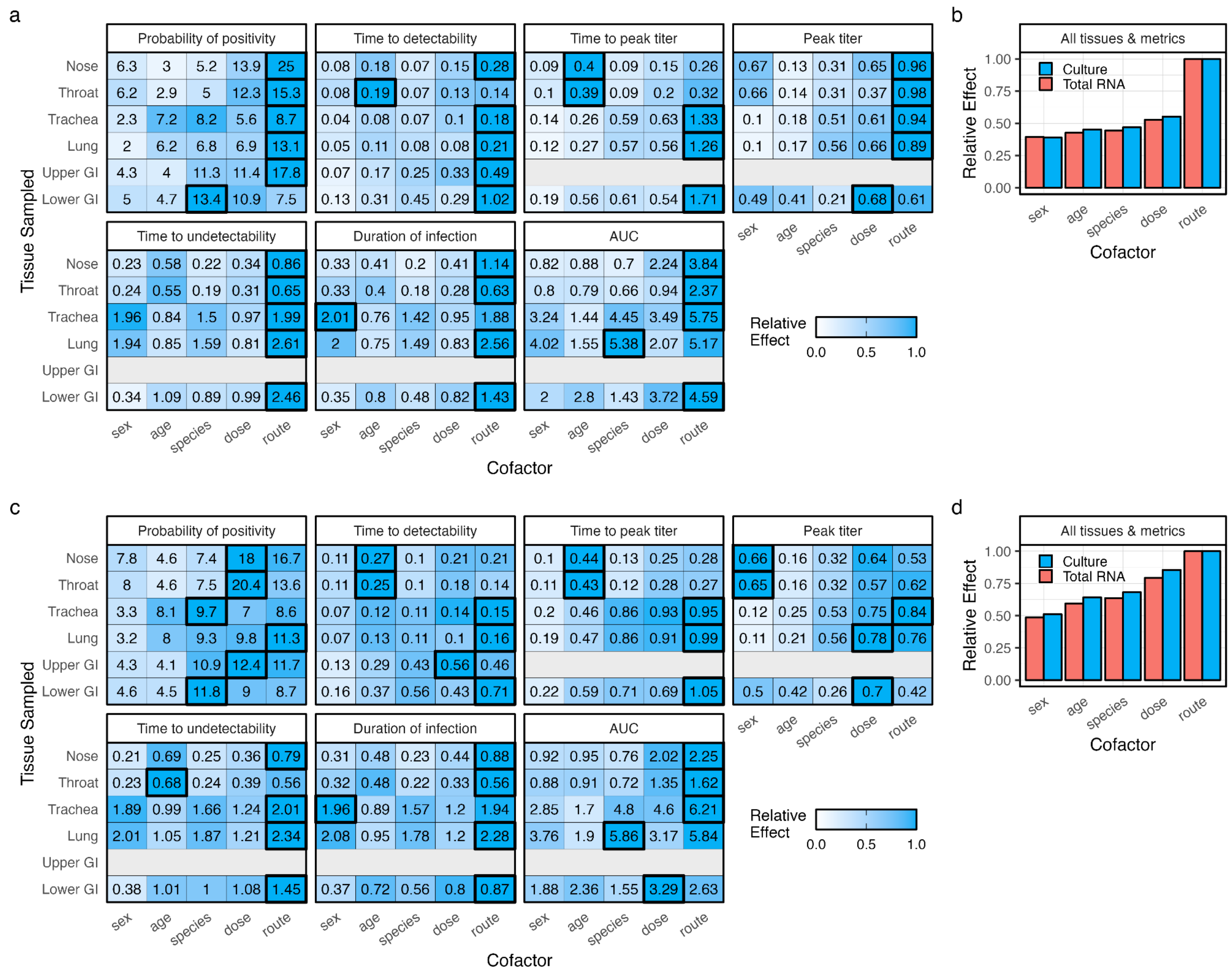

**Figure S17. Sensitivity of rankings to summary statistics and dose comparisons.**

**a-b,** Rankings of cofactor effects as in Figure 3, except calculated using the median differences. **c-d,** Rankings of relative cofactor effects as in Figure 3, except allowing dose comparisons between $10^1$ and $10^4$ pfu in addition to $10^4$ and $10^7$ pfu. Note that aerosol inoculation is the only exposure route with data in our dataset below $10^{3.5}$ pfu, so these analyses required generating predictions for a much lower dose ($10^1$ pfu) than has been observed for IN, IT, IG, or IN+IT exposures. Panels b and d also compare the results when analyses are conducted based on total RNA detection (red) instead of culture positivity (blue).

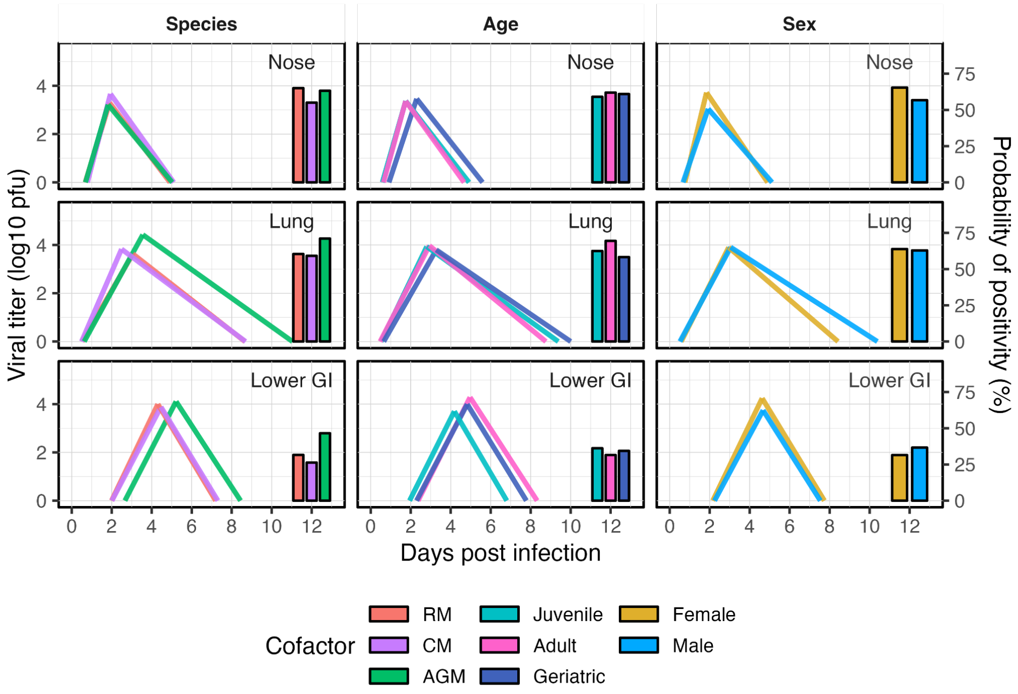


**Figure S18. Effects of demographic factors on within-host kinetics across exposure routes.** Each row of panels displays model predictions for one tissue (top: nose; middle: lung; bottom: lower GI), and each column stratifies predictions based on the cofactor indicated in the column label (e.g., Species distinguishes results among rhesus macaques [RM], cynomolgus macaques [CM], and African green monkeys [AGM]). Colors within a column of panels distinguish among the constituent cofactors, as shown in the legend. Lines are the median infection trajectories when averaging across all exposure routes with a dose of $10^4$ pfu. Bar plots centered at 12 days post infection display the median probability of positivity, which corresponds with the y axis label on the right-hand side.

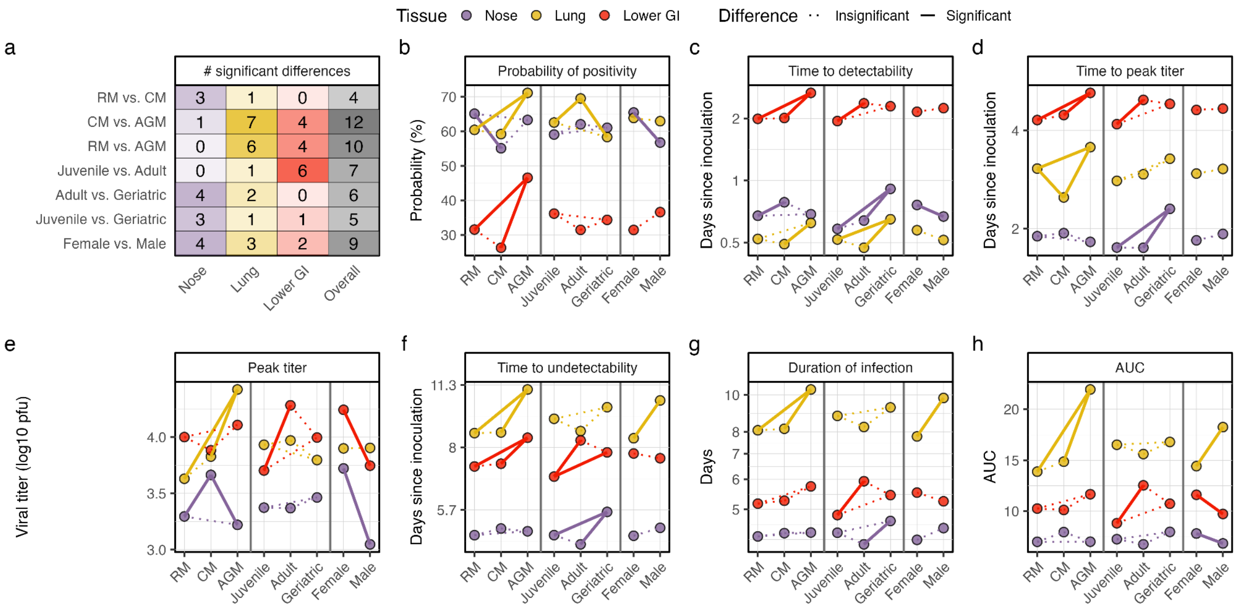


**Figure S19. Effects of demographic factors on infection metrics.**

Colors in all panels denote the tissue being sampled (purple: nose; yellow: lung; red: lower GI). All results were generated for viral culture. Species acronyms are: RM, rhesus macaque; CM, cynomolgus macaque; AGM, African green monkey. **a**, The number of differences that are significant in each location (columns) between the demographic groups labeled on the rows. The 'Overall' column sums the values for each tissue to give the total number of significant differences for those demographic groups. There were 6 possible differences for each cell, except for the overall column where there were 18 possible differences. The intensity of the color scales with the percent of differences that were significant. **b**, The median probability of positivity for each demographic group indicated along the x axis. All estimates were based on predictions generated for all routes, one dose ($10^4$ pfu), and all possible demographic factors except the one being varied (e.g., the RM estimates include predictions for all ages and sexes but not for CM or AGM). Solid, colored lines connecting the dots indicate the corresponding difference was significant. Dotted lines indicate the difference was insignificant. Other panels are similar except they display the median predictions of: **c**, the median time to detectability; **d**, the median time to peak titer; **e**, the mean peak titer, **f**, the time to undetectability; **f**, the duration of infection; and, **g**, AUC.

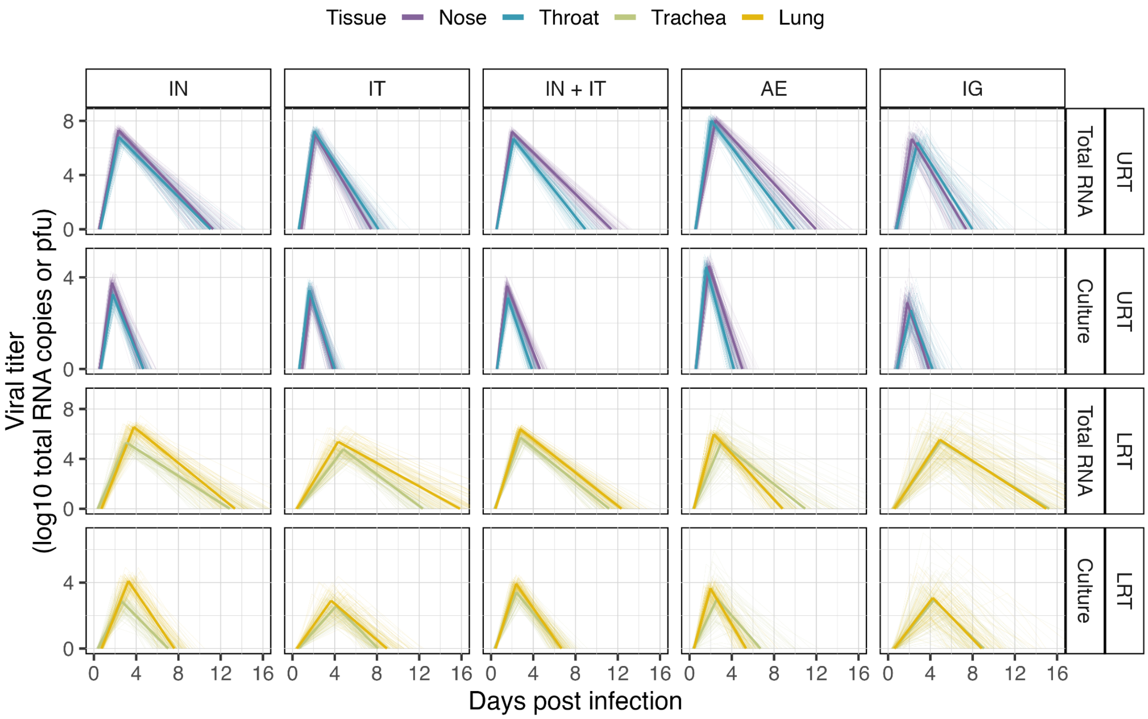


**Figure S20. Similarities in trajectories within respiratory tissues.**

Rows stratify results into upper respiratory tract (URT) vs. lower respiratory tract (LRT) tissues in addition to detection via total RNA PCR vs. viral culture (plaque assay). Column labels denote the exposure route. Colors distinguish between tissues as in the legend (purple: nose; blue: throat; green: trachea; yellow: lung). Thick, opaque lines give the median trajectory. Thin, transparent lines give samples of the median trajectory.

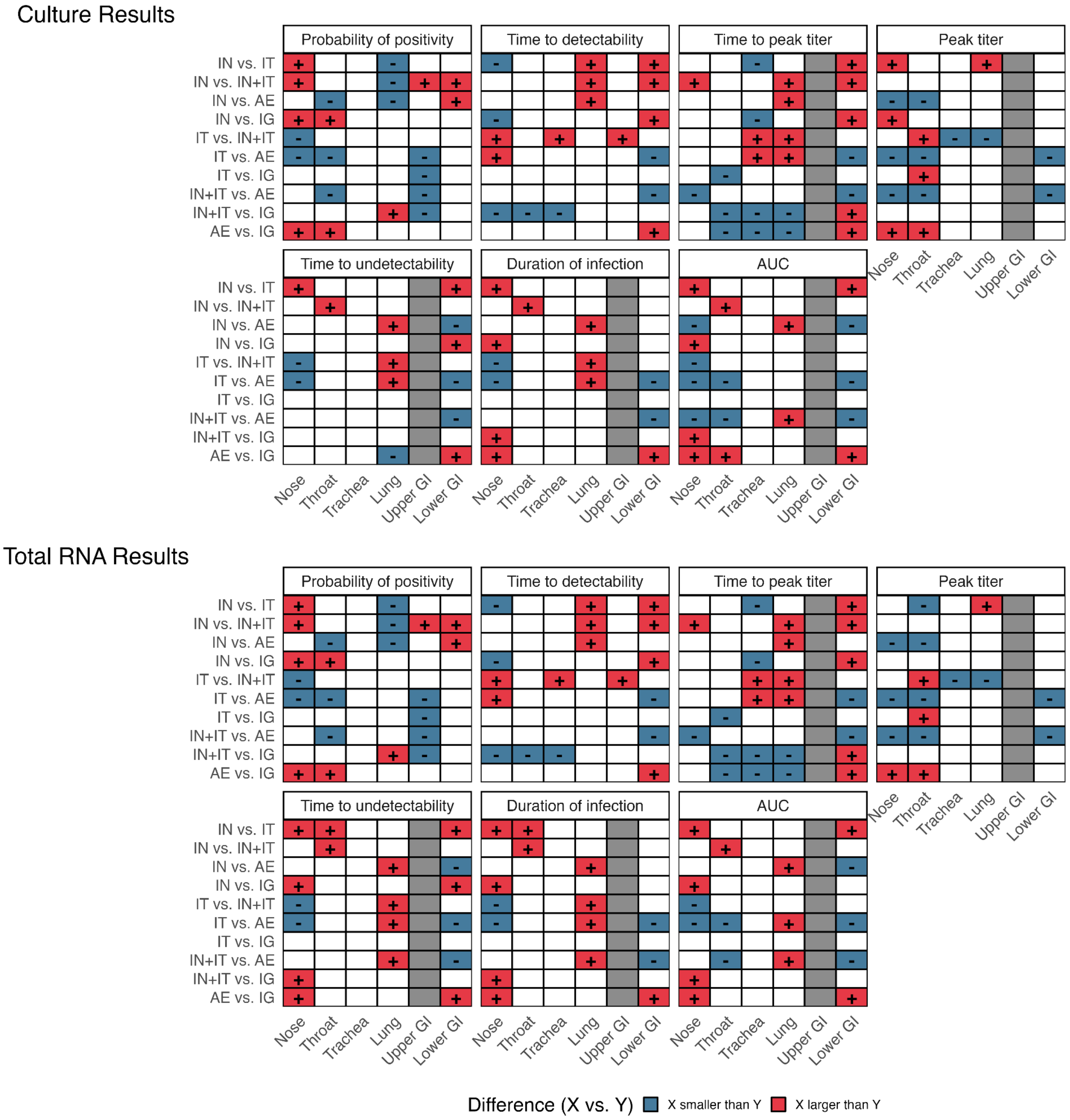


**Figure S21. Significant differences in all metrics among routes and in all tissues.**

Results are stratified by metric (panel label), tissue sampled (x axis label), route comparison (y axis label), and the detection assay (culture: top two rows; total RNA: bottom two rows). Cells indicate whether the difference in metric between route X and Y (labeled X vs. Y) is significant. A "+" with a red cell indicates that the metric for route X is significantly larger than for route Y. A "-" with a blue cell indicates that the metric for route X is significantly smaller than for route Y. A white cell indicates the difference is not significant. For example, the probability of positivity in the nose is significantly larger for IN than IT exposures. Grey cells indicate no comparison was possible. These results integrate all age classes, sexes, and species. Figure S22 shows counts of how many metrics are significantly different among each pair of exposure routes.

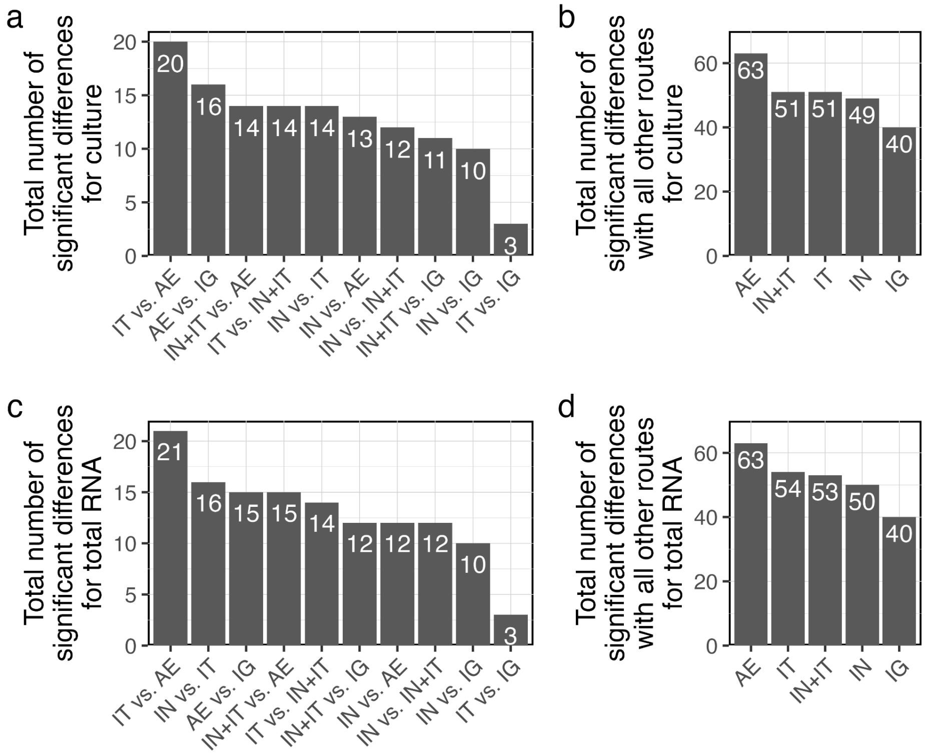


**Figure S22. The total number of significant differences between exposure routes.**

Culture results are shown in panels a and b, and total RNA results are shown in panels c and d. Panels a and c give the total number of significant differences between all possible pairs of exposure routes, integrating all of the metrics and tissues displayed in Figure S21. For example, IT and AE exposure differ significantly for 20 metrics based on culture results. The maximum possible number of significant differences among two pairs of exposure routes for a given detection assay is 37. Panels b and d give the total number of significant differences that each exposure route has with all other exposure routes combined, summing the differences counted in panels a and c. For example, for culture results, AE differs significantly from all other exposure routes for a total of 63 metrics.

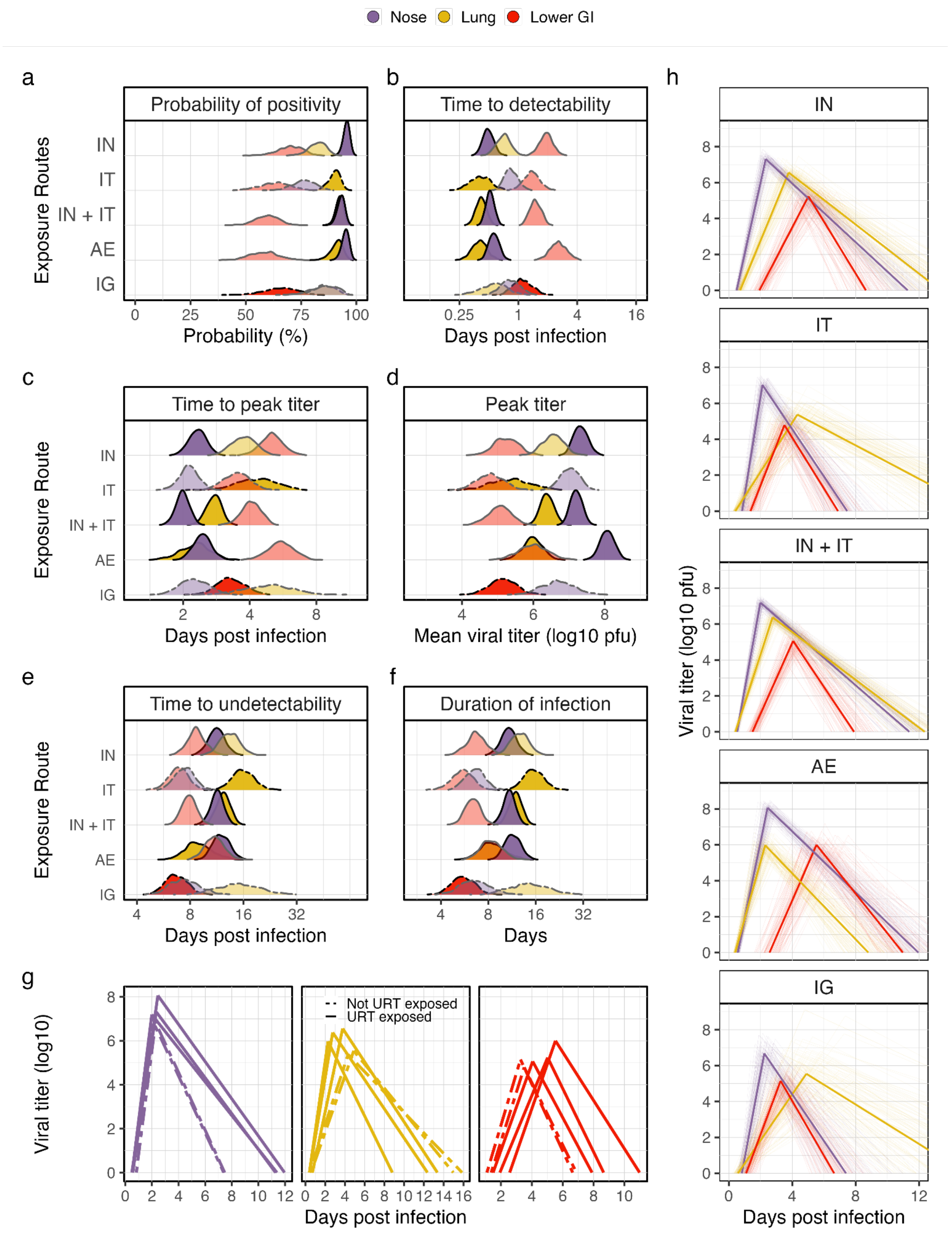


**Figure S23. Route effects visualized for total RNA.**

All panels are presented as in Figure 3, except that this figure displays results for total RNA assays.

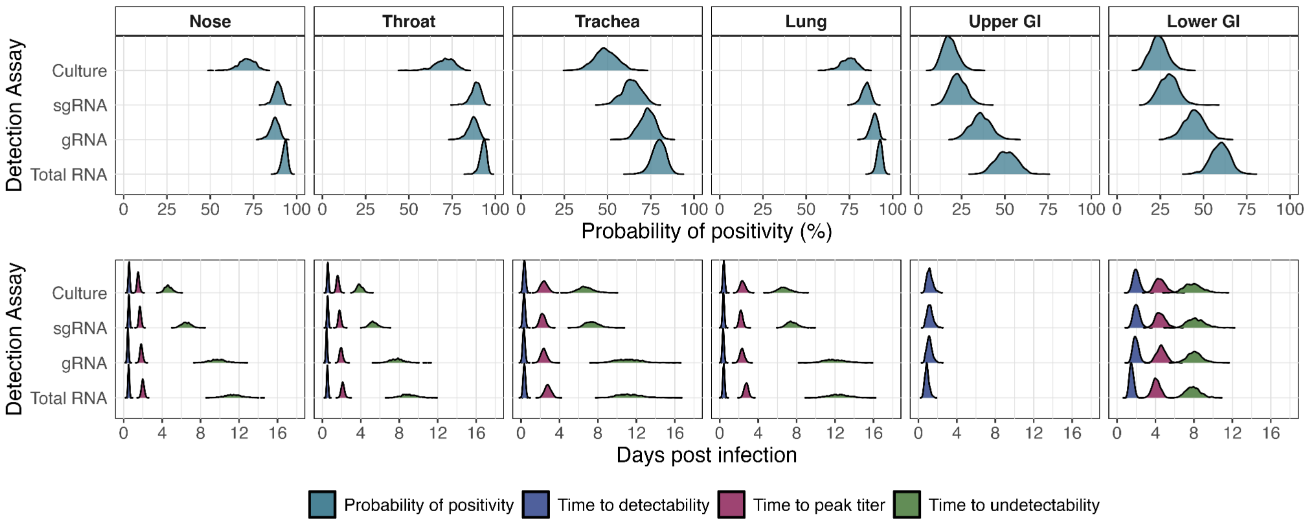


**Figure S24. Comparison of predictions across assay types.**

All panels present predictions for a female, adult rhesus macaque exposed to $10^4$ pfu by IN+IT inoculation. Predictions are stratified into panels by the tissue sampled (columns) and by the set of metrics (probability of positivity in the top row; all temporal metrics in the bottom row). Colors distinguish between the metrics as indicated in the legend. The y-axis in all panels indicate the assay.

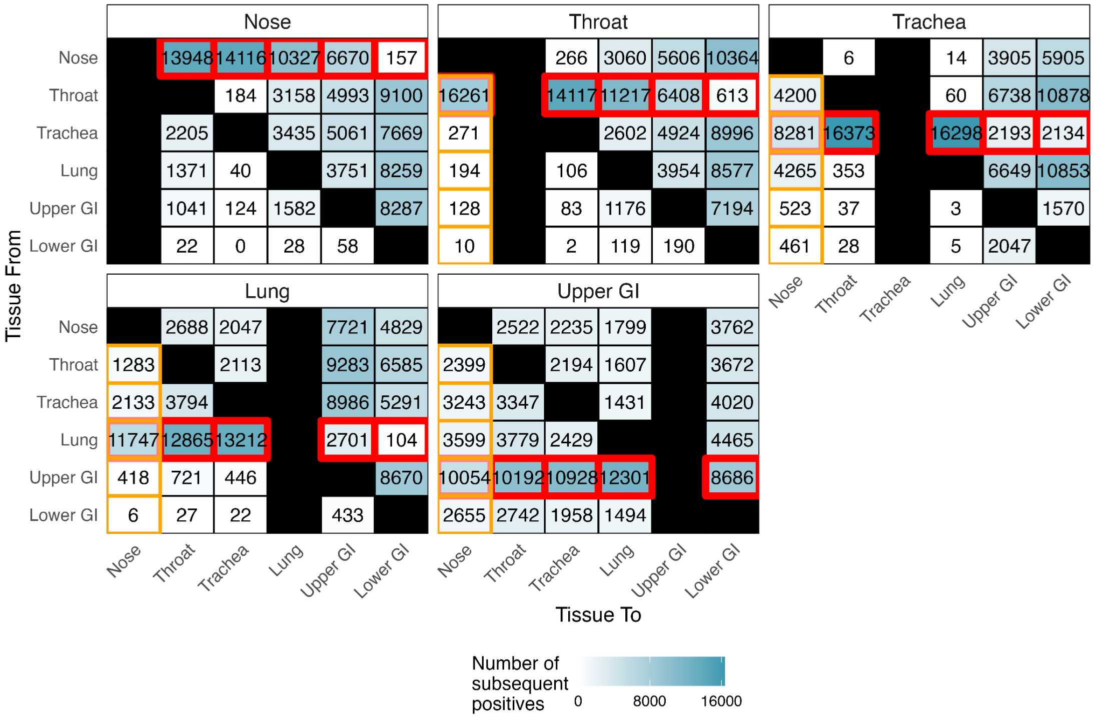


**Figure S25. Adjacency matrices used to generate tissue connectivity structure.**

Each panel corresponds with predictions generated following single-tissue inoculation into the tissue indicated by the panel label (see Methods). The text in each cell indicates the number of predictions where the tissue on the x axis becomes detectable after the tissue on the y axis, where the fill color of the cell scales with that number. The diagonals are colored black with no text because a tissue cannot become detectable after it has already become detectable. The columns corresponding with the inoculated tissue in the panel label are also colored black because they by assumption become detectable before all other tissues. Red squares within each panel indicate the entries used to characterize the outflows from each tissue indicated in the panel label, represented in Fig. 4a. Orange squares in all panels except ‘Nose’ shows the columns averaged to obtain the inflows into the Nose, represented in the ‘Nose’ column in Fig. 4b. Similar calculations were made to generate the columns for the other tissues.

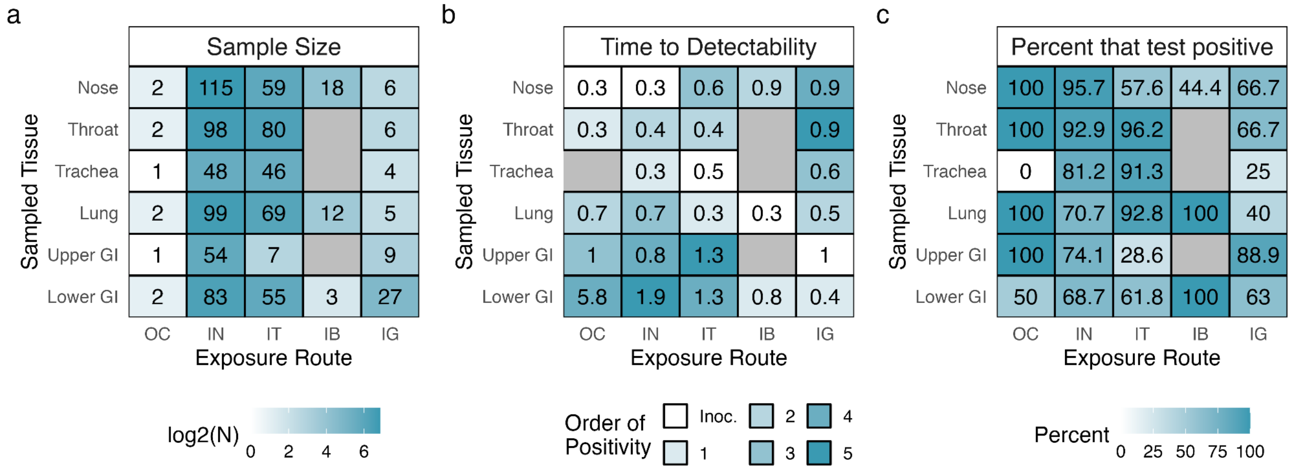


**Figure S26. Model predictions for the probability and time to detectability of individuals exposed via single-route inoculations in the dataset.**

These analyses include all individuals in the dataset that were exposed by single-route inoculations (OC: ocular; IN: intranasal; IT: intratracheal; IB: intrabronchial; IG: intragastric), regardless of the assay used for detection. In all panels, grey cells indicate there was no available data. **a**, The sample sizes available for each tissue (row label) and exposure route (column label). Cell intensity scales with the log2 sample size, and raw sample sizes are indicated by the text label. **b**, The average inferred time to detectability for each tissue (row label) and exposure route (column label). For each individual, we calculated the median inferred times to detectability in all tissues for which they had available data, using all model posteriors. Of these median times, we then computed the average across individuals for each tissue and route, which is indicated by the text label in each cell. Cell intensity indicates the relative ordering of times to detectability, with light blue indicating the earliest detection and darkest blue indicated the latest detection. White cells indicate the tissues that are intentionally inoculated by that exposure route. **c**, The percent of individuals that were ever observed to test positive in each tissue (row label) for each exposure route (column label). Cell intensity scales with the percent of individuals, which is also indicated with the text label.

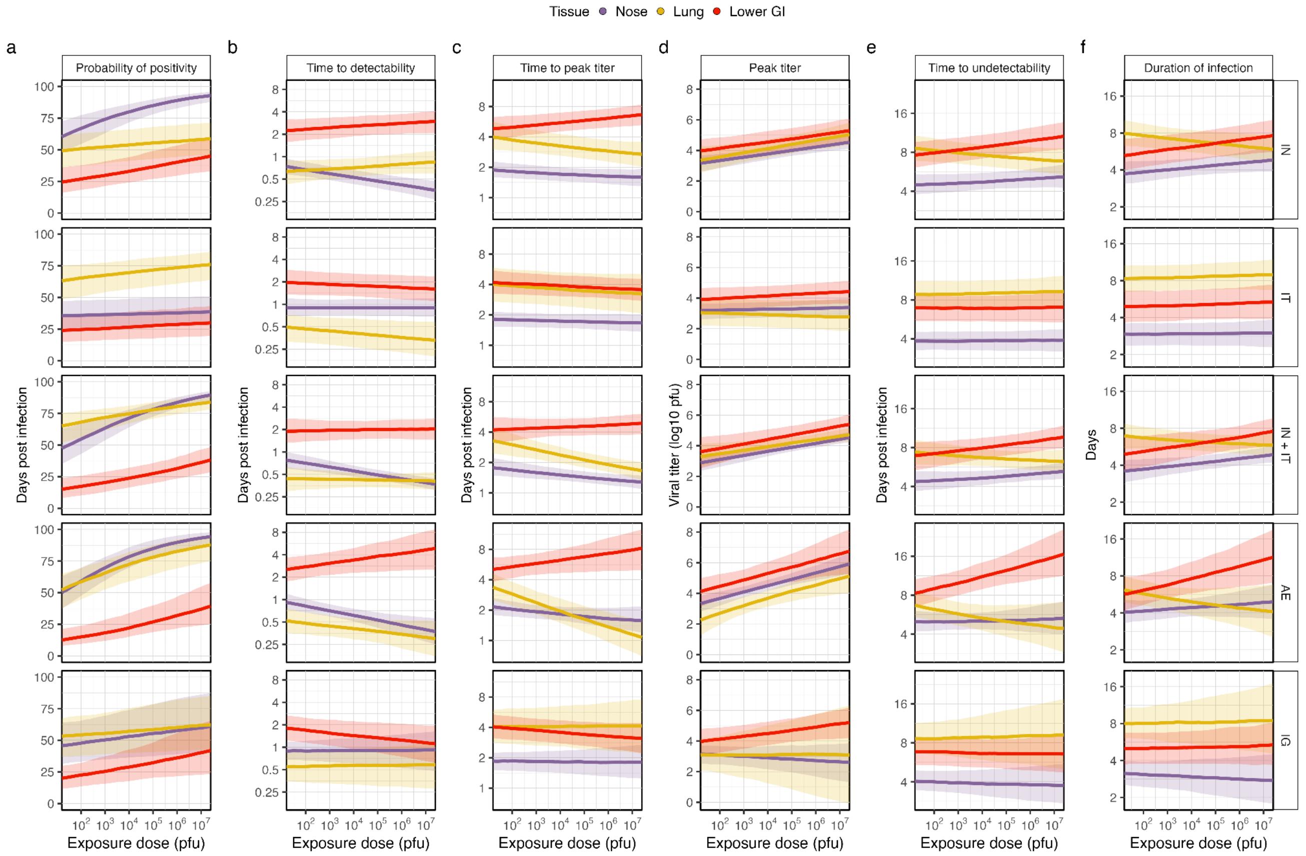


**Figure S27. Dose effects on culture metrics for all exposure routes.**

Visualization techniques are the same as for Extended Data Figure 4, except this figure includes predictions for all exposure routes and for the probability of positivity. Each row shows the results for one exposure route, as indicated with facet labels along the righthand side of the whole figure.

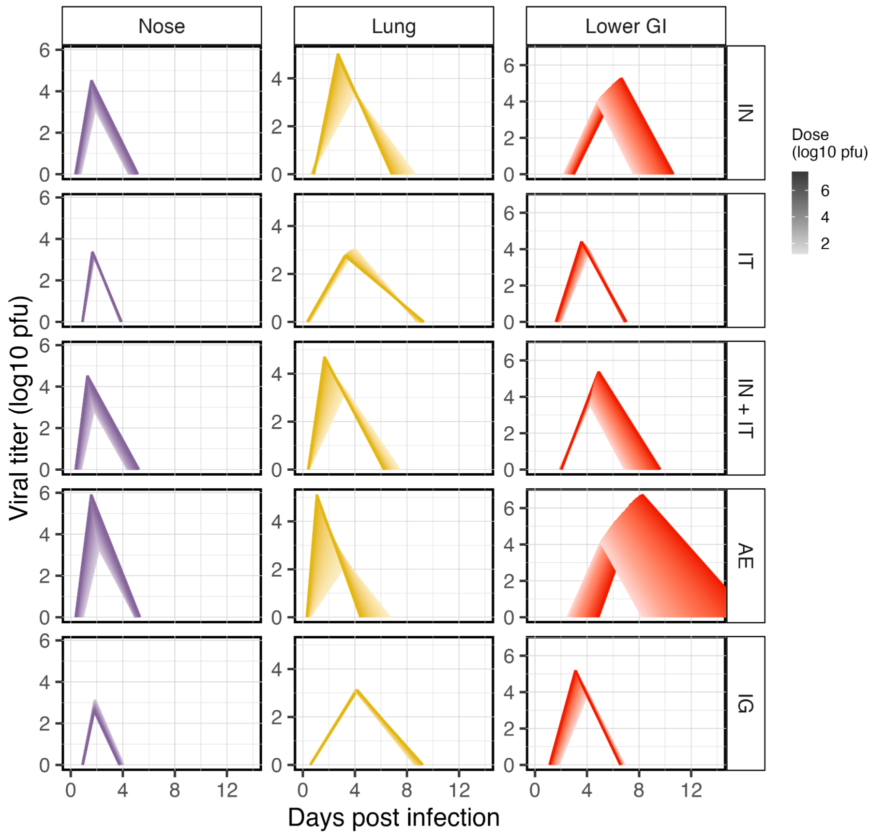


**Figure S28. Dose effects on culture trajectories for all exposure routes.**

Results are presented as in Figure 5d with lighter color intensity corresponding to smaller exposure doses. Different exposure routes are shown in distinct rows. Colors correspond with the tissue being sampled, which are labelled at the top of each column.

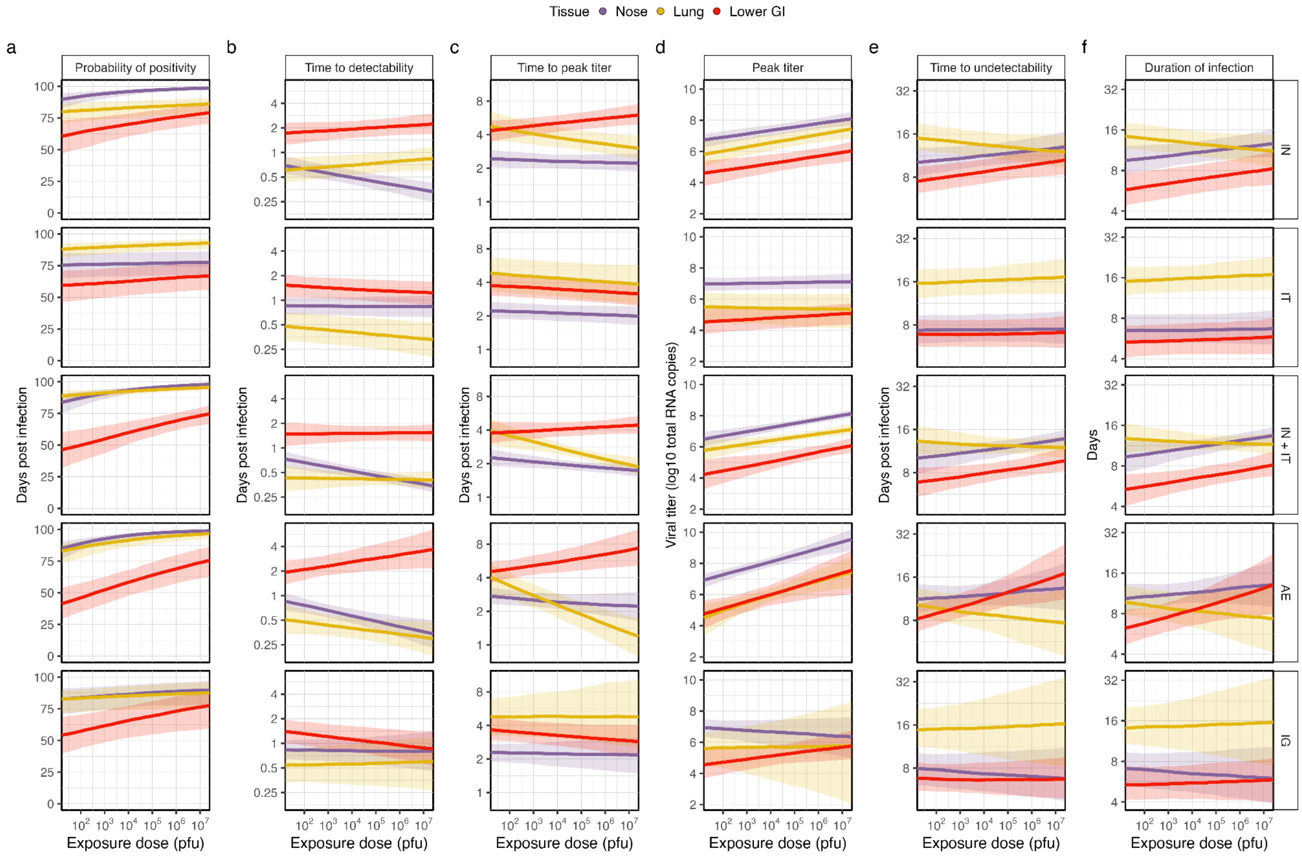


**Figure S29. Dose effects on total RNA metrics for all exposure routes.**

Visualization techniques are the same as for Extended Data Figure 4, except this figure includes predictions for all exposure routes when monitoring infection using total RNA PCR assays. This figure also includes the probability of positivity. Each row shows the results for one exposure route, as indicated with facet labels along the right-hand side of the whole figure. See the next two figures for the significance of these effects as well as dose-specific trajectories.

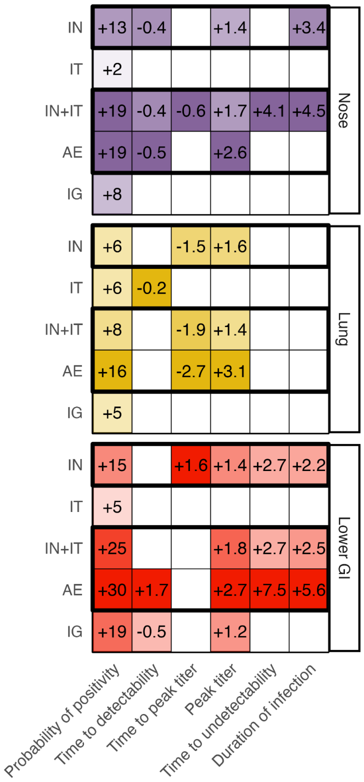


**Figure S30. Significance of the effect of dose on infection metrics when monitoring via total RNA.** The annotated text indicates the mean difference in the predictions between the maximum ($10^{7.4}$ pfu) and minimum doses in our database ($10^{1.2}$ pfu). A "+" indicates the metric significantly increases as dose increases. A "-" indicates the metric significantly decreases as dose increases. No label and a white cell indicates that the difference is not significant. Units depend on the metric (probability: percent; event time: days; peak titer: log10 total RNA copies). Intensities of the colors for significant effects are scaled against the relative effect standardized within each column and panel; more intense colors indicate a larger effect. Dark boxes enclose the routes that included upper respiratory exposure. Predictions integrate across all demographic factors.

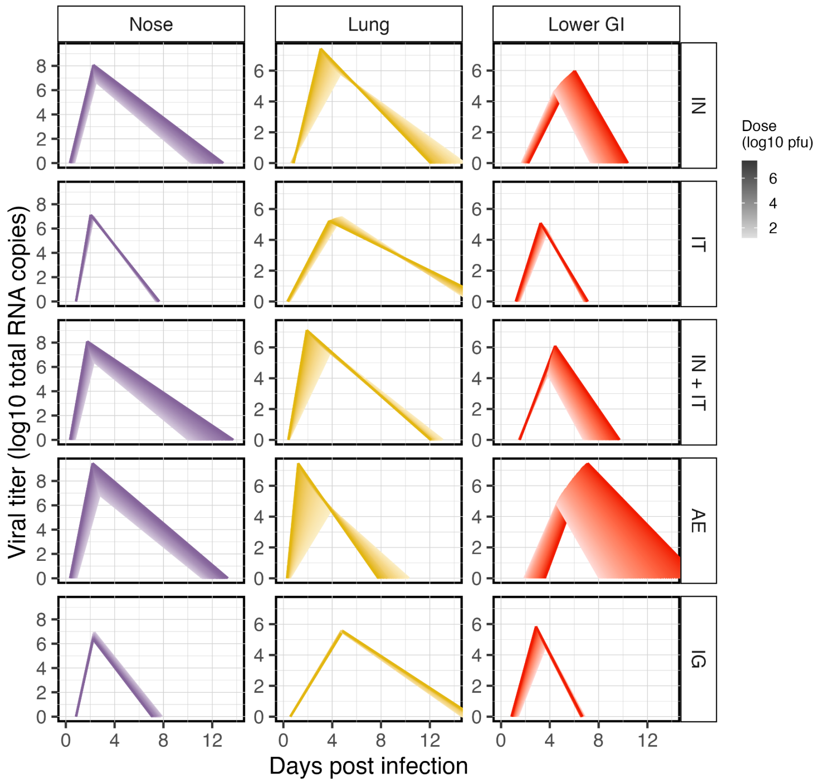


**Figure S31. Dose effects on total RNA trajectories for all exposure routes.**

Results are presented as in Figure 5d with lighter colors corresponding to smaller exposure doses. Different exposure routes are shown in distinct rows. Colors correspond with the tissue being sampled, which are labelled at the top of each column.

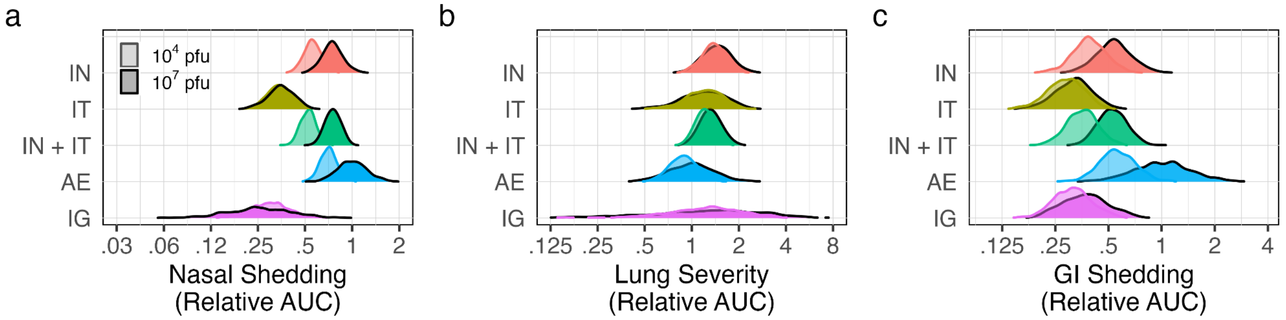


**Figure S32. Variability in clinical profiles across exposure routes and doses.**

Panels display the variability in the AUC values in the nose (a), lung (b), and GI (c) for each route and dose relative to the median AUC value for AE exposure with $10^7$ pfu. Colors correspond with the route, and intensity indicates the dose (lighter colors: $10^4$ pfu; darker colors: $10^7$ pfu).

### Supplementary Tables

**Table S1. Database summary.**

Each row is for a distinct exposure route, so articles with multiple routes have multiple rows. For articles where the first author has the same last name as another author, we include the first name of the first author in parentheses. Acronyms are: IN, intranasal; IT, intratracheal; IC, intracranial; OR, oral; OC: ocular; CJ: conjunctival; IV, intravenous; AE, aerosol; IB, intrabronchial; EB: endobronchial; U, unknown; J, juvenile; A, adult; G, geriatric; M, male; F, female; RM, rhesus macaque; CM, cynomolgus macaque; AGM, African green monkey; T, total RNA; G, genomic RNA; SG, subgenomic RNA; VeroE6-SS2, VeroE6-TMPRS22; I, invasive; NI, non-invasive.

*: dose reported as TCID50 was converted to pfu.

†: includes many doses in the indicated range.

§: more than this number of animals were included in the indicated study, but we were only able to extract individual-level data for some of them.

| Article | # indivs. | # obs. | Route | Dose (log10 pfu) | Age Class | Sex | Species | PCR type | PCR gene | Culture assay | Cell line | Sample Type | Strain |
|---|---|---|---|---|---|---|---|---|---|---|---|---|---|
| **An et al. 2021** | 3 | 42 | IT | 6.1* | J | U | RM | T | N | | | I | CHN/U (ID: 20SF107) |
| **Arunachalam et al. 2020** | 4 | 32 | IT, IN | 6.5 | J, A, U | M | RM | SG | N, E | | | NI | USA/WA1/2020 |
| **Baum et al. 2020** | 10 | 183 | IT, IN | 6, 5 | A, U | M, F | RM | T, SG | N, E | | | NI | USA/WA1/2020 |
| **Berry et al. 2022** | 6 | 262 | IN | 4.5* | U | F | CM | T, SG | E, ORF7 | U | VeroE6-SS2 | NI, I | AUS/VIC01/2020 |
| **Bewley et al. 2020** | 6 | 43 | IT, IN | 6.7 | J | U | RM | T | N | | | NI | AUS/VIC01/2020 |
| **Bixler et al. 2022** | 8 | 264 | AE | 4.8 | A | U | RM, CM | T, SG | U, E | PFU | Vero 76 | NI | USA/WA1/2020 |
| **Bixler et al. 2022** | 8 | 264 | IT, IN | 7.4 | A | U | RM, CM | T, SG | U, E | PFU | Vero 76 | NI | USA/WA1/2020 |
| **Blair et al. 2021** | 4 | 92 | AE | 4 | A, G | M, F | RM, AGM | T | N | | | NI | USA/WA1/2020 |
| **Blair et al. 2021** | 4 | 89 | IT, IN, OR, CJ | 6.6 | A, G | M, F | RM, AGM | T | N | | | NI | USA/WA1/2020 |
| **Böszörményi et al. 2020** | 8 | 1180 | IT, IN | 4.8* | J, A | M | RM, CM | G, SG | ORF1, E | | | NI, I | DEU/BavPat1/2020 |
| **Brouwer et al. 2021** | 4 | 124 | IT, IN | 6 | A | F | CM | G, SG | ORF1, E | | | NI | FRA/1DF0372/2020 |
| **Chandrashekar et al. 2020** | 13 | 341 | IT, IN | 6, 5, 4 | A | U | RM | T, SG | N, E | PFU | Vero E6 | NI, I | USA/WA1/2020 |
| **Chen et al. 2021** | 8§ | 34 | IN | 5.1* | J | U | RM | U | U | | | I | CHN/KMS1/2020 |
| **Corbett et al. 2020** | 15 | 63 | EB | 4, 5 | A | M, F | RM | U | U | | | NI | USA/WA1/2020 |
| **Corbett et al. 2020** | 23 | 167 | IT, IN | 5.9, 5, 4 | J, A | M, F | RM | T, SG, U | N, E, U | | | NI | USA/WA1/2020 |
| **Cross et al. 2020** | 6 | 403 | IN | 6.4 | A | F | AGM | T | N | PFU | Vero E6 | NI, I | ITA/INMI1/2020 |
| **Cross et al. 2021** | 2 | 85 | IT, IN | 5.7 | U | U | AGM | T | N | PFU | Vero E6 | NI, I | ITA/INMI1/2020 |
| **Dabisch et al. 2021** | 16 | 448 | AE | 1.2-3.5*† | J, A | M, F | CM | G | ORF1 | TCID50 | Vero | NI | USA/WA1/2020 |
| **Dagotto et al. 2021** | 4 | 16 | IT, IN | 4 | A | U | RM | T | N, E | | | NI | USA/WA1/2020 |
| **Deng et al. 2020A** | 7 | 207 | IT | 5.8* | J | U | RM | T | E | | | NI, I | CHN/WH-09/2020 |
| **Deng et al. 2020B** | 2 | 90 | CJ | 5.8* | J | M | RM | T | E | TCID50 | Vero E6 | NI, I | CHN/WH-09/2020 |
| **Deng et al. 2020B** | 2 | 83 | IG | 5.8* | J | M | RM | T | E | | | NI, I | CHN/WH-09/2020 |
| **Deng et al. 2020B** | 1 | 74 | IT | 5.8* | J | M | RM | T | E | TCID50 | Vero E6 | NI, I | CHN/WH-09/2020 |
| **Fears et al. 2022** | 8 | 544 | AE | 4* | A | M | RM, AGM | T, SG | N | TCID50 | Vero E6 | NI, I | USA/WA1/2020 |
| **Fears et al. 2022** | 8 | 543 | IT, IN | 6.1* | A | M | RM, AGM | T, SG | N | TCID50 | Vero E6 | NI, I | USA/WA1/2020 |
| **Feng et al. 2020** | 6 | 63 | IT | 4.2, 2.4* | A | M, F | RM | T | S | | | NI, I | CHN/WIV04/2019 |
| **Finch et al. 2020** | 3 | 117 | IB | 6.6 | J | M, F | CM | T | N | | | NI | USA/WA1/2020 |

| | | | | | | | | | | | | | |
|---|---|---|---|---|---|---|---|---|---|---|---|---|---|
| **Fischer et al. 2024** | 4 | 68 | AE | 2.8-3.3*† | J | M | RM | G | ORF1 | | | NI, I | USA/WA1/2020 |
| **Fischer et al. 2024** | 4 | 68 | IN | 5.7* | J | M | RM | G | ORF1 | | | NI, I | USA/WA1/2020 |
| **Francica et al. 2021** | 7 | 42 | IT, IN | 6.5 | U | U | RM | SG | E | | | NI | USA/WA1/2020 |
| **Furuyama et al. 2022** | 4 | 124 | IT, IN, OR, OC | 6.3* | U | F | RM | T, SG | E | | | NI, I | USA/WA1/2020 |
| **Gabitzsch et al. 2021** | 2 | 48 | IT, IN | 5.8* | J | M, F | RM | T, SG | N, E | | | NI | USA/WA1/2020 |
| **Gorman et al. 2021** | 4 | 64 | IT, IN | 6 | A | U | RM | T, SG | N, E | | | NI, I | USA/WA1/2020 |
| **Gu et al. 2020** | 1 | 1 | IT, IN | 6.6 | A | M | RM | T | E | | | I | KOR/U (ID: NCCP43326) |
| **Gu et al. 2021** | 1 | 7 | IT | 4.8* | A | U | RM | T | S | | | NI | CHN/WIV04/2019 |
| **Guebre-Xabier et al. 2020** | 4 | 16 | IT, IN | 4 | J, A | M, F | CM | SG | E | | | NI | USA/WA1/2020 |
| **Guo et al. 2021** | 3 | 45 | IT | 4.8* | A | U | RM | T | N | | | NI, I | CHN/WIV04/2019 |
| **Hassan et al. 2021** | 6 | 138 | IN, IB | 5.8* | A | U | RM | U, G | N, ORF1 | TCID50 | Vero E6 | NI, I | USA/WA1/2020 |
| **He et al. 2021** | 10 | 20 | IT, IN | 4 | A | U | RM | SG | E | | | NI | USA/WA1/2020 |
| **Hoang et al. 2021** | 4 | 164 | IT, IN | 6 | A | M, F | RM | T | N | | | NI, I | USA/WA1/2020 |
| **Huang et al. 2021** | 8 | 104 | AE | 4* | A | M | RM, AGM | T | N | | | NI | USA/WA1/2020 |
| **Ishigaki et al. 2021** | 3 | 393 | IT, IN, OR, CJ | 6.2* | A | M, F | CM | T | N | TCID50 | Vero E6 | NI, I | JPN/WK-521/2020 |
| **Ishii et al. 2022** | 4 | 48 | IN | 4.8* | J | F | CM | T, SG | U | TCID50 | Vero E6-SS2 | NI, I | JPN/WK-521/2020 |
| **Jiao et al. 2021A** | 5 | 335 | IG | 7 | U | M | RM | T | N | TCID50 | Vero E6 | NI, I | CHN/U |
| **Jiao et al. 2021A** | 5 | 326 | IN | 7 | U | M | RM | T | N | TCID50 | Vero E6 | NI, I | CHN/U |
| **Jiao et al. 2021B** | 2 | 100 | IC | 5, 6 | J | M | RM | T | N | | | NI, I | CHN/U |
| **Jiao et al. 2021B** | 5 | 147 | IN | 7 | J | M | RM | T | N | | | NI, I | CHN/U |
| **Johnston et al. 2020** | 11 | 473 | AE | 4.5, 4.6, 4.7 | J, A | M, F | RM, CM, AGM | T | N | PFU | Vero 76 | NI | USA/WA1/2020 |
| **Jones et al. 2021** | 4 | 88 | IT, IN | 5 | A | F | RM | T, SG | N, E | | | NI, I | USA/WA1/2020 |
| **Kim et al. 2021** | 3 | 90 | IT, IN, OR, OC | 6.3* | A | U | RM | T | S | TCID50 | Vero E6 | NI, I | KOR/U |
| **Kobiyama et al. 2021** | 2 | 76 | IT, IN, OR, CJ | 7.3 | A | F | CM | T | N | TCID50 | VeroE6-SS2 | NI, I | U |
| **Koo et al. 2020** | 16 | 567 | IT, IN, OR, CJ, IV | 7.3* | U | U | RM, CM | G | ORF1 | TCID50 | Vero | NI, I | KOR/U (ID: NCCP43326) |
| **Lakshmanappa et al. 2021** | 6 | 24 | IT, IN, OC | 6.4 | J | M, F | RM | T | N | | | NI | USA/CA-CZB-59X002/2020 |
| **Lambe et al. 2021** | 6 | 26 | IT, IN | 6.7 | J | U | RM | T | N | | | NI | AUS/VIC01/2020 |
| **Li (Dandan) et al. 2021** | 2 | 34 | IN | 4.8* | J | M | RM | T | N | | | NI, I | CHN/KMS1/2020 |
| **Li (Dapeng) et al. 2021** | 11 | 182 | IT, IN | 5 | J, A | M, F | CM | T, SG | E, N | | | NI | USA/WA1/2020 |
| **Li (Mingxi) et al. 2021** | 2 | 24 | IT | 6 | A | M | RM | U, SG | U, E | | | NI, I | CHN/Wuhan-Hu-1/2020 |
| **Li (Yuzhong) et al. 2021** | 5 | 15 | IT | 6.5* | J | U | RM | T | N | | | I | CHN/Kunming-BP16/2020 |
| **Liang et al. 2021** | 6 | 136 | IT, IN | 6.3* | U | M | RM | T | N | | | NI, I | CHN/U (ID: 107) |
| **Liu (Xiaolei) et al. 2022** | 4 | 72 | IN | 4.8* | J | M, F | RM | T | E | | | NI, I | CHN/KMS1/2020 |
| **Liu (Jiang-Feng) et al. 2022** | 5 | 10 | IN | 5.7 | A | U | RM | T | N | | | I | CHN/U (ID: GD108#) |
| **Liu (Zezhong) et al. 2022** | 3 | 48 | IT | 5.8* | J | U | RM | T | N | | | NI, I | CHN/WH-09/2020 |
| **Lu et al. 2020A** | 20 | 760 | IT, IN, CJ | 6.7, 6.4 | J, A | M, F, U | RM, CM | T | N | | | NI, I | CHN/Wuhan-Hu-1/2020 |
| **Lu et al. 2020B** | 3 | 27 | IT, IN, CJ | 6.7 | J, A | F | RM | T | N | | | NI | CHN/Wuhan-Hu-1/2020 |
| **Ma et al. 2022** | 4 | 92 | IT | 4.8 | J | M, F | RM | T | S | | | NI, I | CHN/WIV04/2019 |
| **Maisonnasse et al. 2020** | 8 | 214 | IT, IN | 6 | J | M, F | CM | G | ORF1 | | | NI | FRA/1DF0372/2020 |
| **Maisonnasse et al. 2021** | 5 | 100 | IT, IN | 6 | U | F | CM | G, SG | ORF1, E | | | NI | FRA/1DF0372/2020 |
| **McMahan et al. 2020** | 5 | 56 | IT, IN | 4 | J | M, F, U | RM | SG | E | | | NI | USA/WA1/2020 |
| **Mercado et al. 2020** | 20 | 204 | IT, IN | 4 | A | U | RM | SG | E | PFU | Vero E6 | NI | USA/WA1/2020 |
| **Munster et al. 2020** | 8 | 646 | IT, IN, OR, OC | 6.3* | J, A | M, F | RM | T, SG | E, ORF7 | TCID50 | Vero E6 | NI, I | USA/WA1/2020 |
| **Nagata et al. 2021** | 6 | 495 | IT, IN, CJ | 7.4* | A | F | CM | T, SG | N | TCID50 | VeroE6-SS2 | NI, I | JPN/WK-521/2020 |
| **Nawaz et al. 2020** | 4 | 59 | IN, OR | 6.1* | U | U | RM | U | U | | | NI, I | PAK/Lahore-IV/2020 |
| **Nomura et al. 2022** | 6 | 295 | IN | 5.7, 4.7, 3.7* | J, A | M, F | CM | T, SG | U | TCID50 | VeroE6-SS2 | NI | JPN/WK-521/2020 |
| **Pan et al. 2022** | 4 | 68 | IT | 4.8* | A | U | RM | T | S | | | NI, I | CHN/WIV04/2019 |
| **Patel et al. 2021** | 5 | 130 | IT, IN | 4 | J | M, F | RM | T, SG | N, E | | | NI | USA/WA1/2020 |
| **Philippens et al. 2021** | 8 | 224 | IT, IN | 4.8* | J, A | M | RM, CM | G, SG | ORF1, E | | | I | DEU/BavPat1/2020 |
| **Qin et al. 2020** | 2 | 38 | IN | 4.8* | J | F | RM | U | U | | | NI, I | CHN/KMS1/2020 |
| **Rauch et al. 2020** | 6 | 88 | IT, IN | 6.7 | U | M, F, U | RM | U, SG | N, E | | | NI, I | AUS/VIC01/2020 |
| **Rockx et al. 2020** | 8 | 371 | IT, IN | 6.8* | A | F | CM | U | U | TCID50 | Vero E6 | NI, I | DEU/BavPat1/2020 |

| | | | | | | | | | | | | | |
|---|---|---|---|---|---|---|---|---|---|---|---|---|---|
| **Roozendaal et al. 2021** | 7 | 84 | IT, IN | 4.8* | J | M, F | RM | SG | E | | | NI | USA/WA1/2020 |
| **Rosenke et al. 2020** | 10 | 482 | IT, IN, OR, OC | 6.3* | J | M | RM | U | E | TCID50 | Vero E6 | NI, I | USA/WA1/2020 |
| **Rosenke et al. 2021** | 5 | 575 | IN | 6 | A, U | M, F | AGM | SG, T | E, N | TCID50 | Vero E6 | NI, I | USA/RML-7/2020 |
| **Routhu et al. 2021** | 5 | 60 | IT | 4.7 | J | M | RM | SG | E | | | NI | USA/WA1/2020 |
| **Salguero et al. 2021** | 12 | 437 | IT, IN | 6.7 | J | M, F, U | RM, CM | T, SG | N, E | PFU | Vero E6 | NI, I | AUS/VIC01/2020 |
| **Sanchez-Felipe et al. 2021** | 6 | 24 | IT, IN | 4* | J, A | M | CM | G | ORF1 | | | NI | BEL/GHB-03021/2020 |
| **Saunders et al. 2021** | 5 | 46 | IT, IN | 5 | J, A | M, F | CM | SG | E, N | PFU | Vero E6 | NI | USA/WA1/2020 |
| **Seo et al. 2021** | 3 | 54 | IT, IN, OR, CJ, IV | 7.3* | U | U | CM | G | ORF1 | TCID50 | Vero | NI | KOR/U (ID: NCCP43326) |
| **Shan et al. 2020** | 8 | 330 | IT | 6.7, 5.8* | A | M, F | RM | T | S | TCID50 | Vero E6 | NI, I | CHN/WIV04/2019 |
| **Shi et al. 2020** | 1§ | 1 | IT | 4.8* | A | U | RM | T | S | | | NI | CHN/IVDC-HB-envF13/2020 |
| **Singh et al. 2020** | 16 | 592 | IT, IN, OC | 6 | J, G | M, F | RM | T, SG | N, E | PFU | Vero E6 | NI, I | USA/WA1/2020 |
| **Sokol et al. 2021** | 4 | 83 | IT, IN | 7.3 | J, A | F | RM, CM | G | ORF1 | | | NI | FRA/1DF0372/2020 |
| **Song et al. 2020** | 8 | 280 | IT | 6.8* | A | M | RM | T | N | | | NI, I | CHN/U (ID: 107) |
| **Song et al. 2021** | 3 | 48 | IT | 6.8* | A | M | RM | U | U | | | NI, I | CHN/U (ID: 107) |
| **Song et al. 2021** | 3 | 48 | IT, IN | 6.8* | A | M | RM | U | U | | | NI, I | CHN/U (ID: 107) |
| **Speranza et al. 2020** | 8 | 557 | IT, IN, OR, OC | 6.3* | A | M, F | AGM | T, SG | E | TCID50 | Vero E6 | NI, I | USA/WA1/2020 |
| **Sun (Shiyu) et al. 2021** | 6 | 180 | IN | 6.8* | J | M, F | RM | T | N | | | NI, I | CHN/U (ID: 107) |
| **Sun (Shihui) et al. 2021** | 4 | 24 | IT, IN, OC | 6.3* | J | M, F | CM | T | S | | | I | CHN/IME-BJ01/2020 |
| **Sun et al. 2023** | 4 | 83 | IN | 5.7 | A | U | RM | T | N | | | NI, I | CHN/U (ID: GD108#) |
| **Tan (Shudan) et al. 2023** | 4 | 148 | IT | 4.8* | A | M, F | RM | T | S | | | NI, I | CHN/WIV04/2019 |
| **Tan (Janessa) et al. 2023** | 4 | 144 | IT | 6.3* | U | U | CM | T | N | | | NI | SG/WX-56/2020 |
| **van Doremalen et al. 2020** | 6 | 456 | IT, IN, OR, OC | 6.3* | J | U | RM | G, SG | ORF1, E | TCID50 | Vero E6 | NI, I | USA/WA1/2020 |
| **Vogel et al. 2021** | 9 | 130 | IT, IN | 6 | J | M | RM | T | N | | | NI | USA/WA1/2020 |
| **Wang (Gan) et al. 2020** | 3 | 90 | IT, IN | 6.8* | A | M | RM | T | N | | | NI, I | CHN/U (ID: GD108#) |
| **Wang (Shuang) et al. 2020** | 3 | 84 | IT | 4.8* | A | U | RM | T | S | | | NI, I | CHN/WIV04/2019 |
| **Wang (Hui) et al. 2020** | 2 | 26 | IT | 5.8* | J | U | RM | T | N | | | NI, I | CHN/WH-09/2020 |
| **Wang et al. 2022** | 6 | 180 | IT | 4.8* | A | M, F | RM | SG | S | | | NI, I | CHN/WIV04/2019 |
| **Williamson et al. 2020** | 6 | 441 | IT, IN, OR, OC | 6.3* | J | M, F | RM | T | E | TCID50 | Vero E6 | NI, I | USA/WA1/2020 |
| **Woolsey et al. 2020** | 6 | 470 | IT, IN | 5.7 | A | M, F, U | AGM | T | N | PFU | Vero E6 | NI, I | ITA/INMI1/2020 |
| **Yadav et al. 2021A** | 5 | 313 | IT, IN | 6.5* | A | U | RM | T, SG | E | | | NI, I | IND/UN-770/2020 |
| **Yadav et al. 2021B** | 4 | 262 | IT, IN | 6.5* | A | U | RM | T, SG | E | U | Vero CCL-81 | NI, I | U |
| **Yang et al. 2020** | 5 | 166 | IN | 5.7 | A | U | RM | T, SG | N, E | | | NI, I | U |
| **Yao et al. 2021** | 3 | 30 | IT | 5.8* | A | U | RM | T | S | | | NI, I | CHN/WIV04/2019 |
| **Yu (Jingyou) et al. 2020** | 10 | 209 | IT, IN | 4 | A | U | RM | T, SG | N, E | PFU | Vero E6 | NI | U |
| **Yu (Pin) et al. 2020** | 5 | 21 | IT | 5.8* | J, A | U | RM | T | E | | | NI, I | CHN/IVDC-HB-01/2020 |
| **Yu et al. 2022** | 9 | 252 | IG | 5.8* | A | U | RM | T, SG | N, E | | | NI | USA/WA1/2020 |
| **Yu et al. 2022** | 12 | 240 | IT, IN | 4.8* | A | U | RM | SG | E | | | NI | USA/WA1/2020 |
| **Zheng et al. 2020** | 14 | 670 | IN | 5.1* | J | M | RM | G | ORF1 | TCID50 | Vero | NI, I | CHN/KMS1/2020 |
| **Zost et al. 2020** | 4 | 20 | IT, IN | 4 | A | U | RM | SG | E | | | NI | USA/WA1/2020 |

**Table S2. Categorization of reported inoculation routes into exposure categories for model fitting.** Acronyms are as follows: IN: intranasal; OC: ocular; IT: intratracheal; IB: intrabronchial; EB: endobronchial; OR: oral; IV: intravenous; AE: aerosol; IG: intragastric. Underlines indicate the route used when generating predictions for each category.

| Exposure category | URT | LRT | URT + LRT (Liquid) | URT + LRT (Aerosol) | GI |
|---|---|---|---|---|---|
| **Reported Inoculation Routes** | IN<br>OC | IT<br>IB<br>EB | IT, IN<br>IB, IN<br>IT, IN, OC<br>IT, IN, OR, OC<br>IT, IN, OR, OC, IV | AE | IG |

**Table S3. Reported sample types grouped into tissue categories.**

| Tissue Group | Non-invasive sample types | Invasive sample types |
|---|---|---|
| **Nose** | Nasal swab / wash<br>Nasopharyngeal swab / fluid | Nasal mucosa / tissue |
| **Throat** | Throat swab<br>Pharyngeal swab<br>Oropharyngeal swab | Oropharynx<br>Laryngeal mucosa |
| **Trachea** | Tracheal swab / brush / fluid | Trachea<br>Carina |
| **Lung** | Bronchoalveolar lavage (BAL)<br>Bronchial brush / swab | Bronchus<br>Lung (any lobe) |
| **Upper GI** | | Esophagus<br>Stomach<br>Liver<br>Small intestine<br>Duodenum<br>Ileum<br>Jejunum |
| **Lower GI** | Rectal swab / fluid<br>Anal swab<br>Feces / Fecal swab | Colon<br>Cecum |

**Table S4. Parameter estimates and their credible intervals for the probability of positivity.**

All entries display the median parameter estimate followed in brackets by the 90% credible interval. Parameters are stratified into columns by tissue groups (upper respiratory tract, URT; lower respiratory tract, LRT; GI, gastrointestinal tract), with the cofactor group indicated by the lefthand label (e.g., age) and the particular cofactor indicated by the righthand label (e.g., juvenile vs. adult vs. geriatric). Cells with two parameter estimates separated by a "|" are tissue-specific, as follows: for URT, nose | throat; for LRT, trachea | lung; for GI, upper GI | lower GI. All parameters are explained in the methods. For all lab effects, see Figure S11.

| Cofactor | URT | LRT | GI | |
|---|---|---|---|---|
| | 0.54 [-0.09, 1.15] | 0.55 [-0.05, 1.14] | -0.82 [-1.55, -0.14] | **Intercept** |
| | 0.13 [-0.23, 0.49] | 0.2 [-0.15, 0.56] | 0.15 [-0.22, 0.53] | **Location** |
| IN | 0.38 [0.03, 0.74] ǀ 0.34 [-0.04, 0.71] | 0.07 [-0.29, 0.44] ǀ -0.22 [-0.57, 0.13] | 0.17 [-0.18, 0.52] ǀ 0.39 [0.03, 0.73] | **Route** |
| IT | -0.36 [-0.7, 0] ǀ 0.41 [0.04, 0.8] | 0.28 [-0.08, 0.65] ǀ 0.33 [-0.03, 0.67] | -0.01 [-0.41, 0.39] ǀ 0.41 [0.06, 0.78] | **Route** |
| IN+IT | -0.07 [-0.4, 0.27] ǀ -0.38 [-0.75, 0] | -0.31 [-0.67, 0.02] ǀ 0.39 [0.05, 0.72] | -0.35 [-0.71, 0] ǀ -0.23 [-0.58, 0.1] | **Route** |
| AE | -0.03 [-0.38, 0.32] ǀ -0.2 [-0.58, 0.18] | -0.16 [-0.56, 0.24] ǀ -0.2 [-0.58, 0.17] | 0 [-0.4, 0.39] ǀ -0.42 [-0.77, -0.07] | **Route** |
| IG | -0.04 [-0.45, 0.37] ǀ -0.05 [-0.44, 0.36] | -0.09 [-0.49, 0.31] ǀ -0.09 [-0.5, 0.32] | 0.04 [-0.37, 0.44] ǀ 0.02 [-0.36, 0.42] | **Route** |
| Nose | 1.07 [0.14, 2.26] ǀ 0.86 [0.11, 1.88] | 0.15 [0.01, 0.54] ǀ 0.19 [0.02, 0.63] | 0.46 [0.04, 1.29] ǀ 0.48 [0.05, 1.22] | **Dose** |
| Throat | 1.19 [0.16, 2.36] ǀ 0.71 [0.07, 1.72] | 0.16 [0.01, 0.6] ǀ 0.18 [0.01, 0.62] | 0.43 [0.04, 1.21] ǀ 0.42 [0.04, 1.17] | **Dose** |
| Trachea | 0.11 [0.01, 0.4] ǀ 1.1 [0.37, 1.85] | 0.31 [0.03, 0.87] ǀ 0.69 [0.15, 1.32] | 0.13 [0.01, 0.5] ǀ 0.34 [0.03, 0.92] | **Dose** |
| Lung | 0.58 [0.07, 1.42] ǀ 1.85 [0.58, 3.29] | 0.55 [0.04, 1.96] ǀ 0.86 [0.13, 1.86] | 1.2 [0.11, 3.37] ǀ 0.2 [0.02, 0.71] | **Dose** |
| Upper GI | 0.67 [0.07, 2.04] ǀ 0.65 [0.06, 1.97] | 0.43 [0.04, 1.61] ǀ 0.42 [0.04, 1.52] | 2.28 [0.83, 4.04] ǀ 1.2 [0.29, 2.25] | **Dose** |
| Juvenile | -0.07 [-0.4, 0.24] | -0.04 [-0.35, 0.27] | 0.1 [-0.23, 0.42] | **Age** |
| Adult | 0.04 [-0.28, 0.35] | 0.26 [-0.05, 0.57] | -0.12 [-0.44, 0.2] | **Age** |
| Geriatric | 0.04 [-0.36, 0.44] | -0.22 [-0.61, 0.17] | 0.01 [-0.37, 0.4] | **Age** |
| | -0.37 [-0.61, -0.14] | -0.03 [-0.27, 0.21] | 0.23 [-0.02, 0.49] | **Sex** |
| RM | 0.16 [-0.13, 0.46] | -0.14 [-0.44, 0.16] | -0.12 [-0.43, 0.18] | **Species** |
| CM | -0.25 [-0.56, 0.04] | -0.19 [-0.49, 0.12] | -0.38 [-0.69, -0.06] | **Species** |
| AGM | 0.1 [-0.24, 0.42] | 0.34 [0, 0.67] | 0.51 [0.17, 0.85] | **Species** |
| Total RNA | 0.73 [0.42, 1.04] | 0.66 [0.36, 0.94] | 0.86 [0.57, 1.15] | **Assay** |
| gRNA | 0.04 [-0.26, 0.34] | 0.27 [-0.02, 0.56] | 0.25 [-0.06, 0.55] | **Assay** |
| sgRNA | 0.2 [-0.1, 0.5] | -0.14 [-0.43, 0.15] | -0.42 [-0.73, -0.09] | **Assay** |
| Culture | -0.98 [-1.26, -0.68] | -0.77 [-1.08, -0.47] | -0.7 [-1.03, -0.38] | **Assay** |

**Table S5. Parameter estimates and their credible intervals for the time to detectability.**

All entries display the median parameter estimate followed in brackets by the 90% credible interval. Parameters are stratified into columns by tissue groups (upper respiratory tract, URT; lower respiratory tract, LRT; GI, gastrointestinal tract), with the cofactor group indicated by the lefthand label (e.g., age) and the particular cofactor indicated by the righthand label (e.g., juvenile vs. adult vs. geriatric). Cells with two parameter estimates separated by a "|" are tissue-specific, as follows: for URT, nose | throat; for LRT, trachea | lung; for GI, upper GI | lower GI. All parameters are explained in the methods. For all lab effects, see Figure S11.

<table>
<tr><th>Cofactor</th><th>URT</th><th>LRT</th><th>GI</th><th></th></tr>
<tr><td></td><td>1 [1, 1.02]</td><td>1.02 [1, 1.07]</td><td>1.02 [1, 1.06]</td><td>Shape</td></tr>
<tr><td></td><td>0.28 [-2.43, 2.95]</td><td>4.24 [0.19, 8.43]</td><td>-6.59 [-10.19, -2.76]</td><td>Intercept</td></tr>
<tr><td></td><td>0.5 [-0.99, 2.02]</td><td>0.58 [-1.03, 2.19]</td><td>-0.42 [-1.99, 1.16]</td><td>Location</td></tr>
<tr><td>IN</td><td>0.77 [-0.75, 2.32] | 0.48 [-1.08, 2.03]</td><td>-0.22 [-1.78, 1.41] | -1.03 [-2.59, 0.53]</td><td>0.2 [-1.44, 1.91] | -0.44 [-2.02, 1.13]</td><td>Route</td></tr>
<tr><td>IT</td><td>-0.31 [-1.89, 1.22] | 0.45 [-1.01, 1.93]</td><td>-0.51 [-2.13, 1.05] | 0.08 [-1.5, 1.69]</td><td>-0.04 [-1.71, 1.63] | 0.08 [-1.48, 1.65]</td><td>Route</td></tr>
<tr><td>IN+IT</td><td>0.57 [-0.85, 2.02] | -0.02 [-1.56, 1.43]</td><td>0.64 [-0.93, 2.26] | 1.78 [0.32, 3.3]</td><td>0.23 [-1.37, 1.86] | 0.77 [-0.77, 2.29]</td><td>Route</td></tr>
<tr><td>AE</td><td>-1.22 [-2.67, 0.23] | -0.38 [-1.89, 1.08]</td><td>0.01 [-1.63, 1.63] | -0.21 [-1.75, 1.35]</td><td>0.01 [-1.65, 1.65] | -1.31 [-2.92, 0.23]</td><td>Route</td></tr>
<tr><td>IG</td><td>0.03 [-1.62, 1.63] | 0 [-1.57, 1.59]</td><td>-0.08 [-1.73, 1.52] | -0.06 [-1.71, 1.57]</td><td>0.17 [-1.46, 1.76] | 0.48 [-1.11, 2.08]</td><td>Route</td></tr>
<tr><td>Nose</td><td>3.01 [-1.18, 7.26] | 1.17 [-3.06, 5.44]</td><td>3.9 [-0.97, 8.77] | -2.09 [-6.87, 2.52]</td><td>4.55 [-0.56, 9.55] | -2.36 [-6.86, 1.87]</td><td>Dose</td></tr>
<tr><td>Throat</td><td>5.29 [1.07, 9.43] | 2.94 [-1.33, 7.2]</td><td>3.95 [-0.94, 8.79] | -1.41 [-6.06, 3.39]</td><td>4.51 [-0.42, 9.42] | -0.41 [-4.69, 3.96]</td><td>Dose</td></tr>
<tr><td>Trachea</td><td>0.19 [-2.57, 2.93] | 4.11 [1.33, 6.92]</td><td>1.29 [-3.12, 5.71] | 4.61 [1.07, 8.17]</td><td>1.59 [-3.55, 6.98] | 2.44 [-0.89, 5.67]</td><td>Dose</td></tr>
<tr><td>Lung</td><td>1.85 [-1.68, 5.46] | -0.2 [-4.36, 4.01]</td><td>-0.02 [-5.71, 5.66] | 5.01 [0.98, 9.09]</td><td>0.03 [-5.76, 5.75] | -6.28 [-10.95, -1.62]</td><td>Dose</td></tr>
<tr><td>Upper GI</td><td>0.13 [-4.8, 5] | -0.03 [-5.06, 4.64]</td><td>-1.08 [-7.09, 4.82] | -0.86 [-6.38, 4.9]</td><td>2.22 [-3.18, 7.78] | 5.52 [0.83, 10.18]</td><td>Dose</td></tr>
<tr><td>Juvenile</td><td>1.88 [0.67, 3.11]</td><td>0.6 [-0.81, 2.04]</td><td>1.27 [-0.19, 2.75]</td><td>Age</td></tr>
<tr><td>Adult</td><td>0.99 [-0.23, 2.23]</td><td>1.5 [0.09, 2.92]</td><td>-0.74 [-2.19, 0.71]</td><td>Age</td></tr>
<tr><td>Geriatric</td><td>-2.56 [-4.01, -1.06]</td><td>-1.65 [-3.2, -0.13]</td><td>-0.4 [-2, 1.15]</td><td>Age</td></tr>
<tr><td></td><td>1.32 [0.32, 2.33]</td><td>1.07 [-0.3, 2.49]</td><td>-0.37 [-1.72, 1.01]</td><td>Sex</td></tr>
<tr><td>RM</td><td>0.71 [-0.5, 1.88]</td><td>0.48 [-0.92, 1.89]</td><td>1.03 [-0.4, 2.45]</td><td>Species</td></tr>
<tr><td>CM</td><td>-0.85 [-2.05, 0.4]</td><td>1.21 [-0.29, 2.73]</td><td>0.93 [-0.59, 2.43]</td><td>Species</td></tr>
<tr><td>AGM</td><td>0.52 [-0.82, 1.86]</td><td>-1.19 [-2.7, 0.28]</td><td>-1.82 [-3.3, -0.36]</td><td>Species</td></tr>
<tr><td>Total RNA</td><td>-0.07 [-1.21, 1.08]</td><td>-0.47 [-1.8, 0.95]</td><td>2.07 [0.66, 3.48]</td><td>Assay</td></tr>
<tr><td>gRNA</td><td>1.71 [0.47, 2.91]</td><td>1.19 [-0.29, 2.68]</td><td>-0.45 [-1.88, 1.01]</td><td>Assay</td></tr>
<tr><td>sgRNA</td><td>-0.51 [-1.7, 0.66]</td><td>0.35 [-1.06, 1.72]</td><td>-0.8 [-2.33, 0.72]</td><td>Assay</td></tr>
<tr><td>Culture</td><td>-0.78 [-2.02, 0.45]</td><td>-0.61 [-2.11, 0.85]</td><td>-0.65 [-2.24, 0.97]</td><td>Assay</td></tr>
</table>

**Table S6. Parameter estimates and their credible intervals for the time to peak titer.**

All entries display the median parameter estimate followed in brackets by the 90% credible interval. Parameters are stratified into columns by tissue groups (upper respiratory tract, URT; lower respiratory tract, LRT; GI, gastrointestinal tract), with the cofactor group indicated by the righthand row label (e.g., age) and the particular cofactor indicated by the lefthand row label (e.g., juvenile vs. adult vs. geriatric). Cells with two parameter estimates separated by a "|" are tissue-specific, as follows: for URT, nose | throat; for LRT, trachea | lung; for GI, upper GI | lower GI. All parameters are explained in the methods. The Time cofactor group accounts for the dependence of the time to peak titer on the time to detectability. For all lab effects, see Figure S11.

| Cofactor | URT | LRT | GI | |
|---|---|---|---|---|
| | 1.35 [1.24, 1.46] | 3.18 [2.62, 3.85] | 1.04 [1, 1.12] | Shape |
| | -3.74 [-6.45, -1.04] | -11.41 [-15.01, -7.56] | -8.71 [-12.81, -4.38] | Intercept |
| | -0.03 [-1.54, 1.47] | 0.19 [-1.42, 1.76] | 0.07 [-1.52, 1.7] | Location |
| IN | -0.23 [-1.69, 1.27] \| 0.05 [-1.43, 1.56] | -0.11 [-1.74, 1.47] \| -0.36 [-1.93, 1.18] | 0 [-1.66, 1.66] \| -0.39 [-1.99, 1.15] | Route |
| IT | 0.61 [-0.88, 2.14] \| -0.21 [-1.68, 1.26] | -0.66 [-2.25, 0.97] \| -0.45 [-2.05, 1.18] | 0 [-1.62, 1.64] \| 0.29 [-1.3, 1.86] | Route |
| IN+IT | 0.81 [-0.57, 2.21] \| 0.11 [-1.35, 1.62] | 0.61 [-0.98, 2.18] \| 1.01 [-0.48, 2.46] | 0.03 [-1.59, 1.65] \| 0.54 [-1.01, 2.12] | Route |
| AE | -1 [-2.46, 0.47] \| 0.3 [-1.27, 1.85] | -0.01 [-1.63, 1.65] \| -0.03 [-1.63, 1.52] | -0.01 [-1.67, 1.65] \| -0.55 [-2.13, 1.03] | Route |
| IG | 0.12 [-1.48, 1.72] \| -0.29 [-1.94, 1.33] | -0.01 [-1.63, 1.66] \| 0 [-1.62, 1.68] | -0.02 [-1.66, 1.67] \| 0.19 [-1.44, 1.77] | Route |
| Nose | -2.1 [-6.17, 2.02] \| -0.82 [-5.01, 3.38] | 4.03 [-0.62, 8.59] \| 3.45 [-1.27, 8.15] | 0.18 [-5.56, 6.01] \| -2.06 [-6.57, 2.42] | Dose |
| Throat | 1.05 [-3.11, 5.12] \| 0.24 [-3.88, 4.36] | 4.39 [-0.33, 8.88] \| 3.94 [-0.8, 8.63] | 0.09 [-5.5, 5.75] \| -2.11 [-6.7, 2.56] | Dose |
| Trachea | 2.09 [-0.51, 4.69] \| 1.57 [-1.08, 4.26] | 0.02 [-3.68, 3.64] \| 1.93 [-1.46, 5.39] | 0.13 [-5.58, 5.9] \| 1.76 [-1.51, 5] | Dose |
| Lung | -0.9 [-4.18, 2.4] \| 1.15 [-3.06, 5.34] | -0.02 [-5.78, 5.55] \| 5.2 [1.2, 9.7] | -0.03 [-5.85, 5.92] \| -0.85 [-5.54, 3.82] | Dose |
| Upper GI | 1.32 [-3.63, 5.91] \| -3.73 [-8.08, 0.4] | -0.01 [-5.75, 5.8] \| 0.07 [-5.37, 5.97] | 0 [-5.88, 5.69] \| 1.57 [-3.06, 6] | Dose |
| Juvenile | 0.42 [-0.83, 1.64] | 0.71 [-0.63, 2.01] | 0.95 [-0.48, 2.35] | Age |
| Adult | 1.04 [-0.21, 2.24] | -0.24 [-1.56, 1.04] | -0.6 [-2.04, 0.8] | Age |
| Geriatric | -1.21 [-2.73, 0.32] | -0.4 [-2, 1.11] | -0.26 [-1.87, 1.28] | Age |
| | -1.56 [-2.46, -0.67] | -0.79 [-1.93, 0.4] | 0.18 [-1.21, 1.53] | Sex |
| RM | -0.4 [-1.53, 0.75] | -0.36 [-1.63, 0.94] | 0.55 [-0.84, 1.95] | Species |
| CM | 0.22 [-0.97, 1.4] | 1.94 [0.53, 3.36] | 0.03 [-1.43, 1.5] | Species |
| AGM | 0.4 [-0.9, 1.69] | -1.59 [-3.04, -0.09] | -0.48 [-1.98, 1.02] | Species |
| Total RNA | -1.76 [-2.89, -0.63] | -1.43 [-2.73, -0.1] | -0.08 [-1.51, 1.37] | Assay |
| gRNA | -1.28 [-2.46, -0.12] | 0.17 [-1.18, 1.49] | -0.47 [-1.91, 1] | Assay |
| sgRNA | 0.69 [-0.48, 1.83] | 0.93 [-0.32, 2.2] | 0.27 [-1.3, 1.82] | Assay |
| Culture | 2.58 [1.29, 3.89] | 0.38 [-1.19, 1.99] | 0.33 [-1.27, 1.93] | Assay |
| Inoculated | -0.63 [-1.29, 0.02] | -0.75 [-1.89, 0.29] | -0.02 [-1.69, 1.67] | Time |
| Not Inoculated | 0.82 [-0.52, 2.14] | -2.13 [-2.98, -1.31] | 0.08 [-0.37, 0.54] | Time |

**Table S7. Parameter estimates and their credible intervals for the peak titer.**

All entries display the median parameter estimate followed in brackets by the 90% credible interval. Parameters are stratified into columns by tissue groups (upper respiratory tract, URT; lower respiratory tract, LRT; GI, gastrointestinal tract), with the cofactor group indicated by the righthand row label (e.g., age) and the particular cofactor indicated by the lefthand row label (e.g., juvenile vs. adult vs. geriatric). Cells with two parameter estimates separated by a "|" are tissue-specific, as follows: for URT, nose | throat; for LRT, trachea | lung; for GI, upper GI | lower GI. "SD" refers to the standard deviation. All parameters are explained in the methods. The Time cofactor group accounts for the dependence of the peak titer on the time to the peak titer. For all lab effects, see Figure S12.

<table>
<tr><th>Cofactor</th><th>URT</th><th>LRT</th><th>GI</th><th></th></tr>
<tr><td></td><td>5.33 [4.72, 5.94]</td><td>4.86 [3.81, 5.9]</td><td>3.12 [2.11, 4.1]</td><td>Intercept</td></tr>
<tr><td></td><td>-0.05 [-0.4, 0.31]</td><td>0.04 [-0.36, 0.43]</td><td>0 [-0.4, 0.4]</td><td>Location</td></tr>
<tr><td>IN</td><td>0.1 [-0.27, 0.47] | 0.05 [-0.31, 0.41]</td><td>-0.12 [-0.52, 0.29] | 0.05 [-0.34, 0.43]</td><td>0 [-0.42, 0.42] | 0.04 [-0.37, 0.45]</td><td>Route</td></tr>
<tr><td>IT</td><td>-0.12 [-0.48, 0.24] | 0.18 [-0.18, 0.54]</td><td>-0.17 [-0.57, 0.24] | 0 [-0.39, 0.39]</td><td>0 [-0.41, 0.42] | 0.12 [-0.27, 0.51]</td><td>Route</td></tr>
<tr><td>IN+IT</td><td>-0.16 [-0.45, 0.16] | -0.12 [-0.46, 0.24]</td><td>0.26 [-0.13, 0.65] | 0.07 [-0.26, 0.4]</td><td>0 [-0.41, 0.41] | -0.3 [-0.68, 0.07]</td><td>Route</td></tr>
<tr><td>AE</td><td>0.27 [-0.08, 0.61] | -0.13 [-0.53, 0.25]</td><td>0 [-0.42, 0.41] | -0.07 [-0.45, 0.3]</td><td>0 [-0.42, 0.42] | 0.13 [-0.26, 0.53]</td><td>Route</td></tr>
<tr><td>IG</td><td>-0.01 [-0.42, 0.39] | -0.02 [-0.43, 0.38]</td><td>0 [-0.4, 0.4] | 0 [-0.41, 0.42]</td><td>-0.01 [-0.42, 0.41] | 0.01 [-0.4, 0.43]</td><td>Route</td></tr>
<tr><td>Nose</td><td>0.65 [-0.82, 2.18] | 0.47 [-1, 1.96]</td><td>-0.29 [-2.41, 1.78] | 0.32 [-1.66, 2.27]</td><td>0 [-2.94, 2.89] | 0.24 [-1.74, 2.26]</td><td>Dose</td></tr>
<tr><td>Throat</td><td>0.91 [-0.58, 2.33] | 0.3 [-1.2, 1.72]</td><td>0.06 [-2.05, 2.19] | 1.46 [-0.48, 3.44]</td><td>-0.02 [-2.91, 2.81] | 1.03 [-1.03, 3.02]</td><td>Dose</td></tr>
<tr><td>Trachea</td><td>0.29 [-0.27, 0.84] | 0.09 [-0.52, 0.67]</td><td>1.71 [0.5, 2.99] | -0.31 [-1.09, 0.48]</td><td>-0.02 [-2.81, 2.84] | 0.68 [-0.13, 1.48]</td><td>Dose</td></tr>
<tr><td>Lung</td><td>1 [0.29, 1.71] | 2.59 [1.63, 3.53]</td><td>0.02 [-2.87, 2.92] | 0.68 [-0.41, 1.84]</td><td>0.02 [-2.79, 2.78] | 0.82 [-0.4, 2.05]</td><td>Dose</td></tr>
<tr><td>Upper GI</td><td>-0.6 [-1.9, 0.7] | -1.27 [-2.67, 0.15]</td><td>-0.05 [-2.95, 2.93] | 0 [-2.83, 2.86]</td><td>0.01 [-2.9, 2.88] | 1.45 [0.16, 2.7]</td><td>Dose</td></tr>
<tr><td>Juvenile</td><td>-0.02 [-0.31, 0.27]</td><td>0.02 [-0.28, 0.32]</td><td>-0.24 [-0.58, 0.1]</td><td>Age</td></tr>
<tr><td>Adult</td><td>-0.02 [-0.3, 0.27]</td><td>0.09 [-0.21, 0.39]</td><td>0.25 [-0.07, 0.57]</td><td>Age</td></tr>
<tr><td>Geriatric</td><td>0.09 [-0.28, 0.45]</td><td>-0.08 [-0.45, 0.3]</td><td>-0.01 [-0.4, 0.39]</td><td>Age</td></tr>
<tr><td></td><td>-0.67 [-0.82, -0.51]</td><td>0.03 [-0.19, 0.24]</td><td>-0.48 [-0.77, -0.19]</td><td>Sex</td></tr>
<tr><td>RM</td><td>-0.09 [-0.34, 0.17]</td><td>-0.32 [-0.6, -0.03]</td><td>0.04 [-0.29, 0.37]</td><td>Species</td></tr>
<tr><td>CM</td><td>0.28 [0.01, 0.54]</td><td>-0.16 [-0.48, 0.15]</td><td>-0.11 [-0.47, 0.23]</td><td>Species</td></tr>
<tr><td>AGM</td><td>-0.16 [-0.44, 0.12]</td><td>0.51 [0.19, 0.83]</td><td>0.07 [-0.28, 0.43]</td><td>Species</td></tr>
<tr><td>Total RNA</td><td>1.43 [1.19, 1.66]</td><td>0.99 [0.71, 1.27]</td><td>0.42 [0.1, 0.74]</td><td>Assay</td></tr>
<tr><td>gRNA</td><td>0.77 [0.53, 1.02]</td><td>0.85 [0.56, 1.14]</td><td>-0.13 [-0.46, 0.21]</td><td>Assay</td></tr>
<tr><td>sgRNA</td><td>0.07 [-0.18, 0.32]</td><td>-0.38 [-0.65, -0.11]</td><td>-0.05 [-0.42, 0.33]</td><td>Assay</td></tr>
<tr><td>Culture</td><td>-2.23 [-2.49, -1.96]</td><td>-1.44 [-1.76, -1.11]</td><td>-0.24 [-0.64, 0.14]</td><td>Assay</td></tr>
<tr><td>Inoculated</td><td>-0.14 [-0.21, -0.08]</td><td>-0.38 [-0.67, -0.09]</td><td>0 [-0.4, 0.41]</td><td>Time</td></tr>
<tr><td>Not Inoculated</td><td>0.27 [0.1, 0.45]</td><td>-0.04 [-0.24, 0.22]</td><td>0.22 [0.16, 0.29]</td><td>Time</td></tr>
<tr><td>True Titer</td><td>1.35 [1.27, 1.42]</td><td>1.11 [0.99, 1.23]</td><td>1.53 [1.37, 1.71]</td><td>SD</td></tr>
<tr><td>Observed Titer Intercept</td><td>0 [0, 0.01]</td><td>0.01 [0, 0.03]</td><td>0.01 [0, 0.04]</td><td>SD</td></tr>
</table>

**Table S8. Parameter estimates and their credible intervals for the time to undetectability.**

All entries display the median parameter estimate followed in brackets by the 90% credible interval. Parameters are stratified into columns by tissue groups (upper respiratory tract, URT; lower respiratory tract, LRT; GI, gastrointestinal tract), with the cofactor group indicated by the righthand row label (e.g., age) and the particular cofactor indicated by the lefthand row label (e.g., juvenile vs. adult vs. geriatric). Cells with two parameter estimates separated by a "|" are tissue-specific, as follows: for URT, nose | throat; for LRT, trachea | lung; for GI, upper GI | lower GI. All parameters are explained in the methods. The titer cofactor group accounts for the dependence of the time to undetectability on the peak titer. For all lab effects, see Figure S12.

| Cofactor | URT | LRT | GI | |
|---|---|---|---|---|
| | 1.4 [1.3, 1.5] | 1.5 [1.35, 1.68] | 1.09 [1.01, 1.27] | Shape |
| | -3.8 [-7.58, -0.06] | -10.82 [-15.58, -5.8] | -2.37 [-7.66, 3.09] | Intercept |
| | -0.22 [-1.73, 1.3] | 0.57 [-1.03, 2.18] | 0.28 [-1.39, 1.92] | Location |
| IN | 0.18 [-1.28, 1.67] \| -0.55 [-2.06, 0.92] | -0.22 [-1.81, 1.35] \| -0.3 [-1.84, 1.26] | 0 [-1.65, 1.67] \| 0.33 [-1.25, 1.88] | Route |
| IT | 0.97 [-0.57, 2.52] \| -1.12 [-2.63, 0.43] | -0.33 [-1.93, 1.2] \| -0.16 [-1.73, 1.44] | 0 [-1.66, 1.64] \| 0.43 [-1.14, 1.98] | Route |
| IN+IT | -0.25 [-1.68, 1.14] \| 0.61 [-0.9, 2.14] | 0.03 [-1.58, 1.66] \| 1.23 [-0.26, 2.7] | 0 [-1.62, 1.63] \| 0 [-1.52, 1.58] | Route |
| AE | -0.51 [-1.95, 0.94] \| 0.87 [-0.63, 2.4] | 0.01 [-1.64, 1.63] \| -0.24 [-1.85, 1.38] | -0.02 [-1.64, 1.63] \| -0.58 [-2.19, 1.01] | Route |
| IG | 0.05 [-1.59, 1.7] \| 0.02 [-1.67, 1.68] | -0.01 [-1.63, 1.66] \| 0 [-1.65, 1.63] | -0.02 [-1.61, 1.62] \| 0.06 [-1.55, 1.67] | Route |
| Nose | -0.08 [-4.7, 4.33] \| -0.04 [-4.34, 4.24] | 0.23 [-4.51, 4.94] \| 2.36 [-2.11, 6.76] | 0.01 [-5.78, 5.66] \| -0.48 [-5.12, 4.07] | Dose |
| Throat | 0.37 [-4.04, 4.91] \| 0.45 [-3.89, 4.94] | 0.52 [-4.22, 5.15] \| 3.25 [-1.21, 7.92] | 0.03 [-5.86, 5.86] \| -0.24 [-4.83, 4.22] | Dose |
| Trachea | -0.67 [-3.39, 2.16] \| 1.73 [-1.15, 4.61] | -0.13 [-4.53, 4.34] \| -3.42 [-6.87, 0.07] | 0 [-5.67, 5.64] \| -1.43 [-5.07, 2.29] | Dose |
| Lung | 4.24 [0.31, 8.04] \| 3.25 [-0.89, 7.41] | -0.08 [-5.89, 5.8] \| 3.31 [-1.83, 8.29] | -0.03 [-5.72, 5.78] \| -2.11 [-6.92, 2.71] | Dose |
| Upper GI | 0.62 [-4.55, 5.58] \| 0.06 [-5.83, 5.67] | 0.03 [-5.67, 5.84] \| 0 [-5.8, 5.84] | 0.08 [-5.64, 5.77] \| 0.68 [-4.16, 5.59] | Dose |
| Juvenile | -0.1 [-1.35, 1.17] | -0.33 [-1.68, 1.02] | 0.77 [-0.76, 2.25] | Age |
| Adult | 0.68 [-0.58, 1.91] | 1.1 [-0.21, 2.43] | -0.63 [-2.15, 0.88] | Age |
| Geriatric | -0.37 [-1.95, 1.18] | -0.75 [-2.31, 0.8] | 0.17 [-1.42, 1.72] | Age |
| | -1.99 [-2.94, -1.02] | -2.95 [-4.12, -1.79] | -0.09 [-1.5, 1.35] | Sex |
| RM | 0.2 [-0.96, 1.39] | 0.55 [-0.73, 1.79] | 0.39 [-1.07, 1.85] | Species |
| CM | 0.59 [-0.63, 1.8] | -0.08 [-1.44, 1.25] | 0.32 [-1.14, 1.77] | Species |
| AGM | -0.58 [-1.89, 0.7] | -0.48 [-1.85, 0.95] | -0.43 [-1.96, 1.11] | Species |
| Total RNA | -1.42 [-2.65, -0.2] | -1.15 [-2.45, 0.12] | 0.28 [-1.12, 1.75] | Assay |
| gRNA | -1.32 [-2.52, -0.05] | -1.52 [-2.84, -0.16] | 0.21 [-1.26, 1.69] | Assay |
| sgRNA | 2.1 [0.89, 3.32] | 1.56 [0.25, 2.9] | -0.26 [-1.88, 1.35] | Assay |
| Culture | 0.88 [-0.5, 2.24] | 1.14 [-0.38, 2.6] | 0.03 [-1.57, 1.67] | Assay |
| Inoculated | -2.44 [-2.88, -2.01] | -1.98 [-2.81, -1.13] | 0 [-1.63, 1.65] | Titer |
| Not Inoculated | -1.85 [-2.45, -1.26] | -2.38 [-3.05, -1.68] | -2.04 [-2.85, -1.26] | Titer |

**Table S9. Median values and 90% prediction intervals for the probability of positivity.**

All entries display the median estimate followed in brackets by the 90% prediction interval. Estimates are stratified into tissues and detection assay as indicated by the labels on the righthand side. Estimates distinguish between doses indicated on the y axis and between routes as shown on the x axis. These estimates include samples for all demographic groups (i.e., they integrate across ages, sexes, and species), given their effects are generally small. These estimates are based on 1,000 samples from the model posteriors, where each such sample was used to generate an estimate for all cofactor combinations.

Probability of positivity

| Exposure Dose (pfu) | IN | IT | IN + IT | AE | IG | Tissue | Assay |
|---|---|---|---|---|---|---|---|
| $10^4$ | 94 [88, 97] | 69 [52, 82] | 90 [83, 95] | 93 [87, 97] | 81 [63, 93] | Nose | Total RNA |
| $10^7$ | 98 [95, 99] | 70 [53, 83] | 97 [93, 98] | 98 [96, 99] | 85 [67, 96] | Nose | Total RNA |
| $10^4$ | 92 [85, 96] | 90 [81, 95] | 90 [82, 95] | 96 [92, 99] | 83 [66, 93] | Throat | Total RNA |
| $10^7$ | 96 [92, 98] | 93 [86, 97] | 97 [94, 98] | 100 [98, 100] | 87 [69, 96] | Throat | Total RNA |
| $10^4$ | 81 [67, 91] | 84 [71, 93] | 77 [63, 88] | 85 [69, 94] | 80 [62, 93] | Trachea | Total RNA |
| $10^7$ | 84 [70, 92] | 86 [73, 94] | 83 [70, 91] | 91 [77, 98] | 84 [64, 96] | Trachea | Total RNA |
| $10^4$ | 81 [67, 90] | 89 [79, 95] | 91 [84, 96] | 90 [80, 96] | 83 [66, 93] | Lung | Total RNA |
| $10^7$ | 83 [71, 92] | 92 [83, 96] | 95 [90, 97] | 96 [89, 99] | 86 [69, 96] | Lung | Total RNA |
| $10^4$ | 70 [51, 86] | 55 [32, 78] | 58 [40, 79] | 80 [56, 95] | 81 [57, 94] | Upper GI | Total RNA |
| $10^7$ | 78 [62, 90] | 57 [34, 80] | 70 [53, 86] | 92 [71, 99] | 92 [70, 99] | Upper GI | Total RNA |
| $10^4$ | 76 [60, 89] | 70 [51, 86] | 67 [50, 84] | 66 [47, 83] | 73 [53, 88] | Lower GI | Total RNA |
| $10^7$ | 84 [69, 93] | 74 [55, 88] | 79 [65, 90] | 80 [63, 92] | 82 [62, 94] | Lower GI | Total RNA |
| $10^4$ | 73 [57, 85] | 29 [16, 46] | 63 [47, 76] | 71 [55, 83] | 44 [24, 69] | Nose | Culture |
| $10^7$ | 88 [78, 94] | 30 [17, 47] | 83 [72, 90] | 91 [80, 96] | 52 [27, 82] | Nose | Culture |
| $10^4$ | 68 [51, 82] | 63 [43, 79] | 63 [46, 77] | 83 [70, 92] | 47 [26, 71] | Throat | Culture |
| $10^7$ | 81 [66, 91] | 72 [52, 87] | 85 [73, 92] | 97 [92, 99] | 55 [28, 82] | Throat | Culture |
| $10^4$ | 51 [33, 71] | 56 [36, 75] | 45 [29, 64] | 57 [35, 80] | 50 [27, 74] | Trachea | Culture |
| $10^7$ | 56 [36, 75] | 60 [39, 78] | 53 [36, 71] | 71 [44, 93] | 55 [30, 84] | Trachea | Culture |
| $10^4$ | 50 [32, 69] | 66 [47, 82] | 72 [56, 85] | 69 [49, 84] | 54 [32, 76] | Lung | Culture |
| $10^7$ | 55 [36, 73] | 72 [54, 86] | 81 [68, 90] | 85 [66, 95] | 60 [35, 84] | Lung | Culture |
| $10^4$ | 33 [18, 56] | 21 [9, 43] | 23 [12, 44] | 46 [21, 79] | 48 [22, 78] | Upper GI | Culture |
| $10^7$ | 43 [25, 67] | 22 [9, 45] | 34 [18, 57] | 72 [34, 96] | 71 [33, 94] | Upper GI | Culture |
| $10^4$ | 41 [23, 64] | 34 [18, 58] | 30 [17, 52] | 29 [15, 52] | 36 [19, 62] | Lower GI | Culture |
| $10^7$ | 52 [32, 74] | 37 [20, 62] | 44 [27, 66] | 46 [26, 70] | 49 [25, 76] | Lower GI | Culture |

Exposure Route

**Table S10. Median values and 90% prediction intervals for the time to detectability.**

All entries display the median estimate followed in brackets by the 90% prediction interval. Estimates are stratified into tissues and detection assay as indicated by the labels on the righthand side. Estimates distinguish between doses indicated on the y axis and between routes as shown on the x axis. These estimates include samples for all demographic groups (i.e., they integrate across ages, sexes, and species), given their effects are generally small. These estimates are based on 1,000 samples from the model posteriors, where each such sample was used to generate an estimate for all cofactor combinations.

| Exposure Dose (pfu) | IN | IT | IN + IT | AE | IG | Tissue | Assay |
|---|---|---|---|---|---|---|---|
| | Time to detectability | | | | | | |
| $10^4$ | 0.51 [0.36, 0.85] | 0.89 [0.6, 1.48] | 0.54 [0.39, 0.86] | 0.59 [0.42, 0.96] | 0.88 [0.54, 1.56] | Nose | Total RNA |
| $10^7$ | 0.36 [0.25, 0.6] | 0.89 [0.58, 1.49] | 0.37 [0.27, 0.6] | 0.38 [0.24, 0.67] | 0.89 [0.48, 1.75] | Nose | Total RNA |
| $10^4$ | 0.63 [0.44, 1.02] | 0.63 [0.44, 1.03] | 0.55 [0.39, 0.89] | 0.58 [0.41, 0.96] | 0.85 [0.52, 1.48] | Throat | Total RNA |
| $10^7$ | 0.53 [0.36, 0.87] | 0.54 [0.36, 0.9] | 0.39 [0.28, 0.63] | 0.42 [0.26, 0.75] | 0.86 [0.46, 1.68] | Throat | Total RNA |
| $10^4$ | 0.42 [0.26, 0.71] | 0.61 [0.37, 1.05] | 0.38 [0.24, 0.6] | 0.4 [0.23, 0.72] | 0.67 [0.35, 1.3] | Trachea | Total RNA |
| $10^7$ | 0.3 [0.17, 0.55] | 0.58 [0.33, 1.04] | 0.25 [0.15, 0.44] | 0.27 [0.12, 0.62] | 0.69 [0.31, 1.6] | Trachea | Total RNA |
| $10^4$ | 0.79 [0.52, 1.24] | 0.46 [0.28, 0.79] | 0.46 [0.32, 0.7] | 0.45 [0.29, 0.71] | 0.64 [0.34, 1.17] | Lung | Total RNA |
| $10^7$ | 0.92 [0.58, 1.48] | 0.38 [0.22, 0.69] | 0.45 [0.3, 0.69] | 0.35 [0.2, 0.62] | 0.67 [0.3, 1.4] | Lung | Total RNA |
| $10^4$ | 0.94 [0.55, 1.59] | 1.44 [0.83, 2.55] | 0.9 [0.53, 1.51] | 0.93 [0.51, 1.7] | 1.38 [0.77, 2.46] | Upper GI | Total RNA |
| $10^7$ | 0.64 [0.32, 1.22] | 1.34 [0.69, 2.66] | 0.57 [0.28, 1.1] | 0.59 [0.24, 1.42] | 1.25 [0.61, 2.54] | Upper GI | Total RNA |
| $10^4$ | 1.99 [1.36, 3] | 1.41 [0.92, 2.25] | 1.54 [1.08, 2.3] | 2.66 [1.72, 4.13] | 1.14 [0.71, 1.9] | Lower GI | Total RNA |
| $10^7$ | 2.26 [1.53, 3.42] | 1.28 [0.82, 2.08] | 1.58 [1.12, 2.36] | 3.64 [1.98, 6.63] | 0.91 [0.51, 1.65] | Lower GI | Total RNA |
| $10^4$ | 0.55 [0.39, 0.91] | 0.96 [0.64, 1.59] | 0.58 [0.42, 0.93] | 0.63 [0.44, 1.04] | 0.95 [0.58, 1.67] | Nose | Culture |
| $10^7$ | 0.39 [0.26, 0.65] | 0.95 [0.62, 1.61] | 0.4 [0.29, 0.65] | 0.41 [0.25, 0.72] | 0.96 [0.51, 1.89] | Nose | Culture |
| $10^4$ | 0.68 [0.47, 1.1] | 0.68 [0.47, 1.11] | 0.59 [0.42, 0.96] | 0.63 [0.43, 1.03] | 0.91 [0.56, 1.61] | Throat | Culture |
| $10^7$ | 0.56 [0.38, 0.94] | 0.58 [0.39, 0.96] | 0.42 [0.3, 0.68] | 0.45 [0.28, 0.8] | 0.92 [0.5, 1.83] | Throat | Culture |
| $10^4$ | 0.43 [0.25, 0.74] | 0.62 [0.37, 1.09] | 0.38 [0.24, 0.63] | 0.41 [0.23, 0.75] | 0.68 [0.35, 1.35] | Trachea | Culture |
| $10^7$ | 0.31 [0.16, 0.57] | 0.59 [0.33, 1.09] | 0.26 [0.14, 0.45] | 0.28 [0.12, 0.64] | 0.71 [0.31, 1.63] | Trachea | Culture |
| $10^4$ | 0.81 [0.52, 1.28] | 0.47 [0.27, 0.82] | 0.47 [0.32, 0.73] | 0.46 [0.29, 0.74] | 0.65 [0.34, 1.21] | Lung | Culture |
| $10^7$ | 0.94 [0.59, 1.52] | 0.39 [0.22, 0.72] | 0.46 [0.3, 0.71] | 0.35 [0.2, 0.64] | 0.68 [0.3, 1.43] | Lung | Culture |
| $10^4$ | 1.24 [0.72, 2.11] | 1.89 [1.08, 3.38] | 1.19 [0.7, 2.01] | 1.23 [0.67, 2.27] | 1.81 [1.01, 3.26] | Upper GI | Culture |
| $10^7$ | 0.84 [0.42, 1.6] | 1.76 [0.9, 3.53] | 0.75 [0.37, 1.46] | 0.77 [0.31, 1.89] | 1.65 [0.8, 3.38] | Upper GI | Culture |
| $10^4$ | 2.63 [1.75, 4.04] | 1.86 [1.18, 3.03] | 2.04 [1.38, 3.1] | 3.51 [2.18, 5.68] | 1.51 [0.91, 2.53] | Lower GI | Culture |
| $10^7$ | 2.99 [1.94, 4.63] | 1.69 [1.05, 2.8] | 2.1 [1.39, 3.23] | 4.79 [2.49, 9.11] | 1.2 [0.66, 2.19] | Lower GI | Culture |

Exposure Route

**Table S11. Median values and 90% prediction intervals for the time to peak titer.**

All entries display the median estimate followed in brackets by the 90% prediction interval. Estimates are stratified into tissues and detection assay as indicated by the labels on the righthand side. Estimates distinguish between doses indicated on the y axis and between routes as shown on the x axis. These estimates include samples for all demographic groups (i.e., they integrate across ages, sexes, and species), given their effects are generally small. These estimates are based on 1,000 samples from the model posteriors, where each such sample was used to generate an estimate for all cofactor combinations.

Time to peak titer

| Tissue | Assay | Exposure Dose (pfu) | IN | IT | IN + IT | AE | IG |
|---|---|---|---|---|---|---|---|
| Nose | Total RNA | $10^4$ | 2.57 [2.05, 3.55] | 2.32 [1.84, 3.21] | 2.18 [1.79, 3] | 2.7 [2.15, 3.77] | 2.48 [1.83, 3.59] |
| Nose | Total RNA | $10^7$ | 2.52 [1.95, 3.5] | 2.21 [1.7, 3.09] | 1.95 [1.59, 2.66] | 2.5 [1.78, 3.69] | 2.43 [1.61, 3.81] |
| Throat | Total RNA | $10^4$ | 2.6 [2.08, 3.62] | 2.27 [1.82, 3.11] | 2.33 [1.89, 3.23] | 2.28 [1.78, 3.21] | 3.07 [2.3, 4.38] |
| Throat | Total RNA | $10^7$ | 2.54 [2, 3.56] | 2.09 [1.64, 2.88] | 2.09 [1.69, 2.87] | 1.98 [1.36, 3] | 3.5 [2.38, 5.39] |
| Trachea | Total RNA | $10^4$ | 3.08 [2.04, 4.62] | 4.85 [3.29, 7.07] | 2.77 [1.97, 3.9] | 3 [1.95, 4.72] | 5.07 [3.01, 8.76] |
| Trachea | Total RNA | $10^7$ | 2.09 [1.34, 3.27] | 4.83 [3.24, 7.14] | 1.89 [1.34, 2.72] | 2.08 [1.16, 3.93] | 5.13 [2.71, 10.33] |
| Lung | Total RNA | $10^4$ | 3.87 [2.65, 5.53] | 4.31 [2.68, 6.88] | 2.8 [1.98, 3.88] | 2.3 [1.55, 3.37] | 4.88 [2.85, 8.41] |
| Lung | Total RNA | $10^7$ | 3.24 [2.17, 4.7] | 3.87 [2.33, 6.5] | 2.01 [1.43, 2.77] | 1.34 [0.81, 2.12] | 4.99 [2.53, 9.9] |
| Lower GI | Total RNA | $10^4$ | 5.05 [3.82, 6.72] | 3.53 [2.58, 4.81] | 4.08 [3.14, 5.31] | 5.6 [4.1, 7.58] | 3.3 [2.4, 4.6] |
| Lower GI | Total RNA | $10^7$ | 5.94 [4.46, 7.9] | 3.25 [2.37, 4.47] | 4.41 [3.42, 5.71] | 7.11 [4.61, 10.91] | 2.93 [2.02, 4.31] |
| Nose | Culture | $10^4$ | 1.89 [1.51, 2.63] | 1.87 [1.48, 2.66] | 1.64 [1.34, 2.3] | 2 [1.6, 2.82] | 1.99 [1.45, 2.92] |
| Nose | Culture | $10^7$ | 1.79 [1.38, 2.5] | 1.8 [1.38, 2.59] | 1.42 [1.15, 1.97] | 1.79 [1.3, 2.62] | 1.96 [1.29, 3.12] |
| Throat | Culture | $10^4$ | 1.95 [1.56, 2.76] | 1.73 [1.4, 2.42] | 1.74 [1.41, 2.45] | 1.72 [1.35, 2.44] | 2.35 [1.75, 3.37] |
| Throat | Culture | $10^7$ | 1.88 [1.47, 2.66] | 1.58 [1.24, 2.2] | 1.52 [1.22, 2.11] | 1.47 [1.02, 2.21] | 2.64 [1.82, 4.04] |
| Trachea | Culture | $10^4$ | 2.66 [1.72, 4.1] | 4.18 [2.79, 6.24] | 2.4 [1.67, 3.45] | 2.6 [1.64, 4.15] | 4.37 [2.56, 7.79] |
| Trachea | Culture | $10^7$ | 1.81 [1.14, 2.88] | 4.14 [2.74, 6.25] | 1.63 [1.14, 2.38] | 1.8 [0.99, 3.41] | 4.45 [2.31, 9.1] |
| Lung | Culture | $10^4$ | 3.38 [2.3, 5] | 3.68 [2.26, 6.12] | 2.43 [1.71, 3.43] | 2.01 [1.36, 2.96] | 4.22 [2.43, 7.4] |
| Lung | Culture | $10^7$ | 2.88 [1.94, 4.27] | 3.31 [1.96, 5.69] | 1.77 [1.26, 2.45] | 1.19 [0.73, 1.86] | 4.32 [2.18, 8.54] |
| Lower GI | Culture | $10^4$ | 5.59 [4.14, 7.58] | 3.91 [2.81, 5.45] | 4.49 [3.37, 6] | 6.36 [4.47, 8.96] | 3.61 [2.57, 5.05] |
| Lower GI | Culture | $10^7$ | 6.55 [4.79, 8.88] | 3.6 [2.58, 5.06] | 4.83 [3.63, 6.43] | 8.17 [5.03, 13.12] | 3.18 [2.15, 4.66] |

Exposure Route

**Table S12. Median values and 90% prediction intervals for the peak titer.**

All entries display the median estimate followed in brackets by the 90% prediction interval. Estimates are stratified into tissues and detection assay as indicated by the labels on the righthand side. Estimates distinguish between doses indicated on the y axis and between routes as shown on the x axis. These estimates include samples for all demographic groups (i.e., they integrate across ages, sexes, and species), given their effects are generally small. These estimates are based on 1,000 samples from the model posteriors, where each such sample was used to generate an estimate for all cofactor combinations.

| Exposure Dose (pfu) | Peak titer | | | | | | |
|---|---|---|---|---|---|---|---|
| | IN | IT | IN + IT | AE | IG | Tissue | Assay |
| $10^4$ | 7.09 [6.33, 7.86] | 6.86 [6.09, 7.66] | 6.98 [6.25, 7.71] | 7.83 [7.07, 8.61] | 6.53 [5.53, 7.56] | Nose | Total RNA |
| $10^7$ | 7.76 [6.98, 8.55] | 6.95 [6.13, 7.79] | 7.8 [7.07, 8.52] | 9.09 [8.14, 10.03] | 6.25 [4.89, 7.67] | Nose | Total RNA |
| $10^4$ | 6.59 [5.81, 7.38] | 7.08 [6.33, 7.85] | 6.46 [5.72, 7.23] | 7.79 [7.04, 8.56] | 6.27 [5.24, 7.34] | Throat | Total RNA |
| $10^7$ | 6.92 [6.11, 7.73] | 7.08 [6.31, 7.88] | 6.84 [6.11, 7.58] | 9.36 [8.37, 10.37] | 5.86 [4.44, 7.32] | Throat | Total RNA |
| $10^4$ | 5.54 [4.63, 6.56] | 5.04 [3.87, 6.08] | 5.97 [5.12, 6.85] | 5.57 [3.81, 7.39] | 5.7 [3.69, 7.79] | Trachea | Total RNA |
| $10^7$ | 5.5 [4.39, 6.63] | 5.72 [4.57, 6.84] | 6.85 [6.12, 7.67] | 6.49 [3.55, 9.42] | 5.66 [2.53, 8.88] | Trachea | Total RNA |
| $10^4$ | 6.79 [6.05, 7.64] | 5.59 [4.51, 6.81] | 6.57 [5.99, 7.44] | 6.22 [5.49, 7.02] | 5.8 [3.89, 7.78] | Lung | Total RNA |
| $10^7$ | 7.62 [6.75, 8.5] | 5.49 [4.25, 6.81] | 7.24 [6.67, 8.11] | 7.53 [6.41, 8.66] | 5.85 [2.8, 8.83] | Lung | Total RNA |
| $10^4$ | 4.69 [3.79, 5.59] | 4.25 [3.31, 5.2] | 4.51 [3.64, 5.37] | 5.44 [4.49, 6.39] | 4.61 [3.63, 5.55] | Lower GI | Total RNA |
| $10^7$ | 5.37 [4.47, 6.26] | 4.5 [3.54, 5.46] | 5.41 [4.59, 6.21] | 6.78 [5.28, 8.28] | 5.21 [4.11, 6.29] | Lower GI | Total RNA |
| $10^4$ | 3.55 [2.78, 4.33] | 3.08 [2.27, 3.9] | 3.41 [2.65, 4.17] | 4.29 [3.52, 5.07] | 2.73 [1.73, 3.77] | Nose | Culture |
| $10^7$ | 4.23 [3.43, 5.03] | 3.17 [2.32, 4.04] | 4.23 [3.48, 4.97] | 5.54 [4.6, 6.49] | 2.46 [1.1, 3.87] | Nose | Culture |
| $10^4$ | 3.04 [2.25, 3.84] | 3.27 [2.51, 4.07] | 2.9 [2.14, 3.69] | 4.23 [3.47, 5] | 2.41 [1.34, 3.51] | Throat | Culture |
| $10^7$ | 3.37 [2.56, 4.2] | 3.28 [2.48, 4.11] | 3.28 [2.54, 4.03] | 5.79 [4.81, 6.78] | 1.95 [0.5, 3.47] | Throat | Culture |
| $10^4$ | 3.12 [2.12, 4.19] | 2.85 [1.67, 3.9] | 3.68 [2.81, 4.57] | 3.29 [1.54, 5.11] | 3.26 [1.23, 5.34] | Trachea | Culture |
| $10^7$ | 3.06 [1.87, 4.27] | 3.52 [2.35, 4.69] | 4.52 [3.73, 5.39] | 4.17 [1.21, 7.08] | 3.23 [0.03, 6.38] | Trachea | Culture |
| $10^4$ | 4.37 [3.53, 5.27] | 3.17 [2.04, 4.37] | 4.16 [3.47, 5.05] | 3.9 [3.1, 4.74] | 3.36 [1.44, 5.33] | Lung | Culture |
| $10^7$ | 5.18 [4.24, 6.13] | 3.06 [1.81, 4.36] | 4.83 [4.16, 5.7] | 5.16 [4.02, 6.32] | 3.4 [0.38, 6.37] | Lung | Culture |
| $10^4$ | 4.01 [3.04, 4.99] | 3.58 [2.58, 4.58] | 3.83 [2.9, 4.79] | 4.76 [3.7, 5.84] | 3.94 [2.88, 4.97] | Lower GI | Culture |
| $10^7$ | 4.69 [3.7, 5.68] | 3.83 [2.8, 4.85] | 4.74 [3.8, 5.66] | 6.1 [4.48, 7.71] | 4.55 [3.35, 5.71] | Lower GI | Culture |

Exposure Route

**Table S13. Median values and 90% prediction intervals for the time to undetectability.**

All entries display the median estimate followed in brackets by the 90% prediction interval. Estimates are stratified into tissues and detection assay as indicated by the labels on the righthand side. Estimates distinguish between doses indicated on the y axis and between routes as shown on the x axis. These estimates include samples for all demographic groups (i.e., they integrate across ages, sexes, and species), given their effects are generally small. These estimates are based on 1,000 samples from the model posteriors, where each such sample was used to generate an estimate for all cofactor combinations.

| Exposure Dose (pfu) | IN | IT | IN + IT | AE | IG | Tissue | Assay |
|---|---|---|---|---|---|---|---|
| | Time to undetectability | | | | | | |
| $10^4$ | 12.61 [10.19, 16.17] | 8.49 [6.57, 11.41] | 12.66 [10.51, 16.09] | 13.39 [10.86, 17.28] | 8.47 [6.11, 12.42] | Nose | Total RNA |
| $10^7$ | 14.23 [11.26, 18.68] | 8.65 [6.56, 11.82] | 14.96 [12.38, 18.9] | 14.8 [10.63, 21.43] | 8.08 [5.26, 13.51] | Nose | Total RNA |
| $10^4$ | 12.3 [9.93, 15.9] | 9.32 [7.37, 12.36] | 9.99 [8.25, 12.68] | 11.06 [8.87, 14.39] | 9.14 [6.46, 13.86] | Throat | Total RNA |
| $10^7$ | 12.93 [10.09, 17.05] | 8.71 [6.81, 11.64] | 9.82 [8.14, 12.34] | 12.31 [8.5, 18.35] | 9.25 [5.92, 15.86] | Throat | Total RNA |
| $10^4$ | 17.19 [11.56, 27.1] | 15.79 [11.35, 23.18] | 14.92 [10.67, 21.83] | 14.52 [8.96, 25.1] | 20.1 [11.57, 37.69] | Trachea | Total RNA |
| $10^7$ | 15.54 [9.89, 25.76] | 17.48 [12.19, 26.21] | 15.95 [11.08, 23.83] | 15.64 [7.34, 37.61] | 20.38 [9.75, 48.6] | Trachea | Total RNA |
| $10^4$ | 17.75 [12.34, 27.21] | 21.1 [14.32, 33.23] | 16.65 [11.96, 24.85] | 11.82 [7.89, 18.7] | 19.58 [11.22, 36.2] | Lung | Total RNA |
| $10^7$ | 16.41 [11.15, 25.65] | 22.66 [14.84, 36.97] | 16.65 [11.7, 24.94] | 10.77 [6.09, 20.82] | 19.95 [9.51, 48.12] | Lung | Total RNA |
| $10^4$ | 8.24 [6.31, 10.83] | 6.47 [4.84, 8.7] | 7.41 [5.78, 9.57] | 10.36 [7.8, 13.83] | 6.24 [4.6, 8.67] | Lower GI | Total RNA |
| $10^7$ | 9.72 [7.36, 13.03] | 6.54 [4.8, 8.99] | 8.8 [6.86, 11.4] | 14.7 [9.91, 22.84] | 6.2 [4.31, 9.31] | Lower GI | Total RNA |
| $10^4$ | 5.26 [4.36, 6.75] | 4.33 [3.48, 5.67] | 5.17 [4.32, 6.56] | 5.62 [4.65, 7.19] | 4.36 [3.3, 6.1] | Nose | Culture |
| $10^7$ | 5.74 [4.63, 7.44] | 4.37 [3.43, 5.8] | 5.78 [4.79, 7.33] | 5.95 [4.4, 8.34] | 4.23 [2.9, 6.58] | Nose | Culture |
| $10^4$ | 5.22 [4.31, 6.73] | 4.53 [3.68, 5.86] | 4.32 [3.63, 5.51] | 4.67 [3.88, 5.97] | 4.74 [3.53, 6.69] | Throat | Culture |
| $10^7$ | 5.37 [4.32, 6.95] | 4.2 [3.33, 5.53] | 4.11 [3.45, 5.18] | 4.92 [3.63, 6.99] | 4.91 [3.34, 7.7] | Throat | Culture |
| $10^4$ | 9.04 [6.2, 13.73] | 9.93 [7.29, 14.03] | 8.6 [6.24, 12.24] | 8.53 [5.47, 14.07] | 11.35 [6.86, 19.62] | Trachea | Culture |
| $10^7$ | 7.89 [5.12, 12.75] | 10.76 [7.67, 15.78] | 8.75 [6.02, 13.18] | 8.75 [4.37, 20.15] | 11.67 [6.01, 24.6] | Trachea | Culture |
| $10^4$ | 9.67 [6.88, 14.15] | 11.32 [7.87, 17.11] | 8.66 [6.29, 12.52] | 6.85 [4.77, 10.19] | 10.96 [6.68, 18.63] | Lung | Culture |
| $10^7$ | 8.84 [6.17, 13.22] | 11.84 [7.95, 18.52] | 8.35 [5.89, 12.26] | 5.95 [3.54, 10.6] | 11.3 [5.85, 23.82] | Lung | Culture |
| $10^4$ | 8.44 [6.35, 11.25] | 6.56 [4.84, 8.89] | 7.47 [5.71, 9.82] | 10.62 [7.79, 14.5] | 6.26 [4.52, 8.74] | Lower GI | Culture |
| $10^7$ | 9.93 [7.41, 13.41] | 6.55 [4.77, 9.05] | 8.76 [6.68, 11.65] | 15.05 [9.95, 23.33] | 6.13 [4.17, 9.08] | Lower GI | Culture |

Exposure Route

**Table S14. Median values and 90% prediction intervals for the duration of detectable infection.** All entries display the median estimate followed in brackets by the 90% prediction interval. Estimates are stratified into tissues and detection assay as indicated by the labels on the righthand side. Estimates distinguish between doses indicated on the y axis and between routes as shown on the x axis. These estimates include samples for all demographic groups (i.e., they integrate across ages, sexes, and species), given their effects are generally small. These estimates are based on 1,000 samples from the model posteriors, where each such sample was used to generate an estimate for all cofactor combinations.

| Exposure Dose (pfu) | IN | IT | IN + IT | AE | IG | Tissue | Assay |
|---|---|---|---|---|---|---|---|
| | Duration | | | | | | |
| $10^4$ | 12.08 [9.68, 15.52] | 7.56 [5.68, 10.34] | 12.11 [9.97, 15.4] | 12.77 [10.27, 16.56] | 7.52 [5.2, 11.4] | Nose | Total RNA |
| $10^7$ | 13.85 [10.89, 18.27] | 7.71 [5.67, 10.75] | 14.57 [12.01, 18.44] | 14.39 [10.25, 21.02] | 7.05 [4.34, 12.49] | | |
| $10^4$ | 11.66 [9.29, 15.12] | 8.66 [6.74, 11.62] | 9.43 [7.7, 11.98] | 10.45 [8.29, 13.68] | 8.24 [5.64, 12.85] | Throat | Total RNA |
| $10^7$ | 12.38 [9.55, 16.41] | 8.15 [6.26, 11] | 9.42 [7.76, 11.86] | 11.88 [8.06, 17.86] | 8.33 [5.09, 14.74] | | |
| $10^4$ | 16.74 [11.14, 26.62] | 15.14 [10.74, 22.52] | 14.52 [10.29, 21.4] | 14.1 [8.55, 24.64] | 19.37 [10.93, 36.9] | Trachea | Total RNA |
| $10^7$ | 15.23 [9.58, 25.38] | 16.85 [11.59, 25.59] | 15.68 [10.81, 23.53] | 15.32 [7.06, 37.32] | 19.59 [9.04, 47.75] | | |
| $10^4$ | 16.93 [11.59, 26.27] | 20.62 [13.88, 32.72] | 16.18 [11.48, 24.31] | 11.35 [7.43, 18.2] | 18.9 [10.61, 35.44] | Lung | Total RNA |
| $10^7$ | 15.43 [10.28, 24.58] | 22.26 [14.43, 36.55] | 16.19 [11.27, 24.4] | 10.41 [5.73, 20.41] | 19.17 [8.8, 47.35] | | |
| $10^4$ | 6.18 [4.6, 8.4] | 5 [3.61, 6.91] | 5.82 [4.45, 7.66] | 7.61 [5.54, 10.53] | 5.06 [3.58, 7.23] | Lower GI | Total RNA |
| $10^7$ | 7.38 [5.37, 10.31] | 5.21 [3.69, 7.36] | 7.17 [5.47, 9.49] | 10.82 [6.78, 18.26] | 5.24 [3.46, 8.2] | | |
| $10^4$ | 4.7 [3.82, 6.03] | 3.34 [2.57, 4.44] | 4.57 [3.75, 5.82] | 4.96 [4.03, 6.37] | 3.36 [2.38, 4.93] | Nose | Culture |
| $10^7$ | 5.33 [4.24, 6.96] | 3.38 [2.53, 4.6] | 5.36 [4.39, 6.83] | 5.51 [4.01, 7.85] | 3.18 [2.01, 5.38] | | |
| $10^4$ | 4.53 [3.65, 5.84] | 3.82 [3.02, 5.02] | 3.72 [3.06, 4.73] | 4.02 [3.29, 5.16] | 3.79 [2.68, 5.57] | Throat | Culture |
| $10^7$ | 4.79 [3.76, 6.24] | 3.6 [2.77, 4.83] | 3.69 [3.04, 4.65] | 4.44 [3.18, 6.46] | 3.92 [2.5, 6.48] | | |
| $10^4$ | 8.61 [5.8, 13.22] | 9.27 [6.68, 13.29] | 8.2 [5.85, 11.83] | 8.09 [5.07, 13.57] | 10.59 [6.19, 18.72] | Trachea | Culture |
| $10^7$ | 7.56 [4.83, 12.41] | 10.14 [7.08, 15.11] | 8.47 [5.75, 12.9] | 8.44 [4.08, 19.84] | 10.85 [5.27, 23.69] | | |
| $10^4$ | 8.84 [6.12, 13.23] | 10.83 [7.4, 16.56] | 8.17 [5.85, 11.99] | 6.38 [4.33, 9.67] | 10.26 [6.06, 17.85] | Lung | Culture |
| $10^7$ | 7.88 [5.3, 12.11] | 11.43 [7.55, 18.09] | 7.88 [5.45, 11.76] | 5.58 [3.18, 10.19] | 10.54 [5.19, 22.92] | | |
| $10^4$ | 5.72 [4.13, 7.97] | 4.62 [3.24, 6.48] | 5.39 [3.96, 7.29] | 6.97 [4.95, 9.93] | 4.67 [3.2, 6.85] | Lower GI | Culture |
| $10^7$ | 6.84 [4.84, 9.78] | 4.79 [3.3, 6.84] | 6.6 [4.88, 9.01] | 9.91 [6.07, 16.78] | 4.84 [3.11, 7.59] | | |

Exposure Route

**Table S15. Median values and 90% prediction intervals for infection AUC.**

All entries display the median estimate followed in brackets by the 90% prediction interval. Estimates are stratified into tissues and detection assay as indicated by the labels on the righthand side. Estimates distinguish between doses indicated on the y axis and between routes as shown on the x axis. These estimates include samples for all demographic groups (i.e., they integrate across ages, sexes, and species), given their effects are generally small. These estimates are based on 1,000 samples from the model posteriors, where each such sample was used to generate an estimate for all cofactor combinations.

| Exposure Dose (pfu) | AUC | | | | | | |
|---|---|---|---|---|---|---|---|
| | **IN** | **IT** | **IN + IT** | **AE** | **IG** | | |
| $10^4$ | 42.58 [33.35, 56.6] | 25.83 [19.09, 36.51] | 41.99 [33.55, 55.21] | 49.72 [39.31, 66.15] | 24.54 [15.98, 39.47] | **Nose** | **Total RNA** |
| $10^7$ | 53.52 [41.12, 72.68] | 26.68 [19.12, 38.56] | 56.57 [45.5, 73.4] | 65.18 [45.28, 98.1] | 21.85 [12.06, 43.35] | **Nose** | **Total RNA** |
| $10^4$ | 38.2 [29.49, 51.38] | 30.52 [23.44, 42.14] | 30.32 [24.02, 39.97] | 40.5 [31.42, 54.77] | 25.63 [16.6, 42.6] | **Throat** | **Total RNA** |
| $10^7$ | 42.58 [31.86, 58.48] | 28.73 [21.75, 39.9] | 32.04 [25.76, 41.58] | 55.41 [36.51, 86.78] | 24.26 [13.21, 48] | **Throat** | **Total RNA** |
| $10^4$ | 46.04 [28.28, 81.65] | 37.67 [23.87, 62.34] | 43.09 [28.61, 68.51] | 38.97 [18.03, 82.85] | 54.45 [22.95, 132.03] | **Trachea** | **Total RNA** |
| $10^7$ | 41.4 [23.6, 77.43] | 47.84 [30.09, 79.15] | 53.63 [35.35, 84.92] | 49.18 [14.31, 158] | 54.43 [13.65, 193.19] | **Trachea** | **Total RNA** |
| $10^4$ | 57.3 [37.18, 96.33] | 57.28 [34.5, 103.63] | 53.31 [36.07, 87.31] | 35.17 [22.13, 59.25] | 54.02 [23, 128.33] | **Lung** | **Total RNA** |
| $10^7$ | 58.71 [36.95, 99.43] | 60.71 [34.29, 114.43] | 58.99 [39.24, 95.24] | 39.07 [20.3, 80.39] | 54.75 [14.29, 191.39] | **Lung** | **Total RNA** |
| $10^4$ | 14.41 [9.26, 22.34] | 10.61 [6.42, 16.93] | 13.12 [8.54, 19.73] | 20.69 [13.05, 31.96] | 11.6 [7.03, 18.86] | **Lower GI** | **Total RNA** |
| $10^7$ | 19.75 [12.73, 30.78] | 11.68 [7.02, 18.77] | 19.38 [13.15, 28.37] | 36.76 [19.34, 70.32] | 13.6 [7.82, 23.85] | **Lower GI** | **Total RNA** |
| $10^4$ | 8.26 [6.08, 11.54] | 5.11 [3.47, 7.5] | 7.74 [5.66, 10.78] | 10.56 [8.03, 14.46] | 4.53 [2.55, 7.97] | **Nose** | **Culture** |
| $10^7$ | 11.18 [8.28, 15.68] | 5.33 [3.48, 8] | 11.27 [8.57, 15.28] | 15.19 [10.45, 22.91] | 3.81 [1.43, 8.68] | **Nose** | **Culture** |
| $10^4$ | 6.82 [4.79, 9.88] | 6.2 [4.46, 8.87] | 5.36 [3.79, 7.63] | 8.46 [6.42, 11.61] | 4.48 [2.25, 8.17] | **Throat** | **Culture** |
| $10^7$ | 8 [5.57, 11.59] | 5.87 [4.08, 8.59] | 6 [4.43, 8.3] | 12.83 [8.63, 20.02] | 3.71 [0.87, 9] | **Throat** | **Culture** |
| $10^4$ | 13.16 [7.16, 25] | 12.9 [7.02, 22.38] | 14.89 [9.44, 24.54] | 13.17 [4.52, 30.65] | 16.87 [4.59, 45.49] | **Trachea** | **Culture** |
| $10^7$ | 11.37 [5.45, 23.32] | 17.61 [10.13, 30.44] | 19.05 [11.97, 31.61] | 17.21 [3.03, 61.64] | 16.8 [1.96, 67.83] | **Trachea** | **Culture** |
| $10^4$ | 19.05 [11.9, 32.47] | 16.84 [8.96, 32.38] | 16.92 [11.02, 28.52] | 12.34 [7.54, 20.81] | 16.93 [5.24, 42.38] | **Lung** | **Culture** |
| $10^7$ | 20.24 [12.26, 34.7] | 17.2 [8.25, 34.84] | 19.01 [12.19, 31.65] | 14.29 [7.18, 29.07] | 17.22 [1.94, 63.9] | **Lung** | **Culture** |
| $10^4$ | 11.42 [6.71, 18.76] | 8.24 [4.53, 13.85] | 10.31 [6.08, 16.54] | 16.58 [9.79, 27.02] | 9.15 [5.09, 15.88] | **Lower GI** | **Culture** |
| $10^7$ | 15.96 [9.62, 26.11] | 9.18 [5.06, 15.47] | 15.61 [9.82, 24.27] | 30.14 [14.93, 59.71] | 10.94 [5.82, 20.1] | **Lower GI** | **Culture** |

Exposure Route
